# Mitochondrial phosphagen kinases support the volatile power demands of motor nerve terminals

**DOI:** 10.1101/2022.12.21.521290

**Authors:** Karlis A. Justs, Sergio Sempertegui, Danielle V. Riboul, Carlos D. Oliva, Ryan J. Durbin, Sarah Crill, Chenchen Su, Robert B. Renden, Yaouen Fily, Gregory T. Macleod

## Abstract

Neural function relies on cellular energy supplies meeting the episodic demands of synaptic activity, but little is known about the extent to which power demands (energy demands per unit time) fluctuate, or the mechanisms that match supply with demand. Here, in individually-identified glutamatergic motor neuron terminals of *Drosophila* larvae, we leveraged prior macroscopic estimates of energy demand to generate profiles of power demand from one action potential to the next. These profiles show that signaling demands can exceed non-signaling demands 10-fold within milliseconds, and terminals with the greatest fluctuation (volatility) in power demand have the greatest mitochondrial volume and packing density. We elaborated on this quantitative approach to simulate adenosine triphosphate (ATP) levels during activity and drove ATP production as a function of the reciprocal of the energy state, but this canonical feedback mechanism appeared to be unable to prevent ATP depletion during locomotion. Muscle cells possess a phosphagen system to buffer ATP levels but phosphagen systems have not been described for motor nerve terminals. We examined these terminals for evidence of a phosphagen system and found the mitochondria to be heavily decorated with an arginine kinase, the key element of invertebrate phosphagen systems. Similarly, an examination of mouse cholinergic motor nerve terminals found mitochondrial creatine kinases, the vertebrate analogues of arginine kinases. Knock down of arginine kinase in *Drosophila* resulted in rapid depletion of presynaptic ATP during activity, indicating that, in motor nerve terminals, as in muscle, phosphagen systems play a critical role in matching power supply with demand.

**SIGNIFICANCE:** Failure of metabolic processes to supply neurons with energy at an adequate rate can lead to synaptic dysfunction and cell death under pathological conditions. Using a quantitative approach at fruit fly motor nerve terminals we generated the first temporal profiles of presynaptic power demand during locomotor activity. This approach revealed challenges for the known mechanisms that match cellular power supply to demand. However, we discovered that motor nerve terminals in fruit flies and mice alike are supported by phosphagen systems, more commonly seen in muscles where they store energy and buffer mismatch between power supply and demand. This study highlights an understudied aspect of neuronal bioenergetics which may represent a bulwark against the progression of some neuropathologies.

## INTRODUCTION

Synaptic energy requirements in neurons are highly variable over time, reflecting the intermittence of action potentials (APs) that trigger synaptic activity. The mechanisms stabilizing the concentration of cytosolic ATP ([ATP]) are of great interest as the free energy available to power cellular machinery drops rapidly even with small (∼5%) falls in [ATP], an effect modeled extensively in vertebrate heart tissues (Balaban, 2009). Secondary consequences can be expected through ATP’s role as an allosteric regulator of signaling pathways (Cable and Briggs, 1988; Clarke, 2009), and through ATP’s influence on protein solubility and stability as a hydrotrope (Patel et al., 2017). Far more evident are the consequences of large falls in [ATP], where neurons can no longer sustain basic functions such as maintaining Na^+^/K^+^ gradients and removing Ca^2+^ (Caldwell, 1960; Castro et al., 2006). Low [ATP] accompanies cell death and dysfunction in ischemic neural tissues when a scarcity of O_2_ limits ATP production via oxidative phosphorylation [Ox-Phos (Lipton and Whittingham, 1982)] and glycolysis is unable to fully compensate (Malthankar-Phatak et al., 2008). Low [ATP] also characterizes those neurodegenerative diseases associated with mitochondrial dysfunction (Johri and Beal, 2012; Vandoorne et al., 2018).

The first step towards elucidating the molecular identity of these mechanisms, and their relative importance in both physiology and pathology, might be to make empirical estimates of the variable of interest: cytosolic [ATP]. However, while a range of cytosolic [ATP] reporters have been developed (Koveal et al., 2020) they are not capable of delivering estimates of small changes in [ATP] on a time scale of hundreds of milliseconds, which is the time scale of AP bursts. Furthermore, the measure of [ATP] itself is net of supply and demand and additional layers of experimentation are required to elucidate mechanisms coordinating supply with demand. In light of these limitations, we began with a quantitative approach, and modeled presynaptic energy demands on a millisecond timescale in six different *Drosophila* larval motor neuron (MN) terminals (Justs et al., 2022). It allowed us to investigate whether ATP production based on feedback from the concentration of adenine nucleotides and inorganic phosphate (P_i_) could stabilize [ATP] in terminals that undergo large fluctuations (volatility) in power demand. Surprisingly, feedback to Ox-Phos based on the cellular energy state [represented as [ATP]/([ADP][P_i_])] was wholly inadequate to stabilize [ATP]. However, the discovery of an arginine kinase in these terminals alerted us to the possibility that, like muscle cells, a phosphagen system is integral to energy metabolism of MN terminals. Phosphagen systems utilize guanidine compounds to buffer ATP levels via a reversible reaction catalyzed by a phosphagen kinase (phosphagen + MgADP + H^+^ ⇌ guanidine acceptor + MgATP) (Wallimann et al., 1992). Once a phosphagen system was incorporated into our model, it predicted greater [ATP] stability and a reduction in the required acceleration rate of ATP production. Presynaptic imaging of cytosolic [ATP] along with mitochondrial and cytosolic pH corroborated these predictions, as knock down of ArgK1 lead to a steady state deficit in the mitochondrial energetic state (matrix acidification), and a greater draw down of cytosolic ATP levels and pH. These findings provide insight into characteristics of MN energy metabolism that might represent a critical, but overlooked, vulnerability following genetic or environmental insults associated with MN disease.

## RESULTS

This study is comprised of several parts: First, using a quantitative approach, we investigate the magnitude and volatility of the ATP consumption *rate* (power demand) of six MN terminals during physiological activity. This is done by drawing on previous macroscopic estimates of ATP demand at rest and in response to a single AP (Justs et al., 2022). Secondly, we simulate the impact of these ATP consumption rates on ATP levels when ATP production is responsive to changes in adenine nucleotide concentrations and inorganic phosphate (P_i_). Thirdly, we demonstrate the presence and subcellular distribution of mitochondrial-targeted phosphagen kinases, in *Drosophila* and mouse MN terminals. We then simulate the impact of a phosphagen system on ATP levels, ATP production rates and ATP production rate acceleration in *Drosophila* MN terminals. Lastly, we demonstrate that knock down of *Drosophila* ArgK1 compromises presynaptic energy metabolism and ATP availability.

### *Drosophila* Motor Neuron Terminals are Morphologically and Physiologically Distinct

Four glutamatergic MNs innervate 4 muscle fibers in a stereotypical pattern where two morphologically distinct terminals can be found on each muscle fiber; a terminal with big boutons [type-Ib, (big)] and a terminal with small boutons [type-Is, (small)]. Three different MNs supply a type-Ib terminal to muscle fibers 12, 13 and 6/7 (shared), while a single MN supplies type-Is terminals to all 4 muscle fibers (Figure 1A) (Hoang and Chiba, 2001). Functional differentiation has been demonstrated between terminal types (Kurdyak et al., 1994; Lnenicka and Keshishian, 2000; Chouhan et al., 2010; Lu et al., 2016; Newman et al., 2017; Aponte-Santiago et al., 2020; He et al., 2022), and between type-Ib terminals on different muscle fibers (Chouhan et al., 2012; Wang et al., 2021; Justs et al., 2022).

**Figure 1.**
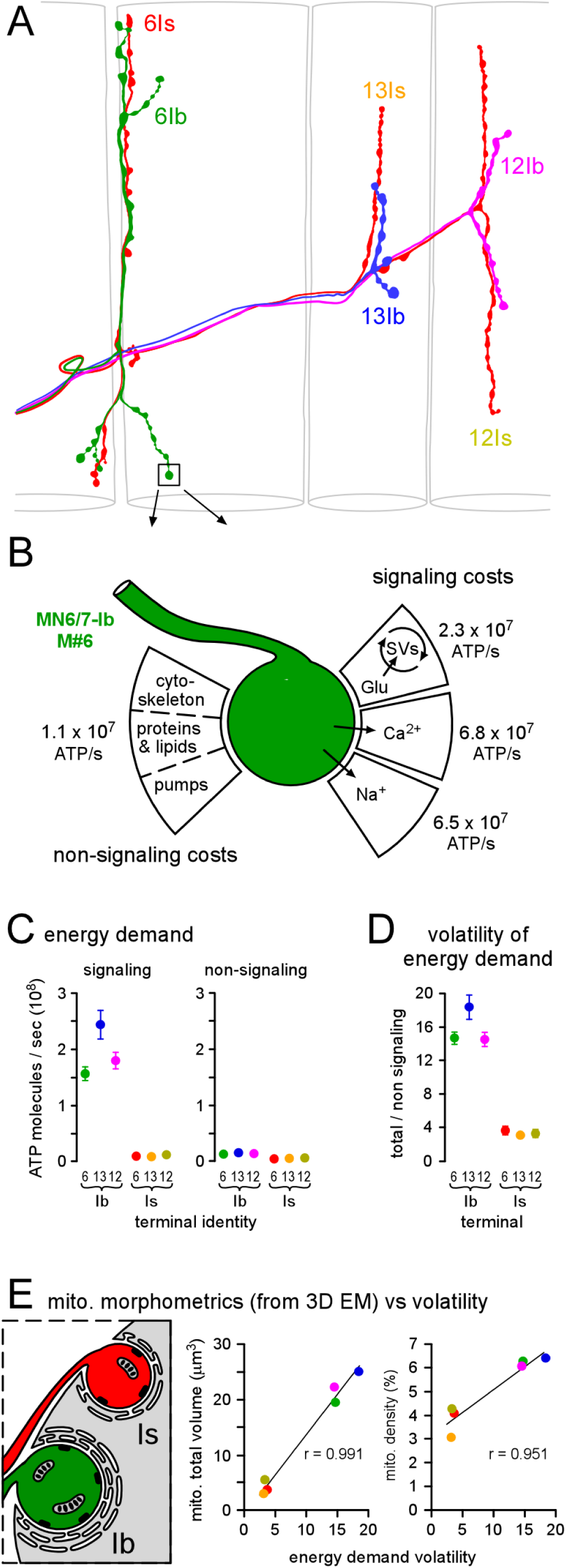
Motor Neuron (MN) Terminals Show Differences in Energy Demand Volatility. (A) A camera lucida interpretation of the stereotypical MN innervation of *Drosophila* larval body-wall muscle fibers in Figure S1. Four MNs (MN6/7-Ib: green; MN13-Ib: blue; MN12-Ib: pink; MNSNb/d-Is: red) innervate four body wall muscles (numbers 7, 6, 13 and 12) with six terminals. This color key is used throughout the manuscript. (B) A diagram representing aspects of energy expenditure previously calculated for each of the six different MN terminals (Justs et al., 2022), with values given for the terminal of MN6/7-Ib averaged over 1 second. Signaling energy demand (ATP/s) was estimated through direct measurements of neurotransmitter release, calcium ion (Ca^2+^) entry, and theoretical estimates of sodium ion (Na^+^) entry. Non-signaling energy demand, encompassing cytoskeleton turnover, lipid/protein synthesis and processes at the plasma membrane (e.g. Na^+^/K^+^ ATPase activity), was estimated by comparing the volume and surface area of each terminal with terminals in nervous tissue in which non-signaling OCR has been measured previously (Engl et al., 2017). (C) Plots of the energy demand during signaling and non-signaling. (D) A plot of the volatility of energy demand, calculated by dividing total energy demand (signaling + non-signaling) by non-signaling energy demand. Error bars for C and D (SEM) were calculated according to propagation of uncertainty theory. (E) Plots of Mitochondrial morphometric estimates versus energy demand volatility. Left: schematic of a single transmission electron micrograph through the NMJs of muscle 6. 3D reconstructions were made from such series from multiple larvae allowing morphometric analyses of presynaptic mitochondria (Justs et al., 2022). Center and right: total mitochondrial volume (* p < 0.0001) and density (* p < 0.005) plotted against energy demand volatility from D. Pearson Product Moment Correlation. Data reported in Tables S1 and S2.

### A Macroscopic (seconds time-scale) Measure of Energy Demand Volatility

In our previous study (Justs et al., 2022), we quantified the signaling presynaptic energy demands associated with a locomotory peristaltic cycle lasting 1 second, and the terminal of MN6/7-Ib is given as an example (Figure 1B). Here, we generated a measure of energy demand volatility by dividing the total energy demand by the non-signaling (housekeeping) energy demand (Figure 1C-1D; Table S1). This measure is best described as energy demand volatility, rather than power demand volatility, as there is a mismatch between the time of the signaling activity and the time of ATP consumption associated with this activity. Our analysis revealed energy demand volatility to be substantially higher for type-Ib terminals than type-Is. We also tested for correlations between previously quantified parameters of mitochondrial morphometry (Table S2) (Justs et al., 2022) and energy demand volatility (Figure 1E). Mammalian muscle fibers with a high peak power demand and range rely heavily on glycolysis (Schiaffino and Reggiani, 2011; Barclay, 2017), but here we found that MNs with the highest energy demand volatility had the highest mitochondrial volume and packing density (Figure 1E), supporting the notion that Ox-Phos can also support highly volatile energy demands in a presynaptic context.

### Computational Modeling Reveals Power Demand Volatility on a Millisecond Time Scale

As alluded to above, while an AP can invade a presynaptic terminal and trigger transmitter release within a millisecond, the ATP required to “reset” a presynaptic terminal after activity is consumed over a much longer period of time. Therefore, to gain an appreciation of the ways in which presynaptic energy demands fluctuate on a millisecond time scale, and how demands might summate during motor patterns, we developed a quantitative approach. In constructing a model capable of reporting instantaneous power demand (Figure 2) we assumed that the energy spent on each signaling process follows an exponential decay over time after each AP and that its integral (the area under the curve) is equivalent to the total amount of ATP consumed per AP (Figures 2A-2D shows values for MN6/7-Ib on muscle fiber 6). The time courses of the processes, and therefore the decay constants, differ by orders of magnitude. The time course for synaptic vesicle (SV) recycling and refilling reflects multiple component processes (Figure 2A), and the rationale for the weighted “blend” of time courses [13% @ 175 ms, 34% @ 7.21 s and 53% @ 20 s (Chanaday and Kavalali, 2018)] is provided in the Experimental Procedures. The time constants of the return of Ca^2+^ and Na^+^ to resting levels after an AP, via ATP-dependent pumps, are 38.9 ms (Justs et al., 2022) and 63.5 s (Mondragao et al., 2016), respectively (Figures 2B-2C). We assumed that the non-signaling power demand remained constant for each terminal, proportional to its surface area and volume (Figure 2B; also see Experimental Procedures). A plot of instantaneous presynaptic power demand can be generated for each peristaltic cycle by summing all processes above (Figure 2D). The slow decay of ATP consumption subsequent to activity is seen in the progressive climb in the cumulative ATP demand after a burst of APs (Figure 2E).

**Figure 2.**
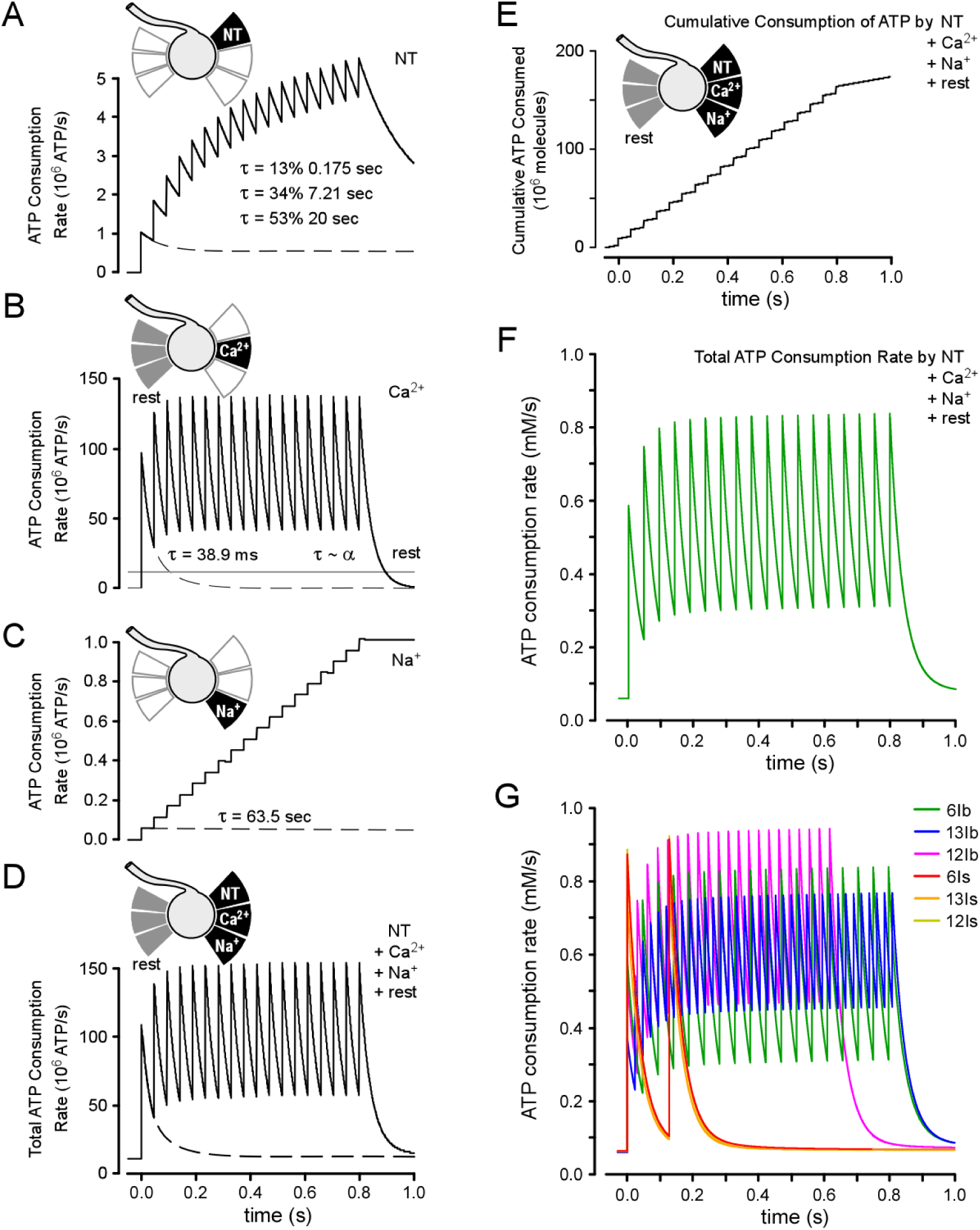
Computational Modeling Reveals Power Demand on a Millisecond Time Scale. (A) Plot of the ATP consumption rate for SV recycling/refilling for each action potential (AP) during a peristaltic cycle of 1 second. The decay subsequent to the first AP (dashed line) represents the summed decay of 3 different processes (modeled as a triple exponential). The integral of this decay is equivalent to the total amount of ATP consumed by SV recycling/refilling after the first AP. A cartoon, based on Figure 1B, is shown above plots A through E to indicate those aspects of energy expenditure plotted. Plots in A-F are for the terminal of MN6/7-Ib on muscle #6. (B) Plot of the ATP consumption rate for Ca^2+^ extrusion and non-signaling (rest) activities during a single peristaltic cycle. The dashed line represents decay after a single AP; τ = 38.9 ms. The ATP consumption rate for non-signaling activities was assumed to be constant throughout the peristaltic cycle (see Experimental Procedures). (C) Plot of the ATP consumption rate for Na^+^ extrusion during a peristaltic cycle. The dashed line represents decay after a single AP; τ= 63.5 seconds. (D) Plot of the total ATP consumption rate which includes each aspect of signaling (3 plots represented in A-C), along with the non-signaling rate (lower plot in B). (E) Plot of the cumulative amount of ATP consumed for signaling and non-signaling activities (shown in A-D). (F) Plot of the instantaneous ATP consumption rate normalized to terminal volume (unlike plots in A-E) through division of the number of ATP molecules by terminal volume. Note: the area under the curve represents only half the amount of ATP (0.45 mM) that will ultimately be needed (0.89 mM) to “reset” the terminal after 1 second of signaling activity on top of its non-signaling activity. (G) Plots of instantaneous ATP consumption rate for each terminal over a period of one peristaltic cycle (1 s).

### Instantaneous Power Demands Peak at Similar Levels in Different Terminals

To facilitate a comparison between terminals, the volume of each terminal was used (Justs et al., 2022) (Table S2) to convert the units of power (ATP molecules/s) to a concentration of ATP ([ATP]) consumed per second (mM ATP/s). Values for the MN6/7-Ib terminal on muscle fiber 6 are plotted in Figure 2F, where power demand transitions from a non-signaling rate of ∼0.06 mM ATP/s and spikes to ∼0.6 within milliseconds. Values for all 6 MNs terminals are plotted in Figure 2G. With each AP, ATP demand spikes to similar levels in type-Is and type-Ib terminals, but APs are less frequent in type-Is terminals and occur for a shorter proportion of each 1 s peristaltic cycle. The progressive summation of power demands from one burst of APs to the next becomes apparent when the plot is extended to 10 peristaltic cycles (Figures 3A), again, reflecting the “payoff period” of some activities (Na^+^ pumping and SV recycling and refilling) exceeding 1 second. The extended payoff period also results in estimates of power demand volatility (Table S3) that are only half those of energy demand volatility in our initial analysis (Figure 1D; Table S1).

**Figure 3.**
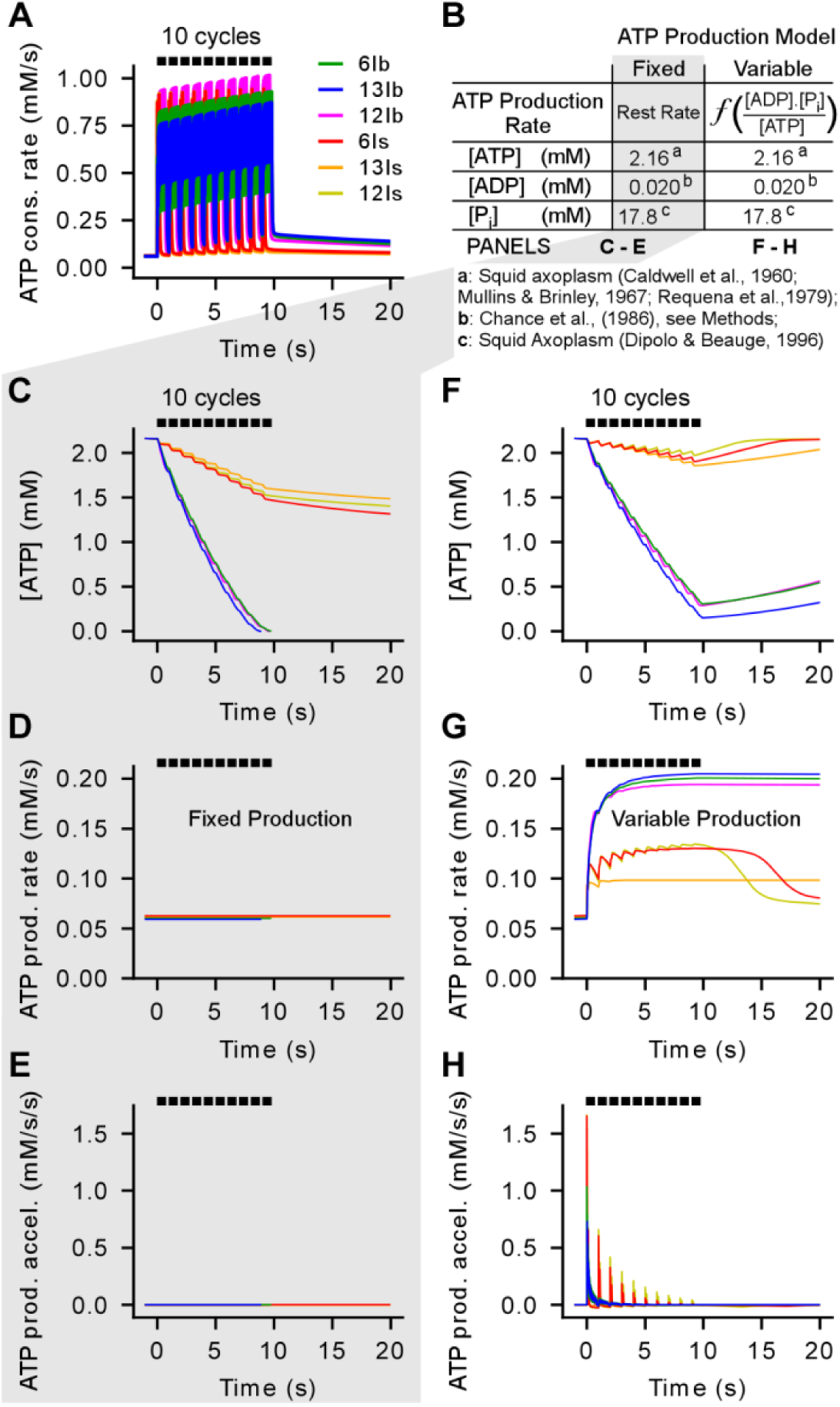
Terminals With High Power Demands Rapidly Draw Down ATP Levels. (A) Plots of the instantaneous the ATP consumption rate for each terminal over a period of 10 peristaltic cycles. (B) Two different models of ATP production; fixed: equivalent to the non-signaling power demand; variable: a function of the reciprocal of the energy state. (C) Plots of cytosolic ATP concentration ([ATP]) in each terminal, if ATP production does not increase. (D) Plots of ATP production rate. (E) Plots of rate of ATP production acceleration. (F) Plots of [ATP] in each terminal if the ATP production rate increases; capped at 0.032 mM ATP/s/1% mitochondrial volume density. (G) As in D. (H) As in E.

### Terminals with High Power Demands Will Rapidly Draw Down ATP Levels

We elaborated on our quantitative approach to investigate net [ATP] under assumptions of either a fixed or variable ATP production rate (Figure 3B), and relying on the dynamics of [ATP], [ADP], [AMP] and [P_i_]. Adenylate kinase activity, essential for adenine homeostasis, was incorporated into all calculations. When the ATP production rate was *fixed* to match the non-signaling ATP production rate, [ATP] exhausted within only 8 peristaltic cycles in type-Ib terminals (Figure 3C). Figures 3D-3E confirm ATP production rates did not increase. In implementing a *variable* rate of ATP production, we adopted a model of adenine nucleotide homeostasis that centered on Ox-Phos, as these terminals are thought to rely on Ox Phos to produce most of their ATP (Justs et al., 2022). Homeostasis was achieved by driving ATP production as a function of the reciprocal of the energy state, where energy state is defined as [ATP]/([ADP].[P_i_]) (Wilson, 2017a). Specifically, the decrease in [ATP], along with the increase in [ADP] and [P_i_], drive ATP production rate. Theoretical ATP production maxima were calculated for each nerve terminal, according to its mitochondrial density, by drawing on experimental data where the maximum output from Ox-Phos and glycolysis have been determined for a given mitochondrial mass in a cellular context. Therefore, the rate of ATP production is asymptotic to theoretical maximal outputs of Ox-Phos and glycolysis combined (see Experimental Procedures). Here (Figure 3F-H), the maxima were determined according to the maximum output of myocytes in a Seahorse analyzer (Mookerjee et al., 2017). Surprisingly, even when ATP production is allowed to accelerate, [ATP] catastrophe in hard working type-Ib terminals is only narrowly averted (Figures 3F-3H). This prediction of [ATP] catastrophe is surprising as larvae have an established capacity for “fast” crawling (∼1 mm/s) for many minutes (Steinert et al., 2006), and alerted us to the likelihood that the model was missing one or more important elements of presynaptic energy metabolism.

### Arginine Kinase (ArgK1), a Phosphagen Kinase, is Located in *Drosophila* Larval MN Terminals

In cell types such as muscle, a phosphagen system is incorporated into models of energy metabolism (Wilson, 2017a), and, while creatine kinases play an acknowledged role in the central nervous system (Andres et al., 2008) and have been described within central MNs of humans (Lowe et al., 2013), there has been no description of a role for phosphagens in lower MN terminals at NMJs. The invertebrate analogue of CK is arginine kinase (ArgK) which uses arginine to buffer ATP levels. Although there is no description of a role for ArgK in the MN terminals of invertebrates, it follows from the measurements of phosphoarginine in the squid giant axon that ArgKs would be active in the presynaptic terminals (Caldwell, 1960). One ArgK gene has been identified in *Drosophila* (CG4929) (from here on called Argk1) and two others are predicted (CG5144 & CG4546) (from here on called ArgK2 and ArgK3; flybase.org). Microarray data available from the FlyAtlas Transcriptome Project revealed that mRNA for ArgK1 are highly expressed in the larval nervous system while mRNA for ArgK2 and ArgK3 is expressed at either very low levels or not at all in the nervous system.

Here, the availability of a protein (GFP) trap line (stock # 51522, BDSC) (Buszczak et al., 2007) allowed us to image the distribution of a subset of ArgK1 isoforms *in situ*. Analysis of the insertion site (Figure 4B) indicates that 2 of 7 ArgK1 isoforms will be GFP-tagged near their N-terminus (ArgK1-RE and ArgK1-RA; Figure 4B-4C). Confocal microscopy of the ventral ganglion in ArgK1-GFP larvae revealed prominent GFP expression in the neuropil (Figure 4D), with punctate GFP in the periphery of MN somata - reminiscent of a mitochondrial distribution (Figure 4E). Unlike MN nuclei, muscle nuclei showed intense GFP expression (Figure 4F), while GFP in the rest of the muscle was relatively weak and displayed a sarcomeric pattern (Figure 4G), as observed via immunohistochemistry by (Lang et al., 1980) and consistent with a mitochondrial isozyme. Notably, the brightest ArgK1-GFP punctae colocalized with type-Ib and -Is MN terminals (Figures 4F-4I). Quantification of ArgK1-GFP fluorescence in the two terminal types revealed that not only was ArgK1-GFP most heavily expressed in the larger hard-working type-Ib terminals it was more heavily expressed per unit volume (Figures 4J-4K). Confirmation of ArgK1 specificity was achieved by demonstrating that RNAi directed against ArgK1 message resulted in near complete knock-down (KD) of the GFP signal (Figure 4L).

**Figure 4.**
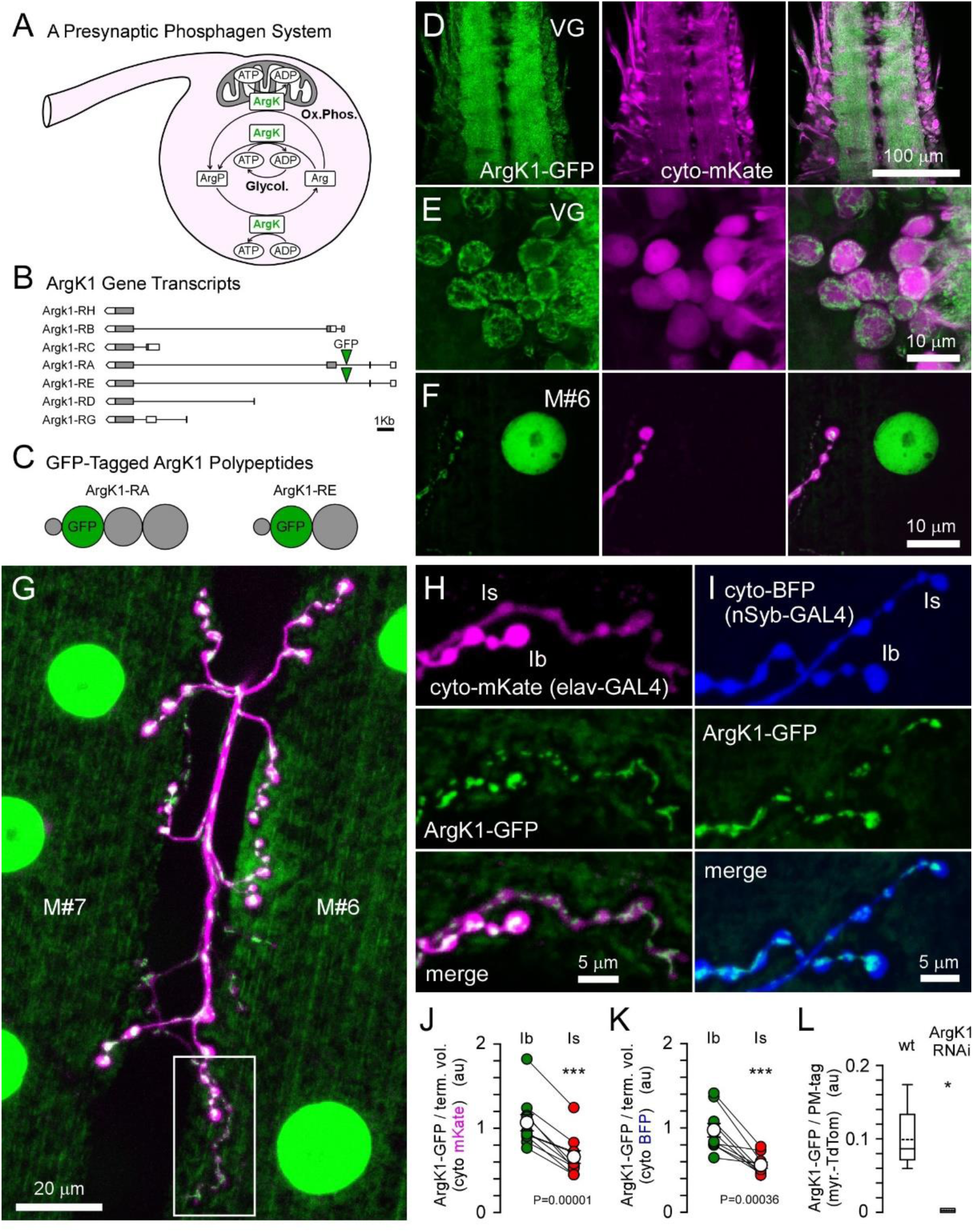
ArgK1 Is Heavily Expressed in *Drosophila* Motor Neurons (MNs). (A) A diagram representing the phosphagen system transferring phosphate groups from ATP to arginine, and back again, facilitated by arginine kinase (ArgK) interacting with other bioenergetics pathways. (B) Predicted splice isoforms of ArgK1 (FlyBase) showing intronic sites of transposon mediated GFP cDNA insertion via MiMIC cassettes. (C) A diagram representing the proteins generated from ArgK1 splice isoforms -RE and –RA, where the volume of each “sphere” represents the length of peptide corresponding to each exon. (D-L) Imaging and analysis performed on “live” dissected *Drosophila* larvae with MN terminals still connected to their somata. (D) A single confocal plane through the ventral ganglion (VG) showing the neuropil localization of ArgK1-GFP (one copy) relative to pan-neuronal (elav-GAL4) expression of mKate free in the cytosol. (E) A short confocal stack (4 μm axial) projection through a group of MN somata in the dorsal cortical region of the VG in D. (F) A short confocal stack projection of a nucleus near the surface of a body wall muscle fiber (#6) in the same larva as in D and E. Note the MN terminals. Same settings were used as in E allowing a comparison of ArgK1-GFP density in the respective nuclei. (G) A stack projection (12 μm) using the same settings as in E and F through terminals of MN6/7-Ib and MNSNb/d-Is in the cleft between muscle fibers #7 and #6 (same genotype as in D-F) (H) An image of terminals in the ROI in G, using the same settings as in E-F. (I) An image of a short confocal stack through terminals of MN6/7-Ib and MNSNb/d-Is on muscle fiber #6, showing ArgK1-GFP (one copy) relative to nSyb-GAL4 expression of TagBFP free in the cytosol. (J-K) Plots of the ArgK1-GFP fluorescence intensity relative to the volume of the terminal (see Experimental Procedures). ArgK1-GFP per unit volume is highest in type-Ib terminals (P<0.001 in both; Students’ T-test); N=10 terminal pairs in both J and K. (L) RNAi mediated knockdown (KD) of ArgK1-GFP using nSyb-GAL4 expression of ArgK1 dsRNA confirmed the identity of GFP-tagged proteins as isoforms of ArgK1 (P<0.001; Student’s T-test; N=5 each, type-Ib terminals).

### ArgK1 is Targeted to Presynaptic Mitochondria

To determine if GFP-tagged ArgK1 isoforms with extended N-termini are mitochondrion specific in MN terminals, we labeled mitochondria with tetramethyl ether [TMRE; (Chouhan et al., 2012)] (Figure 5A). We also saw co-localization of ArgK1-GFP with fluorescent mKate targeted to the mitochondrial matrix in MN somata (Figure 5B) and MN terminals (Figure 5C). The difference in the ArgK1-GFP signal across the two terminal types was similar to the difference in mitochondrial density across the terminals [type-Ib: 6.29±0.41%; type-Is: 4.10±0.39% (Justs et al., 2022)] (Figures 5D-5E). Finally, an increase in mitochondrial content has been observed in mouse skeletal muscle deficient in mitochondrial CK (van Deursen et al., 1993; Kaasik et al., 2003); however, knock down of ArgK1-GFP appeared to have no impact, relative to control, on mitochondrial-targeted GFP fluorescence in ratio to cytosolic TagBFP (Figures S5-5F).

**Figure 5.**
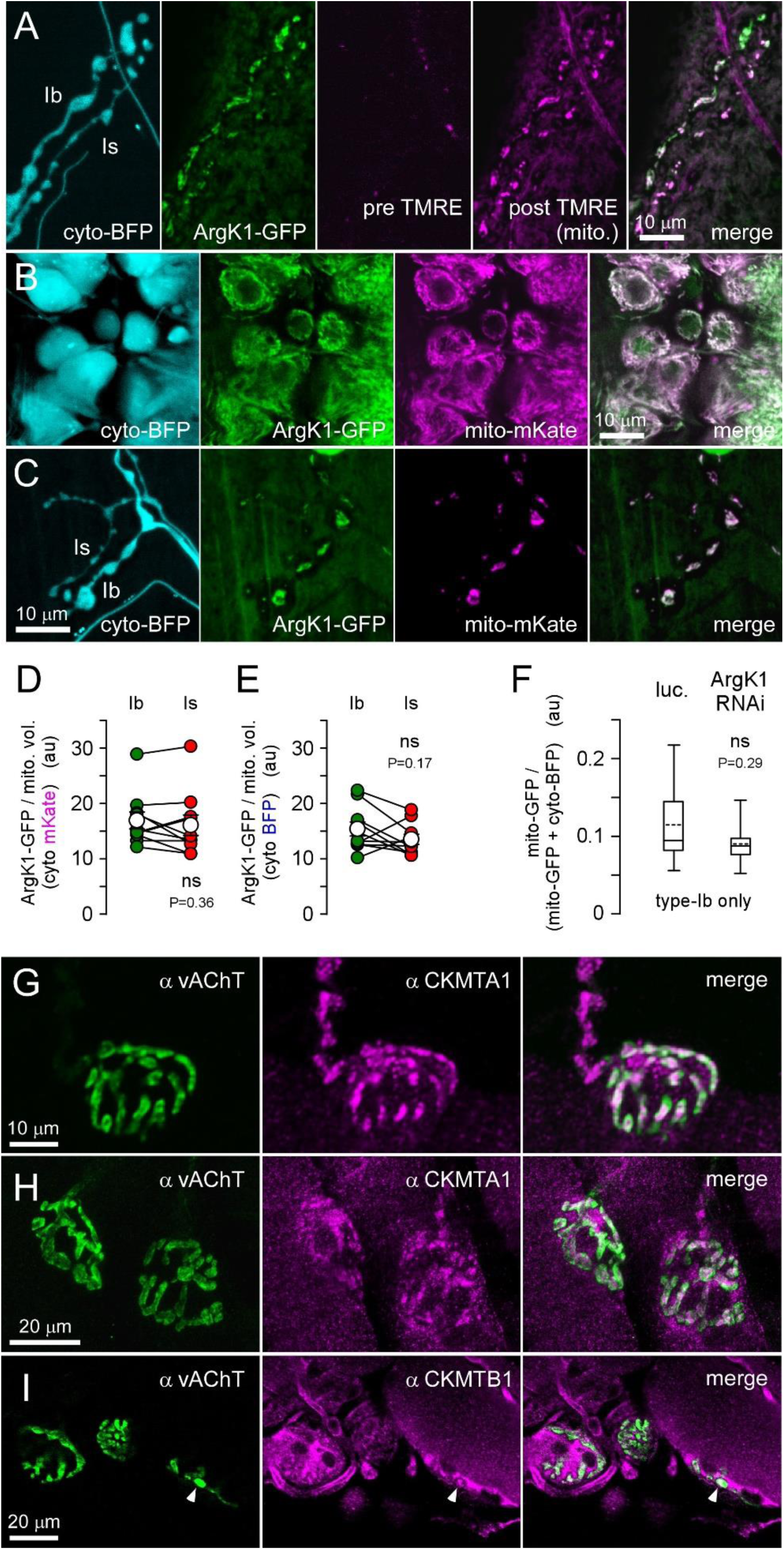
ArgK1 is Targeted to Mitochondria in MN Terminals. (A-F) Imaging and analysis performed on live dissected *Drosophila* larvae with MN terminals connected to their somata. (A) A confocal stack projection of terminals on muscle fiber #6 revealed by nSyb-GAL4 expression of cytosolic TagBFP (panel 1) and endogenous ArgK1-GFP (panel 2), absent of signal at wavelengths used to examine TMRE (panel 3; prior to TMRE exposure). A mitochondrial TMRE signal (panel 4) subsequent to exposure to 100 nM TMRE in 0.001% DMSO for 45 seconds (same imaging settings used in panel 3 and 4). (B) A confocal stack projection through a group of MN somata in the ventral ganglion demonstrating the relative localization of ArgK1-GFP (1 copy) and nSyb-GAL4 driven expression of mitochondrial-targeted mKate and cytosolic TagBFP. (C) A confocal stack projection through terminals on muscle fiber #6 (same larva as in B) (D-E) Plots of ArgK1-GFP fluorescence intensity relative to mitochondrial volume. Data in figure 4J-4K reanalyzed to reflect the proportion of terminal volume occupied by mitochondria in different terminal types (values in Figures 4J and 4K divided by 0.0629 for type-Ib terminals and 0.0410 for type-Is). D: N=10, P=0.36; E: N=10, P=0.17; Paired Student’s T-tests. (F) Box plots of mitochondrial content in type-Ib terminals expressing mitochondrial-targeted GFP and cytosolic BFP, along with either luciferase or ArgK1 dsRNA. Mitochondrial content was quantified by dividing GFP fluorescence by the sum of GFP and BFP fluorescence (images in Figure S5). Mean (dotted line) median (solid line) with 25-75% boxes and 10-90% whiskers [N=11 each, P=0.29; Student’s T-test]. (G-I) Imaging performed on neuromuscular junctions (NMJs) in the fixed mouse levator auris longus (LAL) muscle (G-H) Confocal stack projections through NMJs on single (G), or multiple (H), fiber(s) of the mouse LAL muscle. Synaptic vesicles within MN terminal branches are stained using the vesicular acetylcholine transporter (vAChT) antibody (green). CKMT1A is revealed in magenta using a rabbit polyclonal antibody. (I) An oblique optical section through NMJs on three fibers. Synaptic vesicles labeled with vAChT (green), CKMT1B labelled with a rabbit polyclonal antibody raised against human CKMT1B (magenta). Arrowhead points to CKMT1B signal in a cross-section through a presynaptic terminal.

### Mitochondrial Creatine Kinases are Found in Mouse MN Terminals

In mammals, a total of five nuclear genes code for CKs; two genes produce cytosolic-localized subunits which combine as dimers to form three different isozymes, and three genes produce mitochondrial-localized subunits which form tetramers of dimers. The cytosolic subunits are defined as either Brain or Muscle in their tissue distribution, while mitochondrial localized subunits are defined as either Ubiquitous or Sarcomeric (Schlattner et al., 2006). Here we probed the mouse levator auris longus muscle with antibodies for the vesicular acetylcholine transporter (vAChT) and the two ubiquitous mitochondrial creatine kinases (Figures 5G-5I). Clouds of synaptic vesicles defining the presynaptic compartments of the NMJs were revealed by the vAChT signal, while signal from both CKMT1A (Figure 5G-H) and CKMT1B (Figure 5I) revealed structures that were either central to, or interspersed with, the vAChT signal. The distribution of the CKMT1 signal suggested a mitochondrial localization, confirmed by staining motor terminals with a Tom20 antibody that labels the outer mitochondrial membrane (not shown).

### Phosphagen Systems will Stabilize ATP Levels, Occluding the Need for Rapid Changes in [ATP] Production

Having established the presynaptic presence of a phosphagen kinase, we simulated the impact of a phosphagen system on net [ATP] (Figure 6) as an extension of the energy metabolism model initiated above (Figures 2 and 3). We adopted values of 3.3 mM [Arg] and 7.5 mM [ArgP] from invertebrate neurons (Deffner and Hafter, 1959; Deffner, 1961; DiPolo and Beauge, 1996) (full set of parameters in Experimental Procedures), and limited our simulation to type-Ib and -Is terminals on muscle fiber 6, firing for 10 seconds (10 cycles) (Figures 6A-6B). The phosphagen system had a profound effect on ATP levels. When ATP production was driven as a function of the reciprocal of the energy state, [ATP] previously collapsed (Figures 3F-3H; Figures 6C-6E), but with an arginine-based phosphagen system, [ATP] fell by no more than 5% in hard-working type-Ib terminals (Figure 6F). This rescue of [ATP] will, in turn, have a profound impact on the free energy available from ATP hydrolysis. It is the ratio of [ATP] to the other adenine nucleotides (not shown here) that determines free energy availability (Ellington, 1989; Wallimann et al., 1992), and falls in [ATP] as small as 5%, can result in a fall in free energy of 15% (Figure S4D).

**Figure 6.**
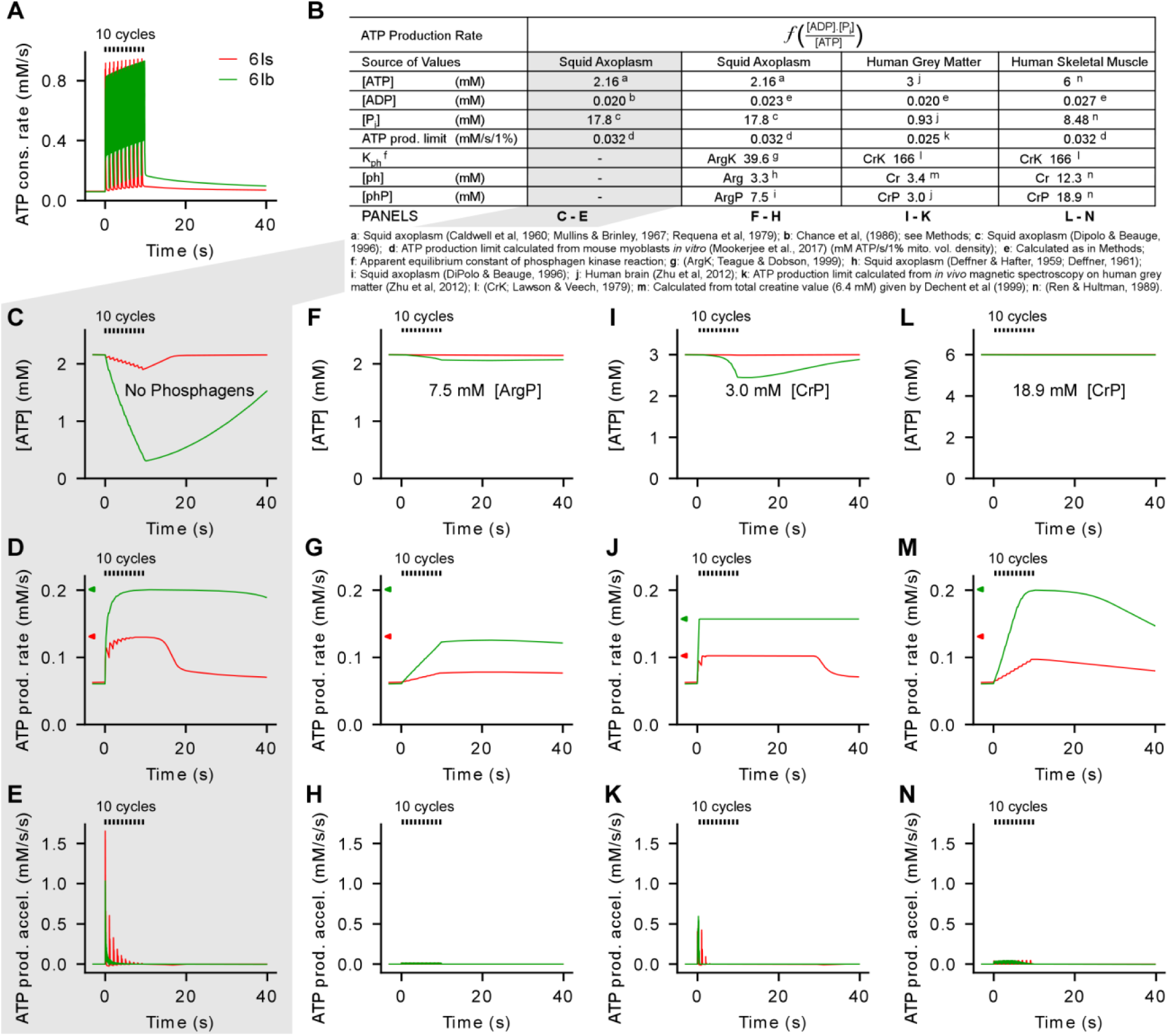
Phosphagen Systems Have the Capacity to Stabilize ATP Levels Without a Need for Rapid Changes in [ATP] Production. (A) Plots of instantaneous ATP consumption rates for terminals on muscle fiber #6 over a period of 10 peristaltic cycles. (B) Table showing parameter values from different animals and tissue types (columns 2-5) as the bases for simulations under the variable ATP production model. Parameters are introduced in column 1 and values are annotated with notes. ATP production limits are drawn from myocytes *in vitro*, not necessarily matching the animal and tissue type appearing at the top of the column. ATP production limits (mM ATP per second for each 1% mitochondrial volume density) are shown on the ordinates in D, G, J and M (green: type-Ib: 6.29%; red: type-Is: 4.10%) (C-E) Plots of the cytosolic ATP concentration (C), ATP production rate (D) and ATP production rate acceleration (E) corresponding to the activity in A, and in the absence of a phosphagen system (column 2 of B) (F-H) As in C-E, but with a phosphagen system, and using parameters from column 3 in B. ATP production limits as in D (I-K) As in C-E, but with a phosphagen system, and using parameters from column 4 in B. ATP production limits calculated from human cortex *in vivo*. (L-N) As in C-E, but with a phosphagen system, and using parameters from column 5 in B. ATP production limits as in D.

In contrast to the initial simulation with a variable ATP production rate (Figures 3F-3H and 6C-6E), when a phosphagen system was enabled ATP production rates did not closely approach their theoretical maxima (Figure 6F-6G). The same ATP demand must be satisfied in each scenario, and this could be reconciled by comparing the areas under the ATP production curves when plots were extended for a full 10 minutes (Figure S2). The most notable impact of the phosphagen system was the reduction of the required acceleration rate - by more than 100-fold (compare Figure 6H with 6E). This would seem to be a critical improvement, as, although glycolysis is capable of rapid acceleration (Walter et al., 1999), we are unaware of any studies indicating that mitochondrial respiration would be capable of increasing ATP levels in its accompanying cytosol at a rate of 1 mM ATP/s/s.

Parameters of cellular energy metabolism in the literature are highly variable, with estimates from muscle cells showing greater phosphagen reserves than neurons. Similarly, *in vivo* estimates of oxidative capacity yield substantially greater capacity than *in vitro* estimates (Tonkonogi and Sahlin, 1997). The capacity of *Drosophila* MN terminals to produce ATP is unknown, but the ATP *demand* during the 1^st^ second of activity averages 0.47 mM/s (Figure 2G) across hard working type-Ib terminals with an average mitochondrial volume density of 6.27% (Table S2); equivalent to 0.075 mM/s/1% mitochondrial mass. Our simulations indicated that the depth of the phosphagen reserve, rather than the combined capacity of glycolysis and Ox-Phos, had the greatest impact on [ATP] stability over short intervals (Figures 6I-6N and Figure S3). Whether the ATP production maximum of the terminals was limited to a nominal value of 0.032 mM/s/1% mitochondrial volume density (Figures 6F-6H; myocytes in *vitro*), 0.079 mM/s/1% (Figures S3F-S3H; *Drosophila* larval brains *ex vivo*), or 0.154 mM/s/1% (Figures S3I-S3K; *Drosophila* flight muscle *ex vivo*), it had little impact on net [ATP] when a phosphagen system was fully enabled.

### Knock Down of Arginine Kinase Compromises Presynaptic Energy Metabolism

To test the prediction that phosphagen kinases play a key role in presynaptic energy metabolism we examined the consequences of knocking down ArgK1 expression in *Drosophila* MNs. Various fluorescent reporters of cytosolic and mitochondrial energy metabolism were expressed in these terminals and the axons were stimulated for 10 seconds (Figure 7). Axons were driven at approximately twice their native firing frequencies to generate signals that overcome limitations of the fluorescent ATP reporters. PercevalHR is a reporter of the ATP to ADP ratio (Tantama et al., 2013), and the ATP to ADP ratio fell substantially more in ArgK1 KD terminals than in terminals where luciferase was expressed as a control (Figures 7A-7D). PercevalHR, like most ATP reporters, is extremely sensitive to the pH of its environment (Koveal et al., 2020), and so we examined activity-induced pH changes in these terminals using the pH reporter pHerry (Rossano et al., 2017). pHerry revealed that cytosolic pH fell substantially more in terminals in ArgK1 KD terminals than in controls (Figures 7E-7H). Subsequent pH-correction of the PercevalHR signals that extinguished nerve stimulus evoked changes in the control trace (Figure 7I), left an ostensibly pH-independent fall of 10% in the ArgK1 KD PercevalHR trace. Importantly, the inflexion in the ArgK1 KD PercevalHR trace (∼3 seconds after commencing axon stimulation) is seen in 15 of the 15 individual ArgK1 KD traces, but none of the 14 control traces (Figure S6).

**Figure 7.**
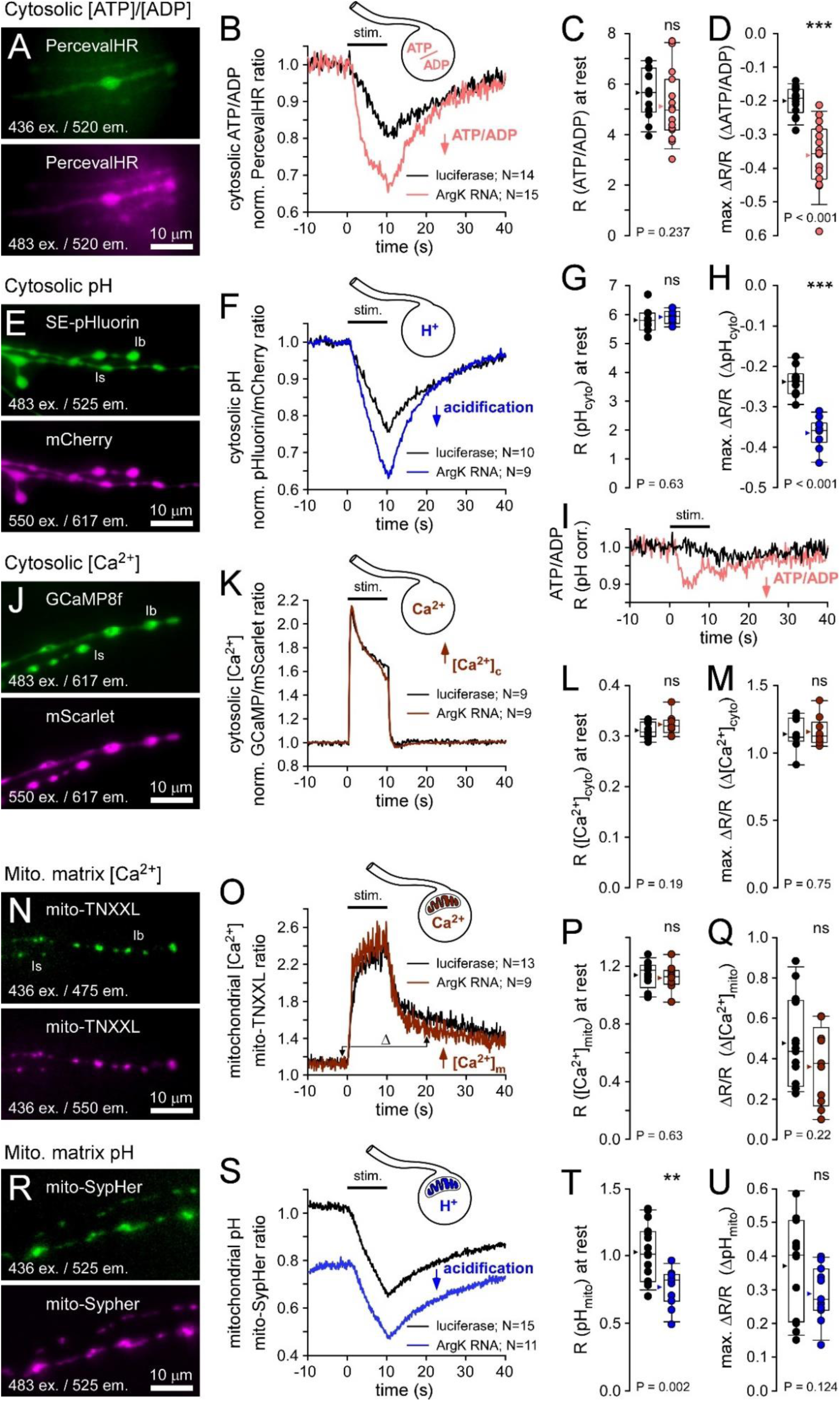
Live Imaging Reveals Deficiencies in Presynaptic Energy Metabolism after Arginine Kinase 1 (ArgK1) Knock Down. (A-M) Fluorescent reporters used in the cytosol of MN13-Ib terminals. (A) Motor neuron (MN) terminals expressing ATP/ADP reporter PercevalHR (nSyb:GAL4 > UAS-PercevalHR) in the 1 second period prior to nerve stimulation. Excitation and emission wavelengths are shown. Field diaphragm restricted to reduce background fluorescence. (B) Normalized PercevalHR ratio (483nm ex./436nm ex.) showing a decrease in ATP/ADP in response to 10 seconds of nerve stimulation in terminals expressing either ArgK1 dsRNA (N=14) or luciferase (N=15). Traces represent the average for each group. (C) Plots of the PercevalHR ratio prior to nerve stimulation (at rest). (D) Plots of the greatest change in the normalized PercevalHR ratio (max ΔR/R). (E) MN terminals expressing pH reporter pHerry (OK6:GAL4 > UAS-pHerry) prior to nerve stimulation. (F) Normalized pHerry ratio (SEpHluorin fluorescence relative to mCherry) showing acidification of the cytosol in response to nerve stimulation. (G) Plots of the pHerry ratio at rest. (H) Plots of the greatest change in the normalized pHerry ratio (max ΔR/R). (I) PercevalHR traces corrected for pH change. Both PercevalHR traces in B were corrected by a common factor (0.75) multiplied by their corresponding pHerry traces in F. The factor was chosen to extinguish any decrease in the control PercevalHR trace (luciferase) during nerve stimulation: Corrected PercevalHR ratio = PercevalHR ratio - [(pHerry ratio - 1) x 0.75], i.e. Panel I = Panel B – [(Panel F – 1) x 0.75]. (J) MN terminals expressing Ca^2+^ reporter Syt::mScarlet::GCaMP8f (OK6:GAL4 > UAS-Syt::mScarlet::GCaMP8f) prior to nerve stimulation. (K) Normalized GCaMP8f to mScarlet ratio showing an increase in [Ca^2+^]c in response to nerve stimulation. (L) Plots of the GCaMP8f to mScarlet ratio at rest. (M) Plots of the greatest change in the GCaMP8f to mScarlet ratio (max ΔR/R) (N-U) Fluorescent reporters used in the mitochondrial matrices of MN6-Ib terminals. (N) MN terminals expressing mitochondrial-targeted TN-XXL (nSyb:LexA > LexAop-mito-TN-XXL) prior to nerve stimulation. (O) The mito-TN-XXL ratio (Citrine/CFP) showing an increase in [Ca^2+^]_m_ in response to nerve stimulation. (P) Plots of the mito-TN-XXL ratio at rest. (Q) Plots of the change in the mito-TN-XXL ratio (ΔR/R), displacement measured between 1 second prior to stimulation and 10 seconds after stimulation cessation (avoids cytosolic TN-XXL signal artefact - see Experimental Procedures). (R) MN terminals expressing mitochondrial-targeted pH reporter SypHer (nSyb:LexA > LexAop-mito-SypHer) prior to nerve stimulation. (S) The mito-SypHer ratio showing a response to nerve stimulation. (T) Plots of the mito-SypHer ratio at rest. (U) Plots of the greatest change in the mito-SypHer ratio (max ΔR/R), i.e. the ratio immediately prior to stimulation (rest levels) minus the peak response to nerve stimulation (lowest ratio) All terminals examined in HL6 (2mM CaCl_2_, 7mM L-glutamate) on segment 4. Nerves electrically stimulated at approximately twice the endogenous firing frequency of the terminal being examined; 80Hz for MN13-Ib (B, F and K) and 42Hz for MN6-Ib (O and S). Box plots show values from all preparations; mean (arrowhead), median (line), 25-75 percentiles box and 10-90% whiskers. Ratios normalized to 1 in panels B, F, I and K only. Mann-Whitney Rank Sum test in D; unpaired Student’s T-tests in all others.

The greater acidification in ArgK1 KD terminals was unexpected, and may reflect either a compensatory increase in glycolysis, as observed in mouse skeletal muscle when deficient in mitochondrial CK (van Deursen et al., 1993; Dzeja et al., 2011), or a greater influx of Ca^2+^ during activity. The latter would result in greater acidification, as the PMCA, a Ca^2+^/2H^+^ exchanger, extrudes much of the Ca^2+^ from these terminals after activity (Rossano et al., 2013). A greater PMCA-mediated exchange of Ca^2+^ for H^+^ was subsequently dismissed, as GCaMP8f revealed similar presynaptic Ca^2+^ levels during activity in ArgK1 KD terminals relative to controls (Figures 7J-7K).

ArgK1 is localized to mitochondria (Figures 5A-5C), and so to determine whether ArgK1 KD impacts presynaptic mitochondrial function we targeted their matrices with ratiometric reporters of Ca^2+^ and pH (Figures 7N-7U). Mitochondria in *Drosophila* larval MN terminals accumulate Ca^2+^ and this Ca^2+^ stimulates their energy metabolism (Chouhan et al., 2012). Here, mitochondrial TN-XXL (Ivannikov and Macleod, 2013) showed similar mitochondrial Ca^2+^ levels at rest between ArgK1 KD and control terminals (Figures 7N-7P), and similar levels of Ca^2+^ accumulation subsequent to stimulation (Figure 7Q). Mitochondrial matrix pH, relative to intermembrane space or cytosolic pH, represents part of the proton motive force used by the F_1_F_0_-ATPase to generate ATP (Nicholls and Ferguson, 2002). We targeted a ratiometric pH reporter, SypHer (Poburko et al., 2011), to the matrix (Figure 7R) where it showed that mitochondria in ArgK1 KD terminals were more acidic (Figures 7S-7T), but they acidified by a similar amount as they accumulated Ca^2+^ (Figure 7U). The reason for mitochondrial matrix acidification at rest is unknown, although without an arginine kinase associated with the adenine nucleotide translocase (ANT), exchange of ATP/ADP across the ANT may become inefficient and rate limiting for respiration.

## DISCUSSION

By tracking ATP expenditure rates within presynaptic terminals of known volume we have been able to determine ATP production rates (mM/s), and rates of acceleration (mM/s/s), required to stabilize ATP levels. Our quantitative approach shows that the intense episodic activity of MNs imposes an exceptional burden on the presynaptic metabolic machinery. Presynaptic energy demands can increase by an order of magnitude in milliseconds, and such volatility necessitates a rapid acceleration in ATP production to meet demand. However, our simulations indicate that ATP levels could exhaust before the ATP production machinery receives sufficient impetus from changes in concentrations of adenine nucleotides and inorganic phosphate, generating dissonance with the expectation that presynaptic metabolic demands are well matched by ATP synthesis (Rangaraju et al., 2014). However, our discovery of a dense colocalization of phosphagen kinases with presynaptic mitochondria, in both fruit flies and mice, allow for a reconciliation. In muscle, phosphagen kinases are essential elements in phosphagen systems capable of ATP regeneration rates exceeding the capacity of either glycolysis or Ox-Phos over the short term (Saupe et al., 1998; Saupe et al., 2000). Here, knock down of phosphagen kinases in *Drosophila* MNs led to energy metabolism deficits, corroborating a key role for phosphagen systems in presynaptic energy metabolism. To our knowledge, this is the first characterization of the presence, distribution and metabolic significance of phosphagen kinases in MN terminals, and warrants further exploration as a mechanism whose failure would surely represent a risk factor in the progression of some MN diseases.

Although not incorporated within our model, elevations in Ca^2+^ at the onset of activity can play a feedforward role in accommodating acceleration in energy demand. Ca^2+^ potentiates the rate of Ox-Phos through the stimulation of the mitochondrial aspartate-glutamate carrier, the ATP synthase, and Kreb’s cycle enzymes (Glancy and Balaban, 2012; Llorente-Folch et al., 2013). We show that, upon arrival of APs in the presynaptic compartment, power demand can elevate 10-fold in type-Ib terminals. As this is triggered by Ca^2+^ entry to the cytosol, mitochondrial Ca^2+^ uptake seems to represent an effective design for integrating power supply with demand. In *Drosophila*, the mitochondrial affinity for Ca^2+^ is similar between type-Is and -Ib terminals (Chouhan et al., 2010), but due to differences in MN firing rates cytosolic Ca^2+^ levels only rise into a range available to mitochondria in the hard working type-Ib terminals (Chouhan et al., 2010; Chouhan et al., 2012). It follows that mitochondria in type-Ib terminals will be operating at a higher rate of Ox-Phos during physiological activity. Hard working type-Ib terminals are therefore tuned to meet sharp elevations in energy demand through the combination of a Ca^2+^-mediated fillip to Ox-Phos (Chouhan et al., 2012; Ivannikov and Macleod, 2013), a high mitochondrial content (Justs et al., 2022) and a high density of phosphagen kinases shown in this study.

While the adaptations above can accommodate rapid changes in ATP demand, ATP levels can only be sustained during ongoing activity if glycolysis and Ox-Phos eventually rise to match demand. The magnitude of the increase in ATP demand from rest to sustained activity can provide some insight into the identity of the mechanisms that accommodate presynaptic energy demands. After only 10 peristaltic cycles, time-averaged energy demand summates to ∼0.6 mM/s in the type-Ib terminal of muscle fiber #6 (for example); a 10-fold increase in sustained power demand. While glycolysis has the capacity to accelerate rapidly (Walter et al., 1999), its total ATP production capacity must be considered. (Mookerjee et al., 2017) demonstrated that glycolysis in cultured myoblasts could increase ∼20-fold beyond a basal rate. However, such a range would only be sufficient to cover the 10-fold increase reported here if glycolysis provided all the power at rest. For example, if glycolysis provides a fraction of the power at rest, comparable with myoblasts (∼7%), then a 20-fold increase in glycolysis would provide an increase in total power output of only ∼2.3-fold [(20 x 7%_glyc_) + 93%_Ox-Phos_ = 2.3]. Therefore, without a substantial increase in Ox-Phos, the efforts of glycolysis alone will be ineffective at stabilizing ATP levels. In addition, high rates of glycolysis will be limited by NAD^+^ availability, and by a drop in pH, possibly caused by glycolysis itself (Erecinska et al., 1995; Ferguson et al., 2018). Although the rate of acceleration of Ox-Phos is inferior to that of glycolysis (Greenhaff and Timmons, 1998), Ox-Phos, nevertheless, can also increase 20-fold according to studies measuring OCR in skeletal muscle *in vivo* (Bangsbo et al., 1990). Therefore, as Ox-Phos provides a substantial fraction of the power at rest, a 20-fold range would enable Ox-Phos to cover the 10-fold increase required in this study [(20 x 93%_Ox-Phos_) + 7%_glyc_ = 18.7], and further reinforces the notion that Ox Phos is the primary source of ATP in these presynaptic terminals.

Type-Ib terminals, with their 10-fold increase in power demand in the rest-to-work transition might be compared with mammalian fast-twitch muscle fibers, while type-Is might be compared with slow-twitch fibers. However, contrary to the association of substantial episodic power demands with a heavy reliance on glycolysis across mammalian muscle fiber types (Schiaffino and Reggiani, 2011; Barclay, 2017), type-Ib terminals with their high power demand volatility have the greatest mitochondrial density (type-Ib: 6.3%; type-Is: 3.8%) and aggregate volume (Justs et al., 2022). This might be contrasted with the human *vastus lateralis* muscle, where the gradation of slow- to fast-twitch fibers is accompanied by a decrease in mitochondrial density (type-1: 6.0%; type-2A: 4.5%; type-2X: 2.3%) (Howald et al., 1985). Where the analogy does hold, is in the positive correlation between the duration of use of mammalian muscle fiber type and mitochondrial density. Type-Ib terminals, in addition to having more volatile power demands, are active for a greater proportion of the time (Lu et al., 2016; Newman et al., 2017). It follows that mammalian MNs might similarly be functionally differentiated, particularly with regard to energy metabolism. For this reason, a closer look at the energy metabolism of MNs innervating different mammalian muscle fiber types may be warranted. It is known that neuromuscular junctions on fast-fatigable muscle fibers are the first to be lost in several diverse mouse models of MN disease (Frey et al., 2000; Pun et al., 2006; Kanning et al., 2010), and it may be that MNs of these motor units also experience a volatile power demand with a heavy reliance on Ox-Phos and critical support from phosphagen kinases.

## EXPERIMENTAL PROCEDURES

### Solutions and Chemicals

*Drosophila* larvae were fillet dissected in a shallow bath with a Sylgard bed (Rossano and Macleod, 2007). All imaging was conducted on female larvae in hemolymph-like 6 (HL6) solution (Macleod et al., 2002). Tetramethylrhodamine ethyl ester (TMRE) was purchased from ThermoFisher; catalog no.T669.

### Fly Stocks

All *Drosophila* stocks were raised at 24°C on standard medium [recipe from Bloomington *Drosophila* Stock Center (BDSC), Bloomington, IN]. BDSC provided the following fly lines: P[PTT-GB]Argk1^CB03789^ (ArgK1-GFP, #51522), nSyb-GAL4 #51635, UAS-mito-GFP #8442, nSyb-LexA #52247, shakB-LexA #52775, and LexAop2-IVS-myrGFP #32209. BDSC also provided the RNAi line that expresses ArgK1 dsRNA (UAS-ArgK1-RNAi, #41697) [*Drosophila* Transgenic RNAi Project (TRiP) (Perkins et al., 2015)], and a control line that expresses firefly luciferase from a VALIUM10 vector (UAS-luciferase, #35788), the same vector used for RNAi lines. MN expression was driven using nSyb-GAL4, nSyb-LexA, OK6-GAL4 (Aberle et al., 2002) or type-Is specific shakB-LexA. Fluorescent reporters of cytosolic ATP/ADP and Ca^2+^ levels were gifts from Dr. Catherine Collins (UAS-PercevalHR) (Wong et al., 2014) and Dr. Dion Dickman (UAS-Syt::mScarlet::GCaMP8f) (Li et al., 2021), respectively. The fluorescent reporter of cytosolic pH (UAS-pHerry) (Rossano et al., 2017) was previously published from the Macleod laboratory. A transgenic line enabling expression of TagBFP free in the cytosol was a gift from Dr. Kenneth Irvine (UAS-TagBFP) (Rauskolb et al., 2014). Lines enabling expression of mKate free in the cytosol (UAS-cyto-mKate), and mKate targeted to the mitochondrial matrix (UAS-mito-mKate) were made in the Macleod lab as described below. The generation of fluorescent reporters of mitochondrial pH and Ca^2+^ by targeting SypHer (pKa ∼8.7) (Poburko et al., 2011) and TN-XXL (K_d_ ∼ 0.8 μM) (Mank et al., 2008) to mitochondrial matrices (LexAop-mito-SypHer and LexAop-mito-TN-XXL, respectively) are also described below.

### Generation of Transgenic Fly Lines

cDNA for mKate (Shcherbo et al., 2007) was acquired from Addgene and sub-cloned into the MCS of pUASt creating the UAS-cyto-mKate plasmid. Subsequently, cDNA for a tandem repeat of the first 36 AAs of subunit VIII of human cytochrome oxidase (2MT8) (Filippin et al., 2005) was sub-cloned 5’ of mKate to create the UAS-mito-mKate plasmid. cDNA for TN-XXL was a gift from Dr. Dirk Reiff at the Max Planck Institute for Biological Cybernetics in Tubingen, Germany, and sub-cloned into pJFRC19 3’ to the 2MT8 mitochondrial signal sequence to create the LexAop-mito-SypHer plasmid. cDNA for SypHer was a gift from Dr. Nicolas Demaurex at the University of Geneva and sub-cloned into the MCS of pJFRC19 to create the LexAop-mito-SypHer plasmid. UAS-mito-mKate and UAS-mito-mKate plasmids were injected into w^1118^ *Drosophila* embryos, and both LexAop-mito-TN-XXL and LexAop-mito-SypHer plasmids were injected into embryos containing the attP40 landing site, all by Rainbow Transgenic Flies (Newbury Park, CA). Strains with transformed germ line cells were mapped for the inserted chromosomes, balanced, and other chromosomes out-crossed to an in-house w^1118^ line.

### *Drosophila* Confocal Imaging

Larvae were dissected and imaged in HL6, with CaCl_2_ added to 0.1 mM. Type-Ib and -Is nerve terminals on muscle fibers #7, #6, #13 and #12 of segment 4 were imaged on a Nikon A1R confocal microscope with either of two Nikon water-immersion objectives; 60X 1.20NA Plan Apo VC or 20X 0.75NA Plan Fluor. For demonstrations of localization, Z-stacks were taken through the entire depth of terminals and rendered as maximum projections, whereas for quantification Z-stacks were rendered as average projections. Lasers (405, 488 or 561 nm) were used sequentially and emission filters were optimized for BFP, GFP or mKate (or TMRE), respectively. A pinhole size of 1 Airy unit was used throughout. Mitochondrial content (Figure 5F and S5) was calculated by dividing the terminal’s GFP (mito) fluorescence intensity by the sum of the terminal’s GFP (mito) and TagBFP (cytosol) intensity. ArgK1-GFP fluorescence intensity was normalized to a correlate of terminal volume (Figure 4J-4K) rather than the intensity of a cytosolic fluorophore to side-step confounds caused by differences in the strength of GAL4 expression between terminal types. As the volume of a varicose axon (terminal) has a linear relationship with the square of its maximum cross-sectional area (plan area of a projection), we divided the amount of ArgK1-GFP fluorescence (from the average projection of a stack) along a 20 μm length of terminal by the square of the cross-sectional area along the same length of terminal. All measurements of fluorescence intensity and cross-sectional area were made in ImageJ (imagej.nih.gov/ij).

### Presynaptic Cytosolic Functional Imaging

Functional imaging on fillet-dissected larvae was performed under an Olympus water-dipping 100X 1.00 NA LUMPlanFl objective on a Nikon Eclipse FN1 microscope fitted with two Andor iXon3 EMCCD cameras (DU-897) mounted on an Andor TuCam beam-splitter (Andor Technology, South Windsor, CT). Filters and dichroic mirrors were obtained from Chroma Technology (Bellows Falls, VT, USA) or Semrock (Lake Forest, IL, USA). Nikon Elements software controlled the illumination sequence and camera image acquisition. A Master 8 (A.M.P.I., Israel) controlled the timing of nerve stimulation via an A-M Systems Model 2100 Isolated Pulse Stimulator. Nerves were not severed, but rather, a loop of the nerve was drawn into a stimulating pipette. Terminals were allowed to equilibrate for at least 2 minutes after any nerve stimulus train.

Changes in the ratio of ATP concentration relative to ADP were monitored through changes in the PercevalHR fluorescence ratio (Tantama et al., 2013). PercevalHR was excited with two different wavelengths by a Lumencor SPECTRA-X light source and emission was collected at a single wavelength. The light source alternated illumination to provide 100ms of 436/20nm excitation interleaved with 100ms of 483/32nm excitation, each reflected off a 505nm edge dichroic mirror. The emitted light passed through a 520/50nm filter and was collected at ∼8 frames-per-second (fps) by a single camera. Image data were analyzed as described previously (Macleod, 2012) using ImageJ. The ratio of intensities from the image pairs, relative to the ratio prior to nerve stimulation (ΔR/R), was plotted against time, to yield ∼4 ATP/ADP estimates (PercevalHR ratios; 483nm ex./436nm ex.) per second (Figure 7B). A decrease in ΔR/R, indicates a decrease in ATP relative to ADP.

Changes in cytosolic pH were monitored through the change in the pHerry fluorescence ratio of pH-sensitive superecliptic pHluorin (SEpHluorin) (Sankaranarayanan et al., 2000) to pH-insensitive mCherry. pHerry was excited with two different wavelengths and emission was collected at two different wavelengths. The light source alternated illumination to provide 100ms of 483/32nm excitation interleaved with 100ms of 550/25nm excitation, reflected by a quad-band dichroic mirror (Semrock FF409/493/573/652). The emitted light passed through a quad-band emission filter (430nm/36nm, 513/44, 595/41, 719/109) before SEpHluorin fluorescence was reflected by a second dichroic mirror (edge at 580nm) through a 525/50nm filter to one camera, and mCherry fluorescence passed through a 617/73nm filter to a second camera. ΔR/R was plotted against time, to yield ∼4 cytosolic pH estimates (pHerry ratios; 483nm ex./550nm ex.) per second (Figure 7F). SE-pHluorin fluorescence decreases in response to acidification while mCherry is unresponsive (Rossano et al., 2017).

Changes in the cytosolic Ca^2+^ concentration [Ca^2+^]_c_ were monitored through the change in the ratio of Ca^2+^-sensitive GCaMP8f to Ca^2+^-insensitive mScarlet, using the same imaging protocol as for pHerry above. ΔR/R was plotted against time, to yield ∼4 [Ca^2+^]_c_ estimates (GCaMP8f to mScarlet fluorescence ratios; 483nm ex./550nm ex.) per second (Figure 7K). The GCaMP8f to mScarlet ratio increases in response to an increase in [Ca^2+^]_c_ (Li et al., 2021).

### Presynaptic Mitochondrial Functional Imaging

Imaging was performed on the “rig” described for cytosolic imaging, but with a Nikon water-dipping 60X 1.20NA Plan Apo VC objective. As above, larvae were fillet dissected and nerves were not severed. Terminals were allowed to equilibrate for at least 5 minutes after any nerve stimulus train.

Changes in mitochondrial matrix Ca^2+^ levels ([Ca^2+^]_m_) were monitored through changes in the mitochondrial TN-XXL emission ratio of Citrine to CFP (Mank et al., 2008). TN-XXL was excited by a single wavelength and emission was collected at two different wavelengths. TN-XXL was excited at 436/20nm, which passed through a triple-band dichroic mirror (Chroma, 69008bs). The emitted light passed through a triple-band emission filter (470/24, 535/30, 632/60) before a second dichroic mirror (edge at 509nm) reflected CFP emission through a 475/28nm filter to one camera and Citrine emission was passed through a 550/49nm filter to the other camera. Each camera collected light at ∼10 fps. The ratio of intensities from the image pairs, relative to the ratio prior to nerve stimulation (ΔR/R), was plotted against time to yield ∼10 mitochondrial Ca^2+^ estimates (mito-TN-XXL ratios) per second (Figure 7O).

It is likely that not all TN-XXL is translocated into mitochondrial matrices, and that some TN-XXL remains in the cytosol, as the fluorescence decay upon nerve stimulation cessation is far more rapid than the decay quantified using non-saturating chemical Ca^2+^ reporters. Mitochondrial Ca^2+^ decays monotonically over several minutes (Chouhan et al., 2012; Macleod and Ivannikov, 2017), while cytosolic Ca^2+^ decays back to rest levels over hundreds of milliseconds subsequent to high frequency nerve stimulation (Macleod et al., 2004). Therefore, the stimulated change in mitochondrial Ca^2+^ concentration was quantified as being proportional to the change in the ratio 10 seconds after cessation of stimulation relative to the level immediately prior to stimulation (rest levels). At a time 10 seconds after stimulation cessation, cytosolic Ca^2+^ levels have returned to baseline, and therefore any elevation in the Citrine/CFP ratio will represent the mitochondrial, rather than cytosolic, Ca^2+^ response.

Changes in mitochondrial matrix pH were monitored through changes in the mitochondrial SypHer emission ratio (Poburko et al., 2011). Imaging was performed on the “rig” described above, but unlike imaging mito-TN-XXL, a 100X 1.10 NA Nikon objective was used and mito-SypHer was excited with two different wavelengths and emission was collected at a single wavelength. The light source alternated illumination to provide ∼50ms of 436/20nm excitation interleaved with ∼50ms of 483/32nm excitation, each reflected off a 505nm dichroic mirror. The emitted light was passed through a 525/50nm filter and reflected off a 580nm edge dichroic to a single camera collecting at ∼20 fps. ΔR/R was plotted against time to yield ∼10 mitochondrial pH estimates (mito-SypHer ratios) per second (Figure 7S).

### Mouse Immunohistochemistry and Confocal Imaging

All mice were used in compliance with the ethical standards established by the University of Nevada, Reno, and NIH guidelines (NRC, 2011). Primary euthanasia was performed using isoflurane, and secondary euthanasia was performed by exsanguination. Levator auris longus (LAL) muscles were removed bilaterally from 2-month old mice. After removal, LALs were pinned to Sylgard-coated dish and fixed in 4% PFA for 20 minutes, following previously established procedures (Wright et al., 2011). Tissue was blocked with 0.1M glycine/PBS for 30 minutes, permeabilized with cold methanol for 5 minutes, and then blocked a second time using antibody blocking buffer (1% fish gelatin, 0.5% Triton-X 100, 0.025% NaN_3_, in 0.1M PBS) for 1 hour. Three 10-minute washes with 0.1M PBS occurred between each blocking and permeabilization step.

Rabbit primary antibodies CKMT1A (Proteintech, 1:500, Cat. #: 15346-1-AP) and CKMT1B (Invitrogen, 1:500, Cat. #: PA5-96224) were applied to separate tissues incubating in a primary goat antibody for the vesicular acetylcholine transporter (vAChT; MilliporeSigma, 1:1000, Cat. #: 618250), and incubated at 4⁰C on a shaker overnight. After three 10-minute washes with 0.1M PBS, Alexa Fluor 647 (Jackson ImmunoResearch, 1:500, Cat #: 705-605-003) and Alexa Fluor 488 (Invitrogen, 1:500, Cat #: A21206) secondary antibodies were added and incubated overnight at 4⁰C. After washing with PBS, the tissue was mounted in Slowfade™ Diamond Antifade mountant (ThermoFisher, Cat# S36963), and imaged on an Olympus FV1000 confocal microscope with a 60x (NA 1.4) oil immersion objective. Images in Figures 5G and 5I were taken using an Olympus FV1000 scope using a 40X 1.30 NA oil-immersion objective, digitized at 218 and 230nm/pixel, respectively. Figure 5H was taken using a Leica Stellaris X8 scope with a 93X 1.30 NA glycerol-immersion objective, digitized at 50nm/pixel.

### Computational Modeling

#### Overview

The computational model simulates the dynamics of the presynaptic ATP concentration ([ATP]), subject to our previous ATP consumption estimates for *Drosophila* motor neuron terminals (Justs et al., 2022), alongside an ATP production model adapted from (Wilson, 2017a). Figure S7 summarizes the metabolite concentrations included in the model and the processes that modify those concentrations over time.

#### Equilibration

Equilibration is assumed to be instantaneous, i.e., the equilibrium equations are always satisfied:

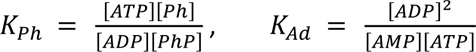

K_Ad_ is equal to 1 (Wilson, 2017a). K_Ph_ depends on the organism, see Table S4.

#### ATP Consumption

Following (Justs et al., 2022), we modeled the presynaptic ATP consumption rate (power) as the sum of a time-dependent signaling rate capturing the energetic cost of triggering neurotransmitter release in response to an action potential (AP), and a constant rest rate capturing all other cell activities (protein/lipid synthesis, cytoskeleton turnover, ongoing Na^+^/K^+^ ATPase activity at the plasma membrane, etc). The consumption rate associated with a single AP is the sum of five distinct signaling processes: Ca^2+^ pumping, Na^+^ pumping, and three types of neurotransmitter recycling processes (NT1, NT2, NT3).

The ATP consumption rate for a single signaling process is zero before the AP, jumps to a nonzero value as an AP initiates the process, then decays exponentially (Figure S8A):

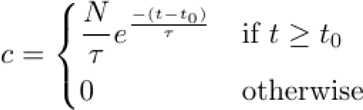

where t_0_ is time of the AP, N is the process’ total ATP cost, and τ is the process’ decay time. The total signaling rate of ATP consumption is obtained by adding up the rates of each of the five processes, then adding up the contribution of every AP. Locomotion in *Drosophila* larvae is driven by cyclical contractions of body wall muscles, with a frequency of ∼ 1/sec (Berrigan and Pepin, 1995), where each contraction is driven by a burst of APs in the MNs. Figure S8B shows an example of such AP bursts.

The rest ATP consumption rate (c_rest_), the total ATP cost of each signaling process (N), the AP firing pattern, and the total terminal volume were quantified for six different *Drosophila* motor neuron terminals in (Justs et al., 2022) (Table S5). The time courses of Ca^2+^ and Na^+^ pumping were given in the Results section [38.9 ms, an average of all six terminals (Justs et al., 2022); 63.5 s (Mondragao et al., 2016), respectively] (Table S6). The time course of SV recycling, the mechanisms that regulate it, and the relative ATP costs of each recycling process is an ongoing area of research. There are four different mechanisms known to be involved in SV recycling: ultrafast endocytosis (UFE), “kiss-and-run” endocytosis, clathrin-mediated endocytosis (CME), and activity-dependent bulk endocytosis (ADBE) (Kononenko and Haucke, 2015). Each mechanism is differentiated by various features such as the time constant for membrane retrieval, clathrin-dependency, Ca^2+^-dependency and the different molecular components involved in the membrane retrieval process. Drawn from a number of sources, the time course of endocytosis for each mechanism is: UFE (50-500ms), “kiss-and-run” (<1 s), CME (10-20 s), and ADBE (seconds) (Kononenko and Haucke, 2015; Soykan et al., 2016). The time courses for these mechanisms have not been defined in *Drosophila* larval MN terminals, but it is known SV recycling does involve CME and ADBE in *Drosophila* (Gan and Watanabe, 2018). UFE has not been reported from *Drosophila*, but UFE has been demonstrated at the *C. elegans* NMJ (Watanabe et al., 2013) and is likely conserved in *Drosophila* (Gan and Watanabe, 2018). In the absence of a set of time constants reflecting all three mechanisms at *Drosophila* synapses, time constants used for SV recycling (and refilling) in this manuscript are based on data obtained by (Chanaday and Kavalali, 2018). Under physiological conditions (2 mM Ca^2+^, 34°C), at rat hippocampal synapses, they determined three time constants for SV recycling: ultrafast (NT1: ∼175 ms, ∼13% events), fast (NT2: ∼7.21 s, ∼34% of events) and ultraslow speeds (NT3: >20 s, ∼53% of events). They were unable to ascertain the extent to which “kiss-and-run” may have given rise to the ultrafast time course, and could not conclude whether the ultra-slow mechanism was due to ADBE or due to components that diffuse on the plasma membrane and re-cluster later; e.g. (Gimber et al., 2015). While it is recognized that these data may not be ideal for modeling SV recycling and refilling in *Drosophila*, they are an internally consistent set collected under the very same conditions and such a cohesive set is not available for *Drosophila*.

#### ATP Production

The ATP production rate is based on Wilson’s Ox-Phos model, which captures the ability of mitochondria to increase production many fold in response to relatively small drops in [ATP] in order to maintain [ATP] homeostasis. First, we compute the energy state [ATP]/([ADP][P_i_]). Then, we use the model and parameters described in (Wilson, 2017a) to compute the cytochrome c turnover rate. The ATP production rate is equal to this turnover rate times the cytochrome c concentration. The latter is inferred by requiring the values of [ADP], [P_i_] and [ATP] found in the literature (see *Organism-Specific Parameters* below) to yield an ATP production rate equal to the terminal’s rest consumption rate:

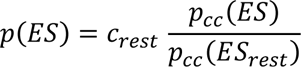

where p is the ATP production rate, p_cc_ is the cytochrome c turnover rate, c_rest_ is the rest-level ATP consumption rate, ES is the energy state, and ES_rest_ is the value of the energy state computed from the [ADP], [P_i_] and [ATP] values from the literature. Finally, we impose a maximum ATP production rate p_max_ (see *ATP Production Limits* below) as follows:

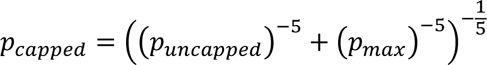

This ensures a smooth transition from the Wilson model when the energy state is large to the maximum rate p_max_ when energy state is low (Figure S9).

#### ATP Production Limits

The (Wilson, 2017b) model recapitulates empirical Ox-Phos ATP production rates, but at very low energy state values it predicts rates that exceed estimates gained from most experiments performed *in vivo,* and all experiments *in vitro*. We therefore sought to apply ATP production rate limits according to the mitochondrial volume of each motor neuron terminal. To do this, we needed *in vivo* estimates of the maximal bioenergetic output from neurons with known mitochondrial volumes. However, *in vivo* estimates have not been achieved, primarily due to the difficulties of isolating critical measures of energy metabolism *in vivo*, such as O_2_ consumption rates (OCRs) and extracellular pH change, from populations of neurons that have been maximally activated. We therefore built estimates of maximal bioenergetic output by drawing on values from disparate sources and presented these estimates for a number of cell types, under various conditions.

##### Myoblasts *in vitro*

Perhaps the best controlled estimates of maximal bioenergetic output come from C_2_C_12_ cells *in vitro* (Mookerjee et al., 2017). Under conditions of sequential and complete oxidation of glucose by glycolysis and Ox Phos, these cells produce 44.8 pmol ATP/min/ug of protein from Ox Phos. Glycolysis can supply ATP at a higher rate, but only for a short period of time as it is limited by NAD^+^ supplies and the buildup of its own products, such as lactic acid (Erecinska et al., 1995; Yellen, 2018). (Mookerjee et al., 2017) determined that the maximum sustainable rate of ATP production occurs when Ox Phos decreases to 36.0 pmol ATP/min/ug of protein, but glycolysis increases to 24.4 pmol ATP/min/ug, yielding a combined output of 60.3 pmol ATP/min/ug. If myoblasts have a specific gravity of ∼1.06, and each 1 ug contains 195 ng of protein (Segal et al., 1986), then these myoblasts can produce 12.46 pmol ATP/min/nL of cell volume (0.208 mM ATP/s). This limit can then be applied to different motor neuron terminals, prorated according to their individual mitochondrial volume density. If the mitochondrial volume density of myoblasts is 6.5% (St-Pierre et al., 2003), we can calculate the maximum bioenergetic output as 0.032 mM ATP/s for each 1% of mitochondrial volume density.

For reasons not fully understood, *in vivo* estimates of maximal Ox Phos ATP production are substantially greater than estimates gained from isolated mitochondria or from cells out of their tissue context (Tonkonogi and Sahlin, 1997; Korzeniewski, 2007). Therefore, we sought estimates of maximal ATP production from different cell and tissue types *ex vivo* and *in vivo*, as follows:

##### Drosophila larval brain ex vivo

OCR measurements from whole larval brains in a respirometer (Seahorse analyzer) yielded estimates of 12.8 pmol ATP/min/mg (Neville et al., 2018); assuming that the consumption of one oxygen molecule signals the net production of 5.58 ATP molecules, i.e. a P/O ratio of 2.79 (Mookerjee et al., 2017). If the whole brain has a specific gravity of ∼1.1 (Barber et al., 1970), then the mix of neurons and glial cells produce 14.12 pmol ATP/min/nl of cell volume by Ox Phos (0.235 mM ATP/s). As the brain is enclosed within a tough sheath and glia within the brain condition the interstitial fluid, extracellular acidification measurements are difficult to interpret, and glycolytic output is difficult to estimate. To side-step this complication, we used the quantitative model of respiration provided by (Mookerjee et al., 2017) to calculate combined output from Ox Phos and glycolysis (JATP<ox> + JATP<glyc>) based on the ATP output of Ox Phos (JATP<ox>) calculated from OCR, i.e. we multiplied the JATP <ox> by the theoretical fraction of [(JATP<ox> + JATP <glyc>) / JATP <ox>], or 60.3/36.0, yielding 0.394 mM ATP/s. The overall mitochondrial volume density of the adult and larval brains is unknown, but estimates have been made from specific neuron types in the peripheral nervous system of both the larva and adult [3.8% and 6.3% (Justs et al., 2022); 5.6% (Zhu and Sun, 2013)]. As neurons represent the cells with the highest mitochondrial volume density, the overall density is likely to be closer to the lower end of this range. Proceeding with an estimate of 5%, we calculated a maximum bioenergetic output from Ox Phos and glycolysis combined of 0.079 mM ATP/s/1% mitochondrial density.

##### *Drosophila* adult fight muscle *ex vivo*

Respirometry measurements of OCR from permeabilized adult thoraces yielded estimates of 151 pmol ATP/min/ug assuming a P/O ratio of 2.79 (Jorgensen et al., 2021; Menail et al., 2022). Given a nominal muscle specific gravity of 1.06, this value increases to 160 pmol ATP/min/nl or 2.66 mM ATP/s. Using the model of Mookerjee and others (2017), the maximum combined output of Ox Phos and glycolysis is calculated to be 4.46 mM ATP/s. *Drosophila* flight muscle has a mitochondrial volume density of 29% (Perkins et al., 2012) and so the maximum bioenergetics output can be expressed as 0.154 mM ATP/s/1% mitochondrial density.

##### Human cortical grey matter *in vivo*

Using phosphorus magnetic resonance spectroscopy (^31^P MRS), Zhu and others (2012) estimated that active grey matter can produce 9.5 mmol ATP/min/g tissue (wet weight). Using a specific gravity of 1.1, this value converts to 10.45 mmol ATP/min/ml tissue, or 0.174 mM ATP/s, which represents a combined output from glycolysis and Ox Phos. Although active, presumably the grey matter of humans was not “maximally” activated. Mitochondrial volume density values of 7.05%, obtained from primate cortex (Bertoni-Freddari et al., 2007), allow us to calculate a bioenergetic output of 0.025 mM ATP/s/1% mitochondrial density.

##### Human quadriceps muscle *in vivo*

Surprisingly, the highest estimates of Ox Phos ATP production come from the *vastus lateralis* of the quadriceps when men cycled at a maximal work load and the OCR was estimated from blood vessels perfusing the quadriceps (Andersen and Saltin, 1985). Assuming a muscle specific gravity of 1.06 (Segal et al., 1986), a P/O coupling ratio of 2.79 (Mookerjee et al., 2017) and fully coupled Ox-Phos, the OCR of the quadriceps indicated an ATP production rate of 1.37 mM/s (Andersen and Saltin, 1985; Blomstrand et al., 1997). Using the model of Mookerjee and others (2017), we calculated the maximum combined output of Ox Phos and glycolysis to be 2.29 mM ATP/s. Muscles of the quadriceps group have a variable mitochondrial density, but proceeding with an average value of 6% (Johnson et al., 1973; Hoppeler, 1990) we calculated the maximum combined output of Ox Phos and glycolysis to be 0.382 mM ATP/s/1% mitochondrial density.

#### Organism-Specific Parameters

We tailored our model to different tissues by changing the initial metabolite concentrations, the tissue temperature, and the phosphagen equilibrium constant K_Ph_ (Table S4). We used squid axoplasm as a model invertebrate. For vertebrates, which utilize creatine rather than arginine, we use two metabolite concentration sets, one from human gray matter and one from human skeletal muscle. Note that, in the absence of direct terminal-level measurements, both are measured at the tissue level as proxies for terminal-level concentrations. Also, experimentally measured [ADP] inevitably includes cytoskeleton-bound ADP, which is not available for phosphorylation at mitochondria and should not be included in the Wilson model. We infer free [ADP] from the measured concentrations of ATP, Ph, PhP and the phosphagen equilibrium constant: 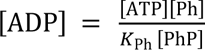. For the squid axoplasm without phosphagen, this approach is not applicable. Instead we use [ADP] = 0.02 mM from (Chance et al., 1986), which is close to the value used for the squid axoplasm with arginine ([ADP]=0.024 mM).

#### Numerical Integration

The time evolution of the system is obtained by numerical integration. ATP consumption and production are handled through the Euler integration scheme below:

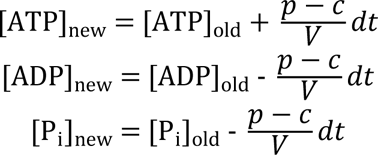

where c is the consumption rate, p is the consumption rate, V is the terminal’s volume, and dt is the time step. The phosphagen exchange reaction and the adenylate kinase reaction are then brought back to equilibrium using a nonlinear solver before moving on to the next time step. The simulation code is available as Supplementary File 1.

### Statistical Analysis and Data Presentation

Tests were performed using SigmaStat 13.0 software, and each test is described where used (Figures 1, 4, 5 and 7). Propagation of uncertainty theory (Farrance and Frenkel, 2012) was used to calculate variance of means based on uncertainty measurements (SEM) combined from different experiments (Figure 1C and 1D). Pearson product-moment correlation coefficient was calculated to test the strength and direction of associations, and the ordinary least-squares method was used to provide linear fits (Figure 1E). Functional imaging data were assessed for outliers using the median absolute deviation (MAD) (Leys et al., 2013) where an outlier was considered to be any value beyond 3 X MAD of the median and subsequently removed (Figure 7).

## Acknowledgements

This work was supported by NIH NINDS awards NS061914 and NS123377 to GTM, NS117686 to RBR, and NSF BIO-1943514 to RBR. Karlis Justs’ current affiliation is with Agilent Technologies USA. Stocks obtained from the Bloomington Drosophila Stock Center (BDSC: NIH P40OD018537) were used in this study. We are grateful for discussions with Profs. Paul De Weer, Kai Kaila and Gary Yellen.

## Supplemental Figures and Tables

**Figure S1.**
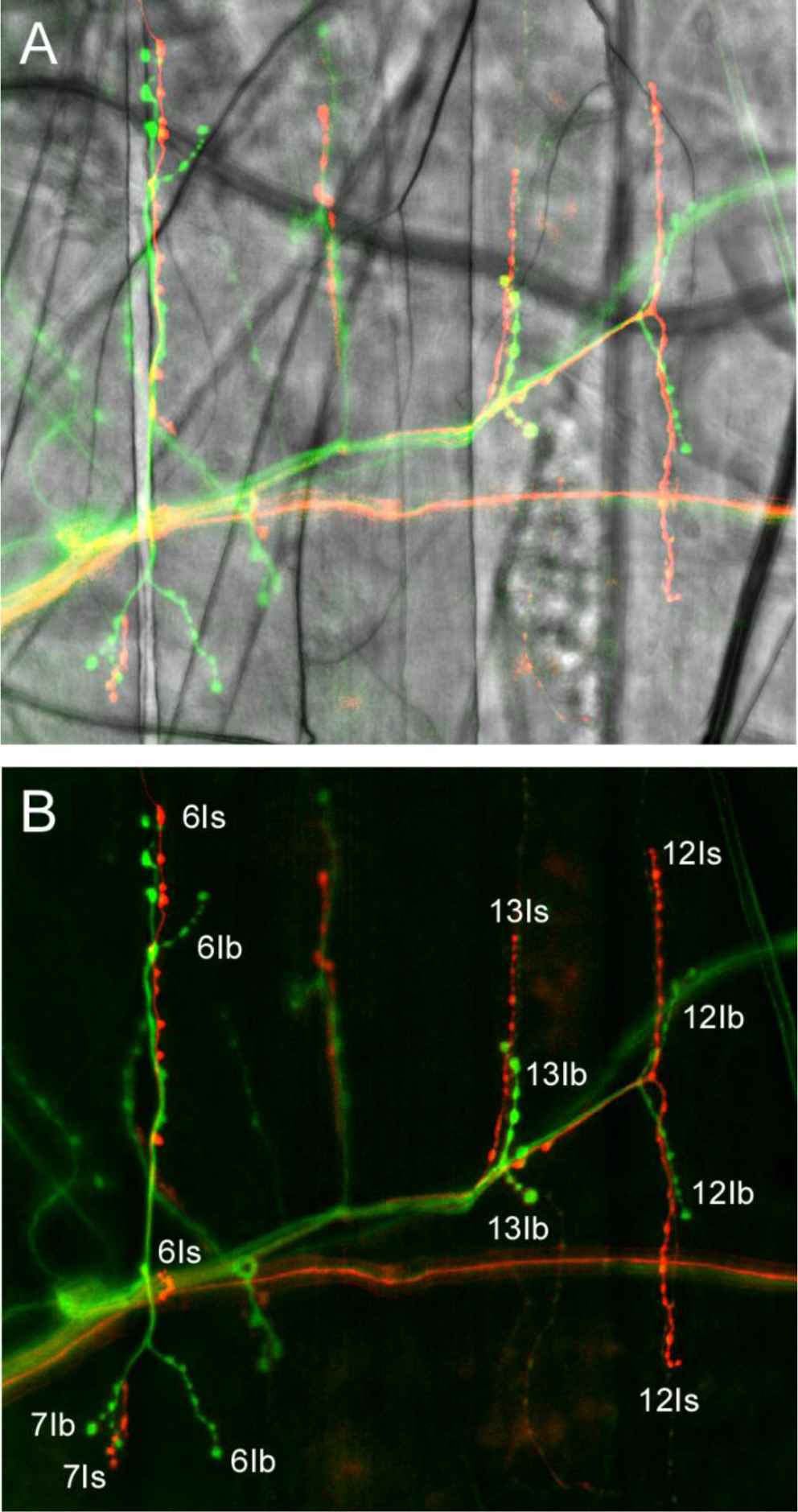
*Drosophila* Larval Motor Neurons (MNs) with Different Terminal Types Can be Revealed with Different Neuronal Drivers. (A) The stereotypical innervation of *Drosophila* larval body-wall muscle fibers #7, #6, #13 and #12, revealed in a confocal micrograph of green fluorescence (shakB-LexA > LexAop2-IVS-myrGFP) emitted from MNSNb/d-Is and its type-Is terminals (represented here in red) and blue fluorescence (nSyb-GAL4 > UAS-cyto-TagBFP) of MN6/7-Ib, MN13-Ib and MN12-Ib; three MNs with type-Ib terminals (represented here in green). Body wall muscle fibers, revealed in the overlain transmitted light image, run top-to-bottom beneath the fluorescent terminals. Note: while nSyb-GAL4 also expresses in type-Is terminals, they do not appear green in the image, as the fluorophore driven by ShakB-LexA in the type-Is terminals is relatively much brighter. (B) The same terminals in A, identified without the transmitted light image of the muscle fibers.

**Figure S2.**
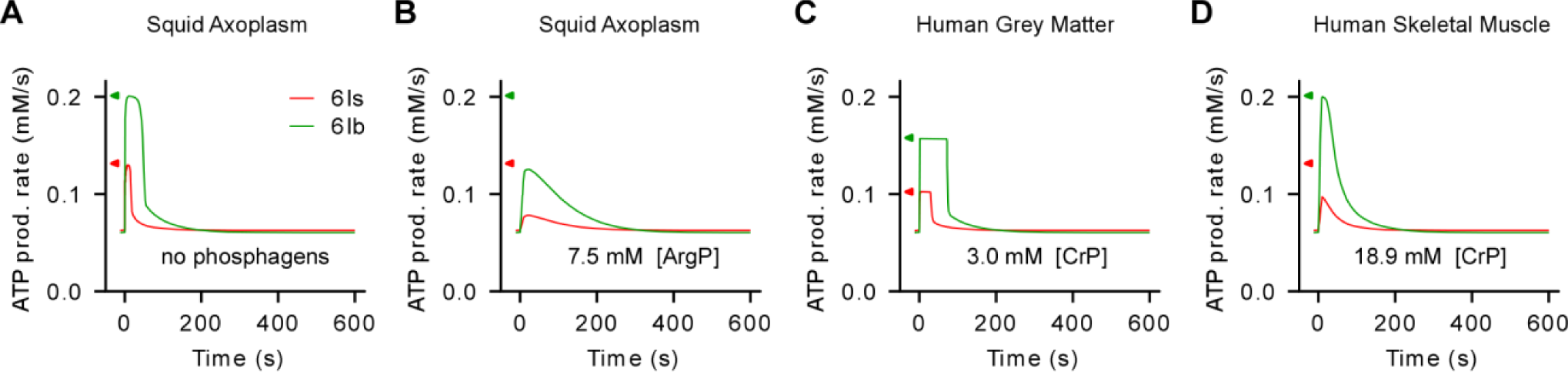
The Total Amount of ATP Produced (Area Under the ATP Production Curve) is Equal Between ATP Production Models. (A-D) Plots of the ATP production rates for terminals of MN6/7-Ib and MNSNb/d-Is on muscle fiber #6 over a period of 10 consecutive peristaltic cycles and the subsequent 9 minutes and 50 seconds. Panels A-D can be matched to panels D, G, J and M in Figure 6, respectively, i.e. the conditions are identical. The ATP production limits shown on the ordinates are the same as those calculated for Figure 6 (Ib: green; Is, red). Curves of the same color (corresponding to the same terminal) have the same area under the curve.

**Figure S3.**
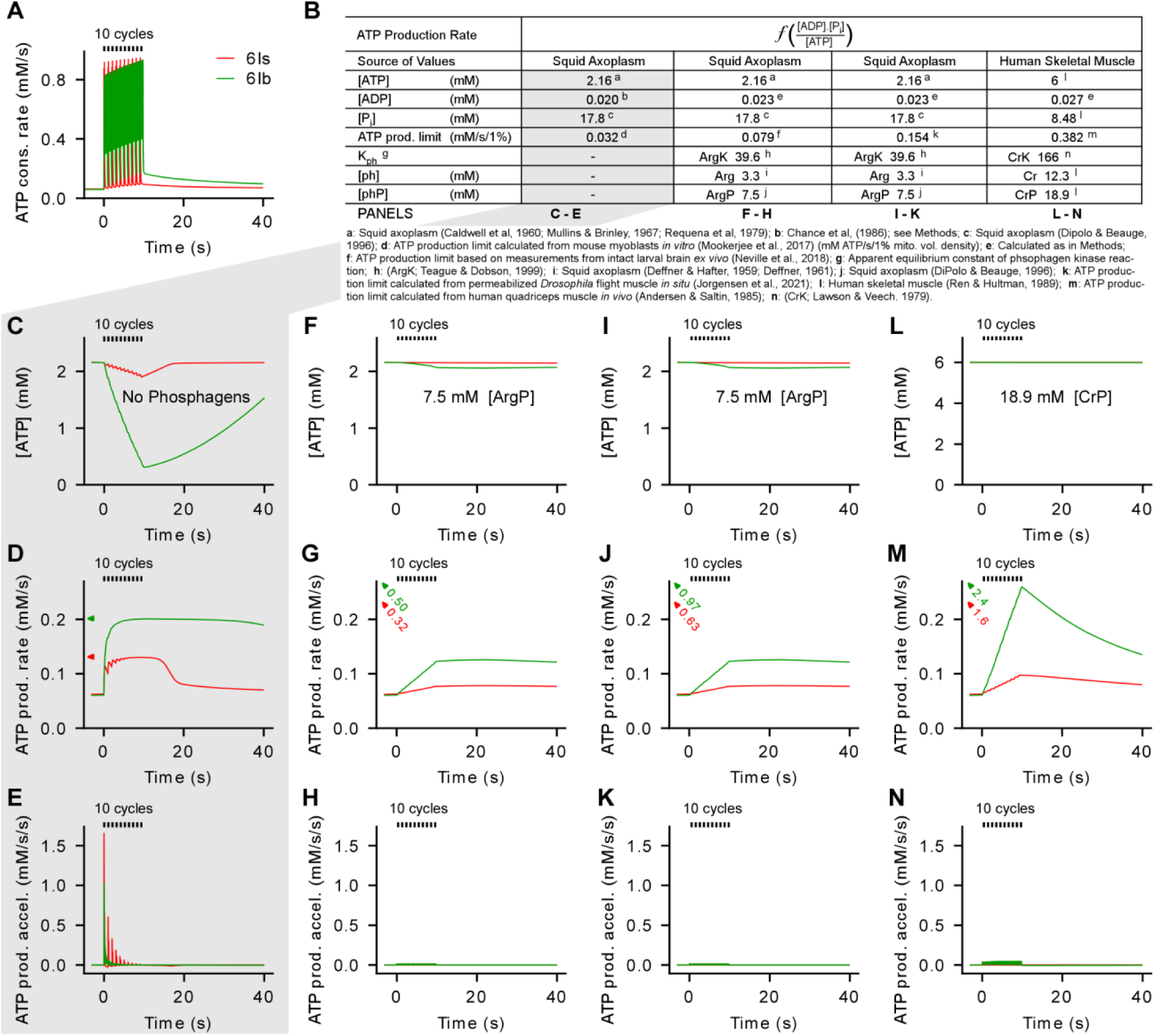
The High ATP Production Capacity Associated with Cells in a Tissue Context is Not Called Upon to Support Motor Neurons (MNs) During Short Periods (10 seconds) of Activity. (A) Plots of the instantaneous ATP consumption rate for terminals of MN6/7-Ib and MNSNb/d-Is on muscle fiber #6 only over a period of 10 consecutive peristaltic cycles. (B) A table showing parameter values from different animals and tissue types (columns 2-5) as the bases for simulations under the variable ATP production model (a function of the cellular energy state ([ADP].[P_i_]/[ATP]). Each parameter is introduced in column 1, and parameter values are further annotated with notes that appear below the table. The ATP production limits are drawn from cells under *in vitro* conditions in the first simulation (column 2; C-E), and ex-vivo or in vivo tissues in the remaining simulations (columns 3-5; F-N). The tissues do not necessarily match the animals and tissue type appearing at the top of the column. The ATP production limits (mM ATP generated per second for each 1% mitochondrial volume density) are applied to each terminal type according to each terminal’s mitochondrial volume density (Ib 6.29%; Is 4.10%), and are shown on the ordinates in D, G, J and M (Ib: green; Is, red). (C-E) Plots of the cytosolic ATP concentration (C), ATP production rate (D) and ATP production rate acceleration (E) during the period of activity shown in A, and in the absence of a phosphagen system (column 2 of table in panel B) (F-H) As in C-E, but with a phosphagen system, and using parameters from column 3 of the table in B. (I-K) As in C-E, but with a phosphagen system, and using parameters from column 4 of the table in B. (L-N) As in C-E, but with a phosphagen system, and using parameters from column 5 of the table in B.

**Figure S4.**
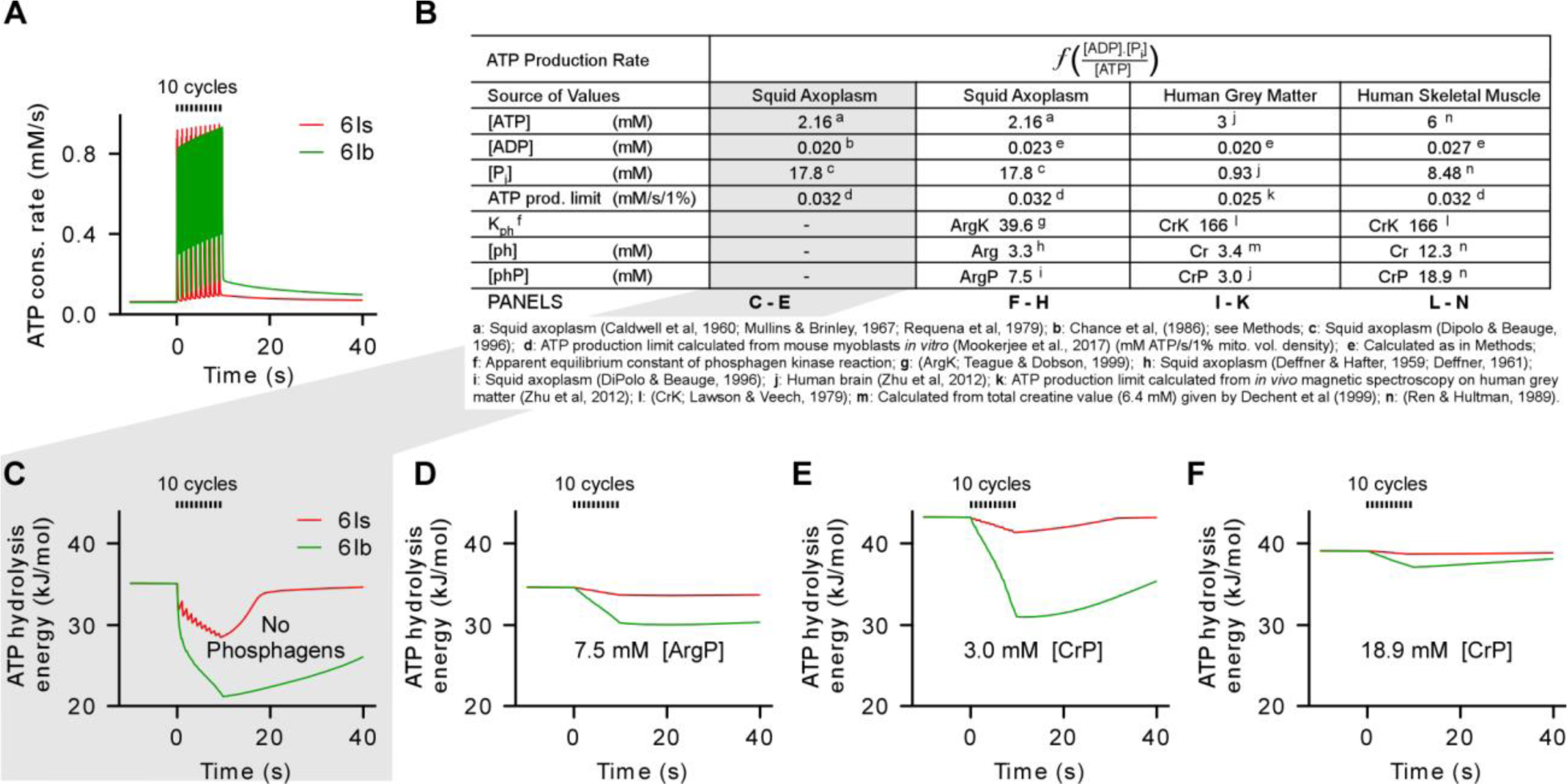
The Free Energy Available in ATP Hydrolysis Falls Faster Than the Cytosolic ATP Concentration. (A) Plots of the instantaneous ATP consumption rate for terminals of MN6/7-Ib and MNSNb/d-Is on muscle fiber #6 only over a period of 10 consecutive peristaltic cycles. (B) A table showing parameter values from different animals and tissue types (columns 2-5) as the bases for simulations under the variable ATP production model, as described for Figure 6. (C-F) Each plot shows the fall in the free energy of ATP hydrolysis for the two different terminals, under each set of parameter values.

**Figure S5.**
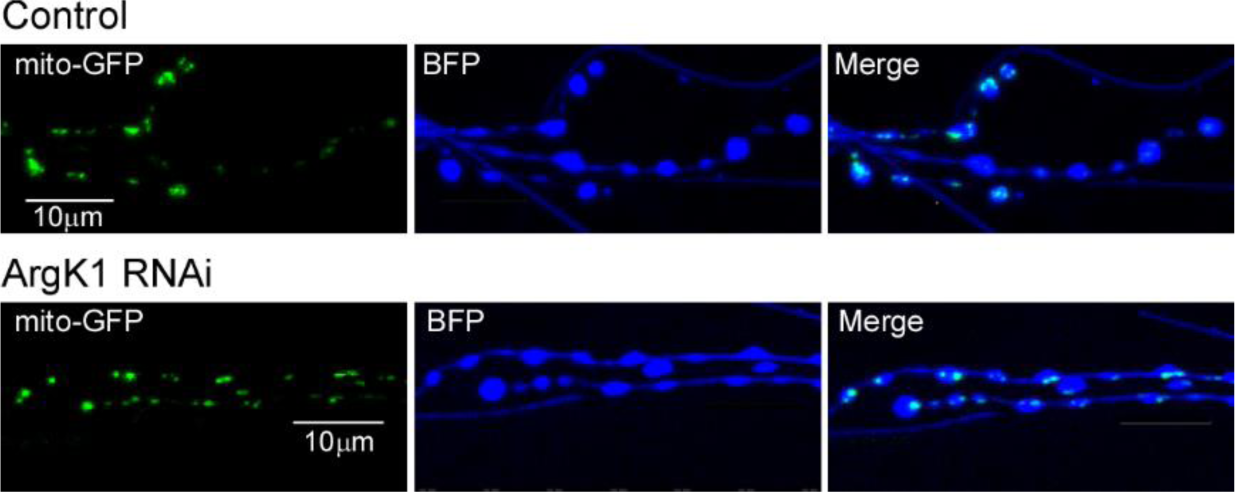
Presynaptic Mitochondrial Content is Unchanged by RNAi Mediated Knock Down of ArgK1. Confocal stack projections through NMJs on muscle fiber 6. Mito-GFP, cytosolic BFP and either ArgK1 dsRNA or firefly luciferase (control) all expressed with a single copy of a motor neuron (MN) driver (OK6-Gal4). Quantification in Figure 5F.

**Figure S6.**
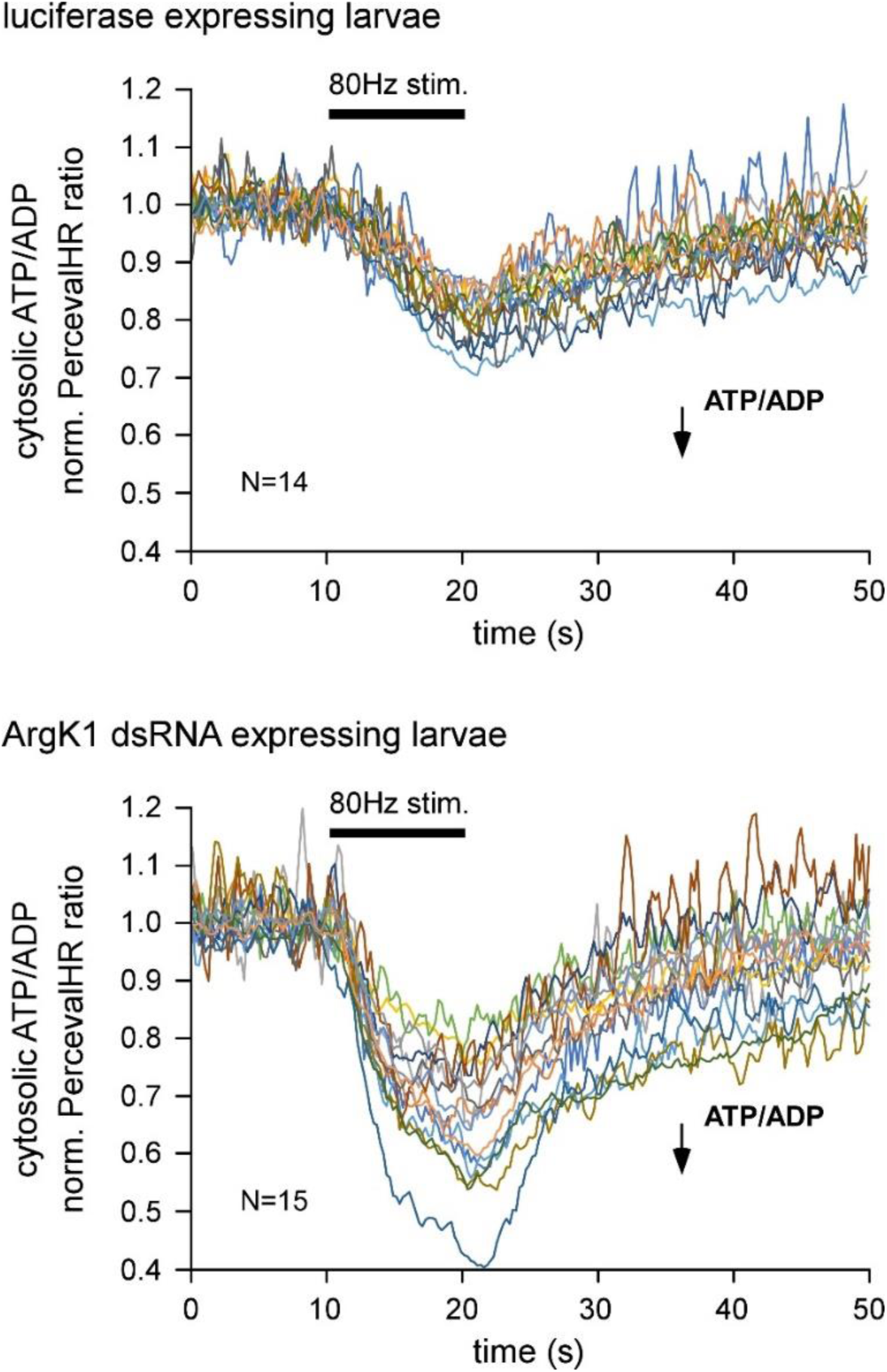
Plots of Stimulation-Induced Changes in Presynaptic PercevalHR Fluorescence From Individual Experiments. Normalized PercevalHR ratios (483nm ex./436nm ex. with 520 nm em.) plotted against time showing a decrease in ATP/ADP in response to a 10 second episode of nerve stimulation. Each trace represent a single trial on the terminal of MN13-Ib in segment 4 on a different larva. The terminals co-expressed PercevalHR with either ArgK1 dsRNA (top panel; N=14) or luciferase (bottom panel; N=15). Each trace was smoothed with a 3-point moving average. Terminals were examined in HL6 (2mM CaCl_2_, 7mM L-glutamate).

**Figure S7.**
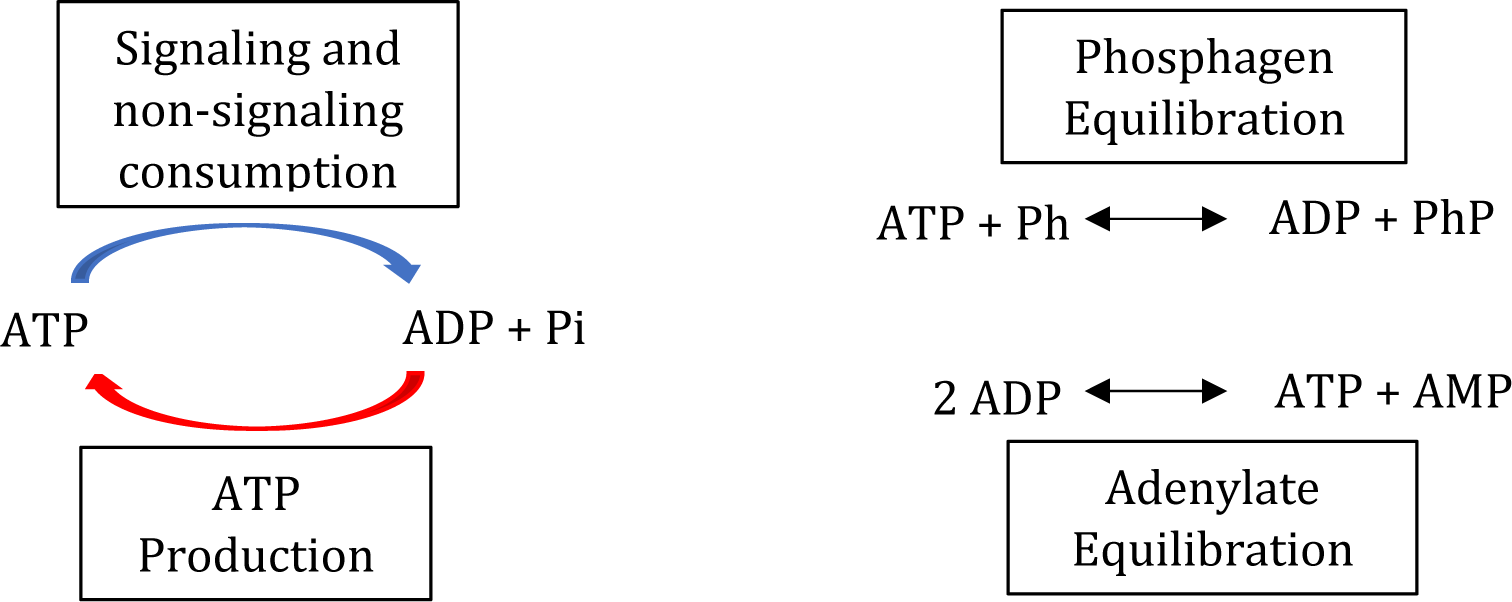
Computational Model Summary Diagram. The model tracks the concentrations of ATP, ADP, AMP, inorganic phosphate (P_i_), guanidine acceptor (Ph, either arginine or creatine), and phosphagen (PhP, either arginine phosphate or creatine phosphate). Four processes modify those concentrations: ATP consumption with consumption rate c, ATP production with production rate p, phosphagen equilibration with equilibrium constant K_Ph_, and adenylate equilibration with equilibrium constant K_Ad_.

**Figure S8.**
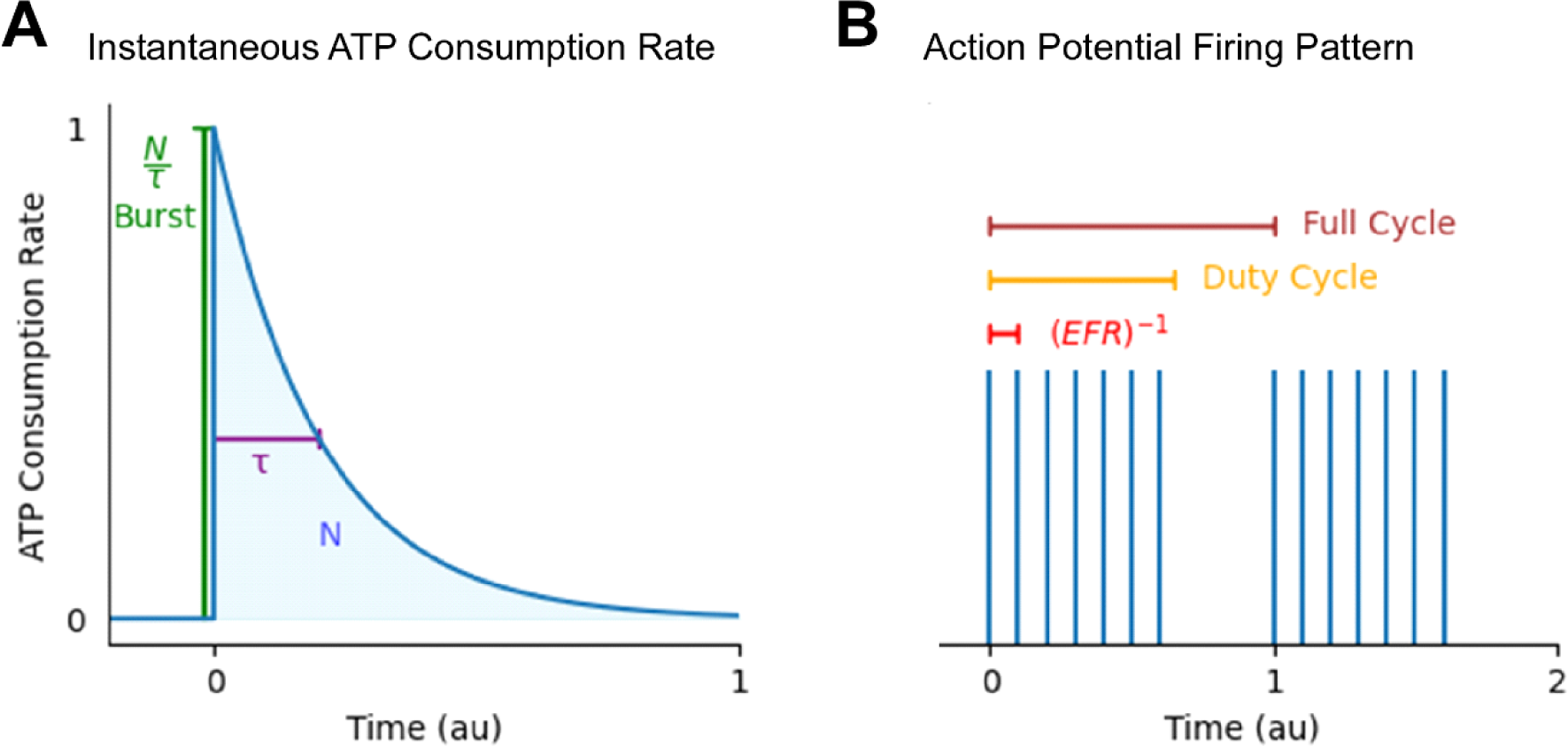
ATP Consumption Rate for Each Signaling Process. (A) Instantaneous ATP consumption rate of a single signaling process from a single AP as a function of the time passed since the AP. N is the number of ATP molecules consumed over the process’ entire lifetime, equal to the area under the curve (shaded blue). τ is the process’ decay time (purple). The instantaneous ATP consumption rate at the onset of the process is N/ τ (dark green). (B) Example AP firing pattern of a drosophila motor neuron. The pattern repeats after a Full Cycle corresponding to one body wall contraction. Each cycle consists of a burst of APs followed by a pause. The Duty Cycle is the ratio of the burst duration to the Full Cycle. The Endogenous Firing Rate (EFR) is the number of AP per second during a burst.

**Figure S9.**
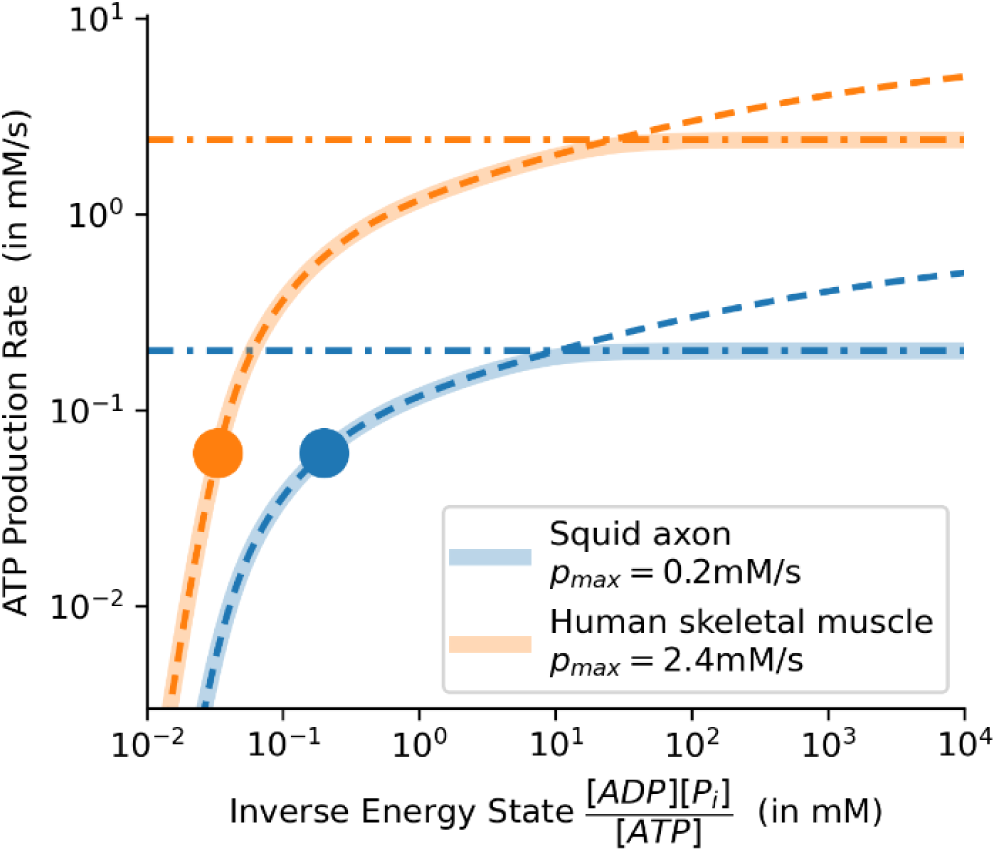
ATP Production Rate as a Function of the Inverse Energy State for Two Different Organisms. A plot of the ATP production rate versus changing inverse energy state. Rest states are shown as solid circles. Their inverse energy states are computed using ATP, ADP, and P_i_ concentrations from the literature. Their ATP production rates are equal to the rest ATP consumption rate in the synaptic terminal of interest (in this case, MN6/7-Ib on muscle fiber #6). The production limits are shown as dash-dotted lines. The dashed lines correspond to uncapped Wilson models, adjusted to pass through the rest states but not subject to the production limit.

**Table S1.** Terminal Specific Parameters Related to Energy Demand and its Volatility. Data in rows 1-3 sourced from Extended Data Table 5-1 of (Justs et al., 2022). Standard error of the mean calculated using propagation of uncertainty principles.

|  | Motor Neuron Terminal Identity |  |  |  |  |  |
| --- | --- | --- | --- | --- | --- | --- |
|  | MN6-lb | MN6-ls | MN13-lb | MN13-ls | MN12-lb | MN12-ls |
| **Sig. Energy Demand**<br>**(10<sup>6</sup> ATP molecules/s)** | 155.8<br>±7.3 | 9.2<br>±0.8 | 244.3<br>±16.6 | 8.0<br>±0.7 | 179.8<br>±9.2 | 11.3<br>±1.0 |
| **Non-Sig. Energy Demand**<br>**(10<sup>6</sup> ATP molecules/s)** | 11.3<br>±0.7 | 3.4<br>±0.7 | 14.0<br>±1.6 | 3.6<br>±0.5 | 13.3<br>±1.3 | 4.8<br>±1.0 |
| **Total Energy Demand**<br>**(10<sup>6</sup> ATP molecules/s)** | 167.1<br>±7.3 | 12.6<br>±1.1 | 258.3<br>±16.7 | 11.6<br>±0.9 | 193.1<br>±9.3 | 16.0<br>±1.4 |
| **Energy Demand Volatility**<br>**(Total/Non-Sig. Demand)** | 14.7<br>±0.70 | 3.7<br>±0.46 | 18.4<br>±1.44 | 3.2<br>±0.31 | 14.6<br>±0.83 | 3.4<br>±0.43 |

**Table S2.** Terminal Specific Parameters Related to Mitochondrial and Terminal Morphology. Data sourced from Extended Data Tables 1-1 and 6-1 of (Justs et al., 2022). Standard error of the mean (SEM) show for each. Propagation of uncertainty theory used to calculate SEM for total mitochondrial volume.

|  | Motor Neuron Terminal Identity |  |  |  |  |  |
| --- | --- | --- | --- | --- | --- | --- |
|  | MN6-lb | MN6-ls | MN13-lb | MN13-ls | MN12-lb | MN12-ls |
| **Total Mito Vol (μm<sup>3</sup>)** | 19.52<br>±1.29 | 3.69<br>±0.35 | 25.13<br>±1.92 | 2.98<br>±0.60 | 22.32<br>±1.51 | 5.5<br>±0.58 |
| **Terminal Volume (μm<sup>3</sup>)** | 310.5<br>±20.0 | 90.1<br>±20.1 | 391.5<br>±48.8 | 96.7<br>±15.6 | 366.4<br>±37.7 | 128.2<br>±28.6 |
| **Mito Vol Density (%)** | 6.29<br>±0.41 | 4.10<br>±0.39 | 6.42<br>±0.49 | 3.08<br>±0.62 | 6.09<br>±0.41 | 4.29<br>±0.45 |

**Table S3.** Terminal Specific Parameters Related to Power Demand and its Volatility. Estimates of total power demand for each terminal during, the 1^st^, 10^th^ and 120^th^ second after the start of peristaltic locomotion. Estimates of power demand volatility were generated by dividing the total power demand by the non-signaling power demand.

|  | Motor Neuron Terminal Identity |  |  |  |  |  |
| --- | --- | --- | --- | --- | --- | --- |
|  | MN6-Ib | MN6-Is | MN13-Ib | MN13-Is | MN12-Ib | MN12-Is |
| **1<sup>st</sup> Second of Activity**<br>**Total Power Demand (mM/s)** | 0.45 | 0.13 | 0.50 | 0.12 | 0.46 | 0.13 |
| **1<sup>st</sup> Sec Pwr Demand Volatility**<br>**(Total/Non-Sig. Power Demand)** | 7.5 | 2.2 | 8.4 | 2.0 | 7.7 | 2.2 |
| **10<sup>th</sup> Second of Activity**<br>**Total Power Demand (mM/s)** | 0.54 | 0.16 | 0.61 | 0.14 | 0.54 | 0.15 |
| **10<sup>th</sup> Sec Pwr Demand Volatility**<br>**(Total/Non-Sig. Power Demand)** | 9.0 | 2.7 | 10.2 | 2.2 | 9.0 | 2.5 |
| **120<sup>th</sup> Second of Activity**<br>**Total Power Demand (mM/s)**<br>**(Power Demand Volatility)** | 0.87 | 0.23 | 1.06 | 0.20 | 0.85 | 0.21 |
| **120<sup>th</sup> Sec Pwr Demand Volatility**<br>**(Total/Non-Sig. Power Demand)** | 14.5 | 3.8 | 17.6 | 3.3 | 14.2 | 3.5 |

**Table S4.** Phosphagen System Values for Different Organisms and Tissues. Concentrations are in mM. Temperatures are in °C. **a.** (Zhu et al., 2012). **b**. (Dechent et al., 1999). **c**. (Lawson and Veech, 1979). **d**. (Sharma et al., 2020). **e**. (Ren and Hultman, 1989). **f**. (Ren and Hultman, 1989). **g**. (Flouris et al., 2015). **h**. (Caldwell, 1960). **i**. (Requena et al., 1979). **j**. (Mullins and Brinley, 1967). **k**. (Deffner and Hafter, 1959). **l**. (Deffner, 1961). **m**. (DiPolo and Beauge, 1996). **n**. (Teague and Dobson, 1999). **o**. (Jacobson, 2005).

| Organism / Tissue | [ATP] | [Ph] ; Phos | [PhP] ; Phos | [P<sub>i</sub>] | K<sub>Ph</sub> | Temp |
| --- | --- | --- | --- | --- | --- | --- |
| Human Grey Matter | 3<sup>a</sup> | 3.4<sup>b</sup> ; Creatine | 3<sup>a</sup> ; Creatine | 0.93<sup>a</sup> | 166<sup>c</sup> | 37.2<sup>d</sup> |
| Human Skeletal Muscle | 6<sup>e</sup> | 12.3<sup>f</sup> ; Creatine | 18.9<sup>f</sup> ; Creatine | 8.48<sup>f</sup> | 166<sup>c</sup> | 35.85<sup>g</sup> |
| Squid Axoplasm | 2.16<sup>h,i,j</sup> | 3.33<sup>k,l</sup> ; Arginine | 7.5<sup>m</sup> ; Arginine | 17.8<sup>m</sup> | 39.64<sup>n</sup> | 15° |
| Squid Axoplasm<br>(No phosphagen) | 2.16<sup>h,i,j</sup> | — | — | 17.8<sup>m</sup> | — | 15° |

**Table S5.**
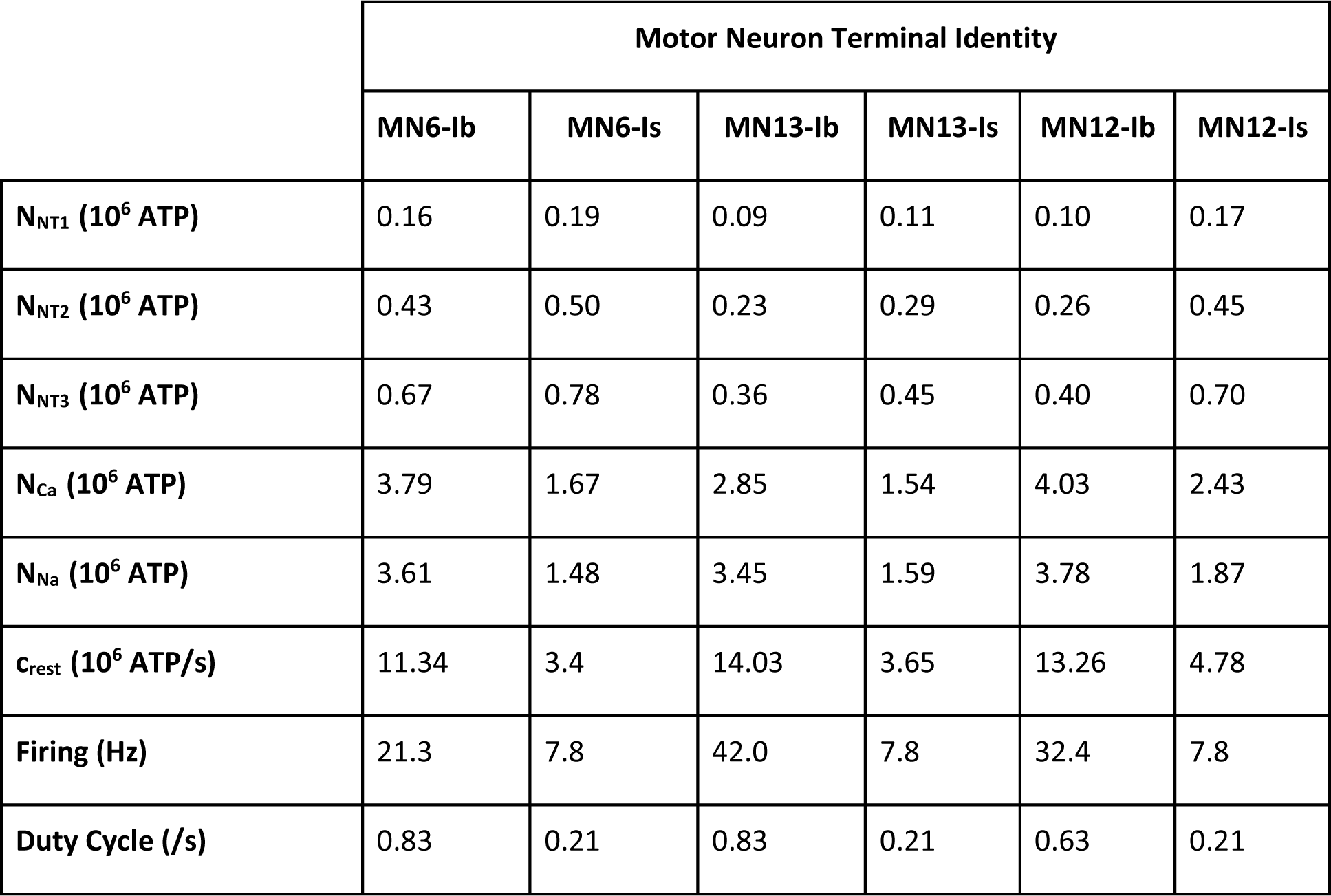
Terminal Specific Parameters Related to ATP Consumption. Duty Cycle is the proportion of a second. Data sourced from Extended Data Tables of (Justs et al., 2022).

**Table S6.** Time Courses of Energy-Consuming Signaling Processes. Values of τ (seconds) for each signaling process used in the construction of ATP consumption rate c (Justs et al., 2022).

|  | Energy Consuming Signaling Processes |  |  |  |  |
| --- | --- | --- | --- | --- | --- |
|  | Ca<sup>2+</sup> | Na<sup>+</sup> | NT1 | NT2 | NT3 |
| **Time course (τ)** | 0.0389 | 63.5 | 0.175 | 7.21 | 20.0 |

Supplementary File S1. Simulation code.

Zip file containing the jupyter notebook used to perform simulations and create related figures: https://drive.google.com/file/d/10HvcA8YV0xOv2M0knVbTayeynguU09le/view?usp=share_link

## References

Aberle H, Haghighi AP, Fetter RD, McCabe BD, Magalhaes TR, Goodman CS (2002) wishful thinking encodes a BMP type II receptor that regulates synaptic growth in Drosophila. Neuron 33:545–558.

Andersen P, Saltin B (1985) Maximal perfusion of skeletal muscle in man. J Physiol 366:233–249.

Andres RH, Ducray AD, Schlattner U, Wallimann T, Widmer HR (2008) Functions and effects of creatine in the central nervous system. Brain Res Bull 76:329–343.

Aponte-Santiago NA, Ormerod KG, Akbergenova Y, Littleton JT (2020) Synaptic Plasticity Induced by Differential Manipulation of Tonic and Phasic Motoneurons in Drosophila. J Neurosci 40:6270–6288.

Balaban RS (2009) Domestication of the cardiac mitochondrion for energy conversion. J Mol Cell Cardiol 46:832–841.

Bangsbo J, Gollnick PD, Graham TE, Juel C, Kiens B, Mizuno M, Saltin B (1990) Anaerobic energy production and O2 deficit-debt relationship during exhaustive exercise in humans. J Physiol 422:539–559.

Barber TW, Brockway JA, Higgins LS (1970) The density of tissues in and about the head. Acta Neurol Scand 46:85–92.

Barclay CJ (2017) Energy demand and supply in human skeletal muscle. J Muscle Res Cell Motil 38:143–155.

Berrigan D, Pepin DJ (1995) How Maggots Move: Allometry and Kinematics of Crawling in Larval Diptera. Journal of Insect Physiology 41:329–337.

Bertoni-Freddari C, Fattoretti P, Giorgetti B, Grossi Y, Balietti M, Casoli T, Di Stefano G, Perretta G (2007) Preservation of mitochondrial volume homeostasis at the early stages of age-related synaptic deterioration. Ann N Y Acad Sci 1096:138–146.

Blomstrand E, Radegran G, Saltin B (1997) Maximum rate of oxygen uptake by human skeletal muscle in relation to maximal activities of enzymes in the Krebs cycle. J Physiol-London 501:455–460.

Buszczak M, Paterno S, Lighthouse D, Bachman J, Planck J, Owen S, Skora AD, Nystul TG, Ohlstein B, Allen A, Wilhelm JE, Murphy TD, Levis RW, Matunis E, Srivali N, Hoskins RA, Spradling AC (2007) The carnegie protein trap library: a versatile tool for Drosophila developmental studies. Genetics 175:1505–1531.

Cable MB, Briggs FN (1988) Allosteric regulation of cardiac sarcoplasmic reticulum Ca-ATPase: a comparative study. Mol Cell Biochem 82:29–36.

Caldwell PC (1960) The phosphorus metabolism of squid axons and its relationship to the active transport of sodium. J Physiol 152:545–560.

Castro J, Ruminot I, Porras OH, Flores CM, Hermosilla T, Verdugo E, Venegas F, Hartel S, Michea L, Barros LF (2006) ATP steal between cation pumps: a mechanism linking Na+ influx to the onset of necrotic Ca2+ overload. Cell Death Differ 13:1675–1685.

Chanaday NL, Kavalali ET (2018) Optical detection of three modes of endocytosis at hippocampal synapses. Elife 7.

Chance B, Leigh JS, Jr., Kent J, McCully K, Nioka S, Clark BJ, Maris JM, Graham T (1986) Multiple controls of oxidative metabolism in living tissues as studied by phosphorus magnetic resonance. Proc Natl Acad Sci U S A 83:9458–9462.

Chouhan AK, Zhang J, Zinsmaier KE, Macleod GT (2010) Presynaptic mitochondria in functionally different motor neurons exhibit similar affinities for Ca2+ but exert little influence as Ca2+ buffers at nerve firing rates in situ. J Neurosci 30:1869–1881.

Chouhan AK, Ivannikov MV, Lu Z, Sugimori M, Llinas RR, Macleod GT (2012) Cytosolic calcium coordinates mitochondrial energy metabolism with presynaptic activity. J Neurosci 32:1233–1243.

Clarke RJ (2009) Mechanism of allosteric effects of ATP on the kinetics of P-type ATPases. Eur Biophys J 39:3–17.

Dechent P, Pouwels PJ, Wilken B, Hanefeld F, Frahm J (1999) Increase of total creatine in human brain after oral supplementation of creatine-monohydrate. Am J Physiol 277:R698–704.

Deffner GG (1961) The dialyzable free organic constituents of squid blood; a comparison with nerve axoplasm. Biochim Biophys Acta 47:378–388.

Deffner GG, Hafter RE (1959) Chemical investigations of the giant nerve fibers of the squid. I. Fractionation of dialyzable constitutents of axoplasm and quantitative determination of the free amino acids. Biochim Biophys Acta 32:362–374.

DiPolo R, Beauge L (1996) In squid axons phosphoarginine plays a key role in modulating Na-Ca exchange fluxes at micromolar [Ca2+]i. Ann N Y Acad Sci 779:199–207.

Dzeja PP, Hoyer K, Tian R, Zhang S, Nemutlu E, Spindler M, Ingwall JS (2011) Rearrangement of energetic and substrate utilization networks compensate for chronic myocardial creatine kinase deficiency. J Physiol 589:5193–5211.

Ellington WR (1989) Phosphocreatine represents a thermodynamic and functional improvement over other muscle phosphagens. J Exp Biol 143:177–194.

Engl E, Jolivet R, Hall CN, Attwell D (2017) Non-signalling energy use in the developing rat brain. J Cereb Blood Flow Metab 37:951–966.

Erecinska M, Deas J, Silver IA (1995) The effect of pH on glycolysis and phosphofructokinase activity in cultured cells and synaptosomes. J Neurochem 65:2765–2772.

Farrance I, Frenkel R (2012) Uncertainty of Measurement: A Review of the Rules for Calculating Uncertainty Components through Functional Relationships. Clin Biochem Rev 33:49–75.

Ferguson BS, Rogatzki MJ, Goodwin ML, Kane DA, Rightmire Z, Gladden LB (2018) Lactate metabolism: historical context, prior misinterpretations, and current understanding. Eur J Appl Physiol 118:691–728.

Filippin L, Abad MC, Gastaldello S, Magalhaes PJ, Sandona D, Pozzan T (2005) Improved strategies for the delivery of GFP-based Ca2+ sensors into the mitochondrial matrix. Cell calcium 37:129–136.

Flouris AD, Webb P, Kenny GP (2015) Noninvasive assessment of muscle temperature during rest, exercise, and postexercise recovery in different environments. J Appl Physiol (1985) 118:1310–1320.

Frey D, Schneider C, Xu L, Borg J, Spooren W, Caroni P (2000) Early and selective loss of neuromuscular synapse subtypes with low sprouting competence in motoneuron diseases. J Neurosci 20:2534–2542.

Gan Q, Watanabe S (2018) Synaptic Vesicle Endocytosis in Different Model Systems. Front Cell Neurosci 12:171.

Gimber N, Tadeus G, Maritzen T, Schmoranzer J, Haucke V (2015) Diffusional spread and confinement of newly exocytosed synaptic vesicle proteins. Nat Commun 6:8392.

Glancy B, Balaban RS (2012) Role of mitochondrial Ca2+ in the regulation of cellular energetics. Biochemistry 51:2959–2973.

Greenhaff PL, Timmons JA (1998) Interaction between aerobic and anaerobic metabolism during intense muscle contraction. Exerc Sport Sci Rev 26:1–30.

He K, Han Y, Li X, Dickman D (2022) Physiologic and nanoscale distinctions that define glutmatergic synapses in tonic vs phasic neurons. BioRxiv.

Hoang B, Chiba A (2001) Single-cell analysis of Drosophila larval neuromuscular synapses. Dev Biol 229:55–70.

Hoppeler H (1990) The different relationship of VO2max to muscle mitochondria in humans and quadrupedal animals. Respir Physiol 80:137–145.

Howald H, Hoppeler H, Claassen H, Mathieu O, Straub R (1985) Influences of endurance training on the ultrastructural composition of the different muscle fiber types in humans. Pflugers Arch 403:369–376.

Ivannikov MV, Macleod GT (2013) Mitochondrial free Ca(2)(+) levels and their effects on energy metabolism in Drosophila motor nerve terminals. Biophys J 104:2353–2361.

Jacobson LD (2005) Essential fish habitat source document, Longfin inshore squid, Loligo pealeii, life history and habitat characteristics. In: NOAA technical memorandum NMFS-NE ; 193 (Commerce USDo, ed). Woods Hole, Massachusetts: National Oceanic and Atmospheric Administration.

Johnson MA, Polgar J, Weightman D, Appleton D (1973) Data on the distribution of fibre types in thirty-six human muscles. An autopsy study. J Neurol Sci 18:111–129.

Johri A, Beal MF (2012) Mitochondrial dysfunction in neurodegenerative diseases. J Pharmacol Exp Ther 342:619–630.

Jorgensen LB, Overgaard J, Hunter-Manseau F, Pichaud N (2021) Dramatic changes in mitochondrial substrate use at critically high temperatures: a comparative study using Drosophila. J Exp Biol 224.

Justs KA, Lu Z, Chouhan AK, Borycz JA, Lu Z, Meinertzhagen IA, Macleod GT (2022) Presynaptic Mitochondrial Volume and Packing Density Scale with Presynaptic Power Demand. J Neurosci 42:954–967.

Kaasik A, Veksler V, Boehm E, Novotova M, Ventura-Clapier R (2003) From energy store to energy flux: a study in creatine kinase-deficient fast skeletal muscle. FASEB J 17:708–710.

Kanning KC, Kaplan A, Henderson CE (2010) Motor neuron diversity in development and disease. Annu Rev Neurosci 33:409–440.

Kononenko NL, Haucke V (2015) Molecular mechanisms of presynaptic membrane retrieval and synaptic vesicle reformation. Neuron 85:484–496.

Korzeniewski B (2007) Regulation of oxidative phosphorylation through parallel activation. Biophys Chem 129:93–110.

Koveal D, Diaz-Garcia CM, Yellen G (2020) Fluorescent Biosensors for Neuronal Metabolism and the Challenges of Quantitation. Curr Opin Neurobiol 63:111–121.

Kurdyak P, Atwood HL, Stewart BA, Wu CF (1994) Differential physiology and morphology of motor axons to ventral longitudinal muscles in larval Drosophila. J Comp Neurol 350:463–472.

Lang AB, Wyss C, Eppenberger HM (1980) Localization of arginine kinase in muscles fibres of Drosophila melanogaster. J Muscle Res Cell Motil 1:147–161.

Lawson JW, Veech RL (1979) Effects of pH and free Mg2+ on the Keq of the creatine kinase reaction and other phosphate hydrolyses and phosphate transfer reactions. J Biol Chem 254:6528–6537.

Leys C, Ley C, Klein O, Bernard B, Licata L (2013) Detecting outliers: Do not use standard deviation around the mean, use absolute deviation around the median. Journal of Experimental Social Psychology 49:764–766.

Li X, Chien C, Han Y, Sun Z, Chen X, Dickman D (2021) Autocrine inhibition by a glutamate-gated chloride channel mediates presynaptic homeostatic depression. Sci Adv 7:eabj1215.

Lipton P, Whittingham TS (1982) Reduced ATP concentration as a basis for synaptic transmission failure during hypoxia in the in vitro guinea-pig hippocampus. J Physiol 325:51–65.

Llorente-Folch I, Rueda CB, Amigo I, del Arco A, Saheki T, Pardo B, Satrustegui J (2013) Calcium-regulation of mitochondrial respiration maintains ATP homeostasis and requires ARALAR/AGC1-malate aspartate shuttle in intact cortical neurons. J Neurosci 33:13957–13971, 13971a.

Lnenicka GA, Keshishian H (2000) Identified motor terminals in Drosophila larvae show distinct differences in morphology and physiology. J Neurobiol 43:186–197.

Lowe MT, Kim EH, Faull RL, Christie DL, Waldvogel HJ (2013) Dissociated expression of mitochondrial and cytosolic creatine kinases in the human brain: a new perspective on the role of creatine in brain energy metabolism. J Cereb Blood Flow Metab 33:1295–1306.

Lu Z, Chouhan AK, Borycz JA, Lu Z, Rossano AJ, Brain KL, Zhou Y, Meinertzhagen IA, Macleod GT (2016) High-Probability Neurotransmitter Release Sites Represent an Energy-Efficient Design. Curr Biol 26:2562–2571.

Macleod GT (2012) Imaging and analysis of nonratiometric calcium indicators at the Drosophila larval neuromuscular junction. Cold Spring Harb Protoc 2012:802–809.

Macleod GT, Ivannikov MV (2017) Examining Mitochondrial Function at Synapses In Situ. In: Techniques to Investigate Mitochondrial Function in Neurons, Neuromethods (Strack S, Usachev YM, eds), pp 279–197: Springer Science+Business Media LLC.

Macleod GT, Hegstrom-Wojtowicz M, Charlton MP, Atwood HL (2002) Fast calcium signals in Drosophila motor neuron terminals. J Neurophysiol 88:2659–2663.

Macleod GT, Marin L, Charlton MP, Atwood HL (2004) Synaptic vesicles: test for a role in presynaptic calcium regulation. J Neurosci 24:2496–2505.

Malthankar-Phatak GH, Patel AB, Xia Y, Hong S, Chowdhury GM, Behar KL, Orina IA, Lai JC (2008) Effects of continuous hypoxia on energy metabolism in cultured cerebro-cortical neurons. Brain Res 1229:147–154.

Mank M, Santos AF, Direnberger S, Mrsic-Flogel TD, Hofer SB, Stein V, Hendel T, Reiff DF, Levelt C, Borst A, Bonhoeffer T, Hubener M, Griesbeck O (2008) A genetically encoded calcium indicator for chronic in vivo two-photon imaging. Nat Methods 5:805–811.

Menail HA, Cormier SB, Ben Youssef M, Jorgensen LB, Vickruck JL, Morin P, Jr., Boudreau LH, Pichaud N (2022) Flexible Thermal Sensitivity of Mitochondrial Oxygen Consumption and Substrate Oxidation in Flying Insect Species. Front Physiol 13:897174.

Mondragao MA, Schmidt H, Kleinhans C, Langer J, Kafitz KW, Rose CR (2016) Extrusion versus diffusion: mechanisms for recovery from sodium loads in mouse CA1 pyramidal neurons. J Physiol 594:5507–5527.

Mookerjee SA, Gerencser AA, Nicholls DG, Brand MD (2017) Quantifying intracellular rates of glycolytic and oxidative ATP production and consumption using extracellular flux measurements. J Biol Chem 292:7189–7207.

Mullins LJ, Brinley FJ, Jr. (1967) Some factors influencing sodium extrusion by internally dialyzed squid axons. J Gen Physiol 50:2333–2355.

Neville KE, Bosse TL, Klekos M, Mills JF, Weicksel SE, Waters JS, Tipping M (2018) A novel ex vivo method for measuring whole brain metabolism in model systems. J Neurosci Methods 296:32–43.

Newman ZL, Hoagland A, Aghi K, Worden K, Levy SL, Son JH, Lee LP, Isacoff EY (2017) Input-Specific Plasticity and Homeostasis at the Drosophila Larval Neuromuscular Junction. Neuron 93:1388–1404 e1310.

Nicholls DG, Ferguson SJ (2002) Bioenergetics 3, Third Edition. London: Academic.

NRC (2011) Guide for the Care and Use of Laboratory Animals. In, Eighth Edition (Institute for Laboratory Animal Research NRCotNA, ed). Washington, D.C.: National Academies Press.

Patel A, Malinovska L, Saha S, Wang J, Alberti S, Krishnan Y, Hyman AA (2017) ATP as a biological hydrotrope. Science 356:753–756.

Perkins G, Hsiao YH, Yin S, Tjong J, Tran MT, Lau J, Xue J, Liu S, Ellisman MH, Zhou D (2012) Ultrastructural modifications in the mitochondria of hypoxia-adapted Drosophila melanogaster. PLoS One 7:e45344.

Perkins GA, Jackson DR, Spirou GA (2015) Resolving presynaptic structure by electron tomography. Synapse 69:268–282.

Poburko D, Santo-Domingo J, Demaurex N (2011) Dynamic regulation of the mitochondrial proton gradient during cytosolic calcium elevations. J Biol Chem 286:11672–11684.

Pun S, Santos AF, Saxena S, Xu L, Caroni P (2006) Selective vulnerability and pruning of phasic motoneuron axons in motoneuron disease alleviated by CNTF. Nat Neurosci 9:408–419.

Rangaraju V, Calloway N, Ryan TA (2014) Activity-driven local ATP synthesis is required for synaptic function. Cell 156:825–835.

Rauskolb C, Sun S, Sun G, Pan Y, Irvine KD (2014) Cytoskeletal tension inhibits Hippo signaling through an Ajuba-Warts complex. Cell 158:143–156.

Ren JM, Hultman E (1989) Regulation of glycogenolysis in human skeletal muscle. J Appl Physiol (1985) 67:2243–2248.

Requena J, Mullins LJ, Brinley FJ, Jr. (1979) Calcium content and net fluxes in squid giant axons. J Gen Physiol 73:327–342.

Rossano AJ, Macleod GT (2007) Loading Drosophila nerve terminals with calcium indicators. J Vis Exp:250.

Rossano AJ, Chouhan AK, Macleod GT (2013) Genetically encoded pH-indicators reveal activity-dependent cytosolic acidification of Drosophila motor nerve termini in vivo. J Physiol 591:1691–1706.

Rossano AJ, Kato A, Minard KI, Romero MF, Macleod GT (2017) Na(+) /H(+) exchange via the Drosophila vesicular glutamate transporter mediates activity-induced acid efflux from presynaptic terminals. J Physiol 595:805–824.

Sankaranarayanan S, De Angelis D, Rothman JE, Ryan TA (2000) The use of pHluorins for optical measurements of presynaptic activity. Biophys J 79:2199–2208.

Saupe KW, Spindler M, Tian R, Ingwall JS (1998) Impaired cardiac energetics in mice lacking muscle-specific isoenzymes of creatine kinase. Circ Res 82:898–907.

Saupe KW, Spindler M, Hopkins JC, Shen W, Ingwall JS (2000) Kinetic, thermodynamic, and developmental consequences of deleting creatine kinase isoenzymes from the heart. Reaction kinetics of the creatine kinase isoenzymes in the intact heart. J Biol Chem 275:19742–19746.

Schiaffino S, Reggiani C (2011) Fiber types in mammalian skeletal muscles. Physiol Rev 91:1447–1531.

Schlattner U, Tokarska-Schlattner M, Wallimann T (2006) Mitochondrial creatine kinase in human health and disease. Biochim Biophys Acta 1762:164–180.

Segal SS, White TP, Faulkner JA (1986) Architecture, composition, and contractile properties of rat soleus muscle grafts. Am J Physiol 250:C474–479.

Sharma AA, Nenert R, Mueller C, Maudsley AA, Younger JW, Szaflarski JP (2020) Repeatability and Reproducibility of in-vivo Brain Temperature Measurements. Front Hum Neurosci 14:598435.

Shcherbo D, Merzlyak EM, Chepurnykh TV, Fradkov AF, Ermakova GV, Solovieva EA, Lukyanov KA, Bogdanova EA, Zaraisky AG, Lukyanov S, Chudakov DM (2007) Bright far-red fluorescent protein for whole-body imaging. Nat Methods 4:741–746.

Soykan T, Maritzen T, Haucke V (2016) Modes and mechanisms of synaptic vesicle recycling. Curr Opin Neurobiol 39:17–23.

St-Pierre J, Lin J, Krauss S, Tarr PT, Yang R, Newgard CB, Spiegelman BM (2003) Bioenergetic analysis of peroxisome proliferator-activated receptor gamma coactivators 1alpha and 1beta (PGC-1alpha and PGC-1beta) in muscle cells. J Biol Chem 278:26597–26603.

Steinert JR, Kuromi H, Hellwig A, Knirr M, Wyatt AW, Kidokoro Y, Schuster CM (2006) Experience-dependent formation and recruitment of large vesicles from reserve pool. Neuron 50:723–733.

Tantama M, Martinez-Francois JR, Mongeon R, Yellen G (2013) Imaging energy status in live cells with a fluorescent biosensor of the intracellular ATP-to-ADP ratio. Nat Commun 4:2550.

Teague WE, Jr., Dobson GP (1999) Thermodynamics of the arginine kinase reaction. J Biol Chem 274:22459–22463.

Tonkonogi M, Sahlin K (1997) Rate of oxidative phosphorylation in isolated mitochondria from human skeletal muscle: effect of training status. Acta Physiol Scand 161:345–353.

van Deursen J, Heerschap A, Oerlemans F, Ruitenbeek W, Jap P, ter Laak H, Wieringa B (1993) Skeletal muscles of mice deficient in muscle creatine kinase lack burst activity. Cell 74:621–631.

Vandoorne T, De Bock K, Van Den Bosch L (2018) Energy metabolism in ALS: an underappreciated opportunity? Acta Neuropathol 135:489–509.

Wallimann T, Wyss M, Brdiczka D, Nicolay K, Eppenberger HM (1992) Intracellular compartmentation, structure and function of creatine kinase isoenzymes in tissues with high and fluctuating energy demands: the ‘phosphocreatine circuit’ for cellular energy homeostasis. Biochem J 281 (Pt 1):21–40.

Walter G, Vandenborne K, Elliott M, Leigh JS (1999) In vivo ATP synthesis rates in single human muscles during high intensity exercise. J Physiol 519 Pt 3:901–910.

Wang Y, Lobb-Rabe M, Ashley J, Anand V, Carrillo RA (2021) Structural and Functional Synaptic Plasticity Induced by Convergent Synapse Loss in the Drosophila Neuromuscular Circuit. J Neurosci 41:1401–1417.

Watanabe S, Liu Q, Davis MW, Hollopeter G, Thomas N, Jorgensen NB, Jorgensen EM (2013) Ultrafast endocytosis at Caenorhabditis elegans neuromuscular junctions. Elife 2:e00723.

Wilson DF (2017a) Oxidative phosphorylation: regulation and role in cellular and tissue metabolism. J Physiol 595:7023–7038.

Wilson DF (2017b) Oxidative phosphorylation: unique regulatory mechanism and role in metabolic homeostasis. J Appl Physiol (1985) 122:611–619.

Wong CO, Chen K, Lin YQ, Chao Y, Duraine L, Lu Z, Yoon WH, Sullivan JM, Broadhead GT, Sumner CJ, Lloyd TE, Macleod GT, Bellen HJ, Venkatachalam K (2014) A TRPV channel in Drosophila motor neurons regulates presynaptic resting Ca2+ levels, synapse growth, and synaptic transmission. Neuron 84:764–777.

Wright M, Kim A, Son YJ (2011) Subcutaneous administration of muscarinic antagonists and triple-immunostaining of the levator auris longus muscle in mice. J Vis Exp.

Yellen G (2018) Fueling thought: Management of glycolysis and oxidative phosphorylation in neuronal metabolism. J Cell Biol 217:2235–2246.

Zhu H, Sun X (2013) Three-dimensional structure of axonal mitochondria reflects the age of drosophila. Neural Regen Res 8:616–621.

Zhu XH, Qiao H, Du F, Xiong Q, Liu X, Zhang X, Ugurbil K, Chen W (2012) Quantitative imaging of energy expenditure in human brain. Neuroimage 60:2107–2117.

