## Supplementary File S1 for "Mitochondrial phosphagen kinases support the volatile power demands of motor nerve terminals": perform-simulations.ipynb

```
# %% [markdown]
## Prepare simulations

# %% [code]
import os, sys, pickle, warnings
from pathlib import Path
import numpy as np
import scipy as sp
from scipy.integrate import cumtrapz
from scipy.constants import Avogadro, R
import matplotlib.pyplot as plt
from matplotlib.colors import hsv_to_rgb, rgb_to_hsv, to_rgb
from matplotlib.patches import FancyArrowPatch, ArrowStyle
from datetime import datetime
from tqdm import tqdm

cm = 1/2.54 # Inch-to-cm conversion: 1*cm is 1cm in inches.

from setup_simulation import *
terminal_data = load_terminal_data('terminal-data.npz')
terminals     = sorted(terminal_data.keys())[::-1]
terminals     = terminals[1::2]+terminals[::2]
print(terminals)

# %% [markdown]
## Make manuscript figures

- Figure 3:
  10 seconds of activity, simulate from t=-1 to t=20.
  All 6 terminals.
  Models: rest production, and Wilson wo/ phosphagen.
  
- Figure 6:
  10 seconds of activity, simulate from t=-1 to t=40.
  Terminals 6b and 6s.
  Models: Wilson wo/ phosphagen, Wilson+creatine, Wilson+arginine.

- Figure 7:
  sustained activity, simulate from t=-1 to t=120.
  Terminals 6b and 6s.
  Models: Wilson wo/ phosphagen, Wilson+creatine, Wilson+arginine.

# %% [code]
# ---------- Output options ----------

data_dir = 'figure-data'
if not os.path.exists(data_dir):
    os.mkdir(data_dir)

fig_dir = 'figures'
if not os.path.exists(fig_dir):
    os.mkdir(fig_dir)

def data_file(figname):
    fn = ( f'{data_dir}/{figname}_{terminal}_conc={concentration_set}'
           f'_prod={production_model}_rateCapPpc={rate_cap_per_pct_mito}' ) 
    return fn+'.npz'

def savefig(filename):
    fn = f'{fig_dir}/{filename}'
    plt.savefig(fn+'.png', dpi=250)
    plt.savefig(fn+'.eps')

# ---------- Plotting options ----------

plt.rcParams.update({ 
    'savefig.facecolor'    : 'w',
    'lines.linewidth'      : 0.5,
    'lines.linewidth'      : 0.5,
    'font.size'            : 7,
    'legend.frameon'       : False,
    'legend.loc'           : 'upper right',
    'legend.fontsize'      : 7,
    'legend.handlelength'  : 1,
    'legend.labelspacing'  : 0.25, 
    'legend.borderpad'     : 0, 
    'legend.borderaxespad' : 0,
})

colors    = { 'M6Ib':'#009b00', 'M6Is':'#ff0000', 'M13Ib':'#0000ff', 
              'M13Is':'#ffa500', 'M12Ib':'#ff00ff', 'M12Is':'#c8c800' }
zorders   = { term:i for i,term in enumerate(['M12Ib','M12Is','M13Is',
                                              'M6Is','M6Ib','M13Ib'], 1) }
term_opt  = lambda term: dict(color=colors[term], label=term[1:])

panel_opt = dict(x=-0.1, y=1.1, ha='right', va='bottom', 
                 fontdict={'fontsize':10, 'weight':'bold'} )

def create_color_shades(rgb):
    if rgb==colors['M6Is']:
        return np.array(['#F1B2DC', '#fc327e', '#c90704', '#651C32'])
    if rgb==colors['M6Ib']:
        return np.array(['#AADB1E', '#B1BC55', '#009B77', '#4E5b31'])
    rgb = to_rgb(rgb)
    hsv = np.array([rgb_to_hsv(rgb)]*6)
    hsv[:,0] += np.array([-2,-1,0,1,2,3])*0.05      # Slight hue gradient.
    hsv[:,1]  = np.array([0.2,0.4,1,1   ,1 ,1  ])   # Saturation gradient.
    hsv[:,2]  = np.array([0.8,1  ,1,0.8,0.5,0.2])   # Value gradient.
    return np.array(list(map(hsv_to_rgb,hsv)))

# ---------- Simulation wrapper ----------

def run_simulation(settings, fn, recompute_all=False, verbose=True):
    if verbose:
        print(fn)
    if not recompute_all and os.path.exists(fn):
        if verbose:
            print(indent+'[already exists]')
        return False
    t0   = datetime.now()
    globals().update(setup(settings))
    data = integrate(verbose=verbose).T
    if verbose:
        print(indent+'[done in ' +f'{datetime.now()-t0}'[:-4] +'s]')
    np.savez(fn, settings=settings, data=data, times=times, power_consumed=power_consumed)
    return True

# %% [markdown]
### Figure 3

# %% [code]
models = [
    dict(concentration_set='squid axon no arginine', 
         production_model='rest', 
         rate_cap_per_pct_mito=None, 
         model_name='Squid axon concentrations\nRest-level production rate\n'),
    dict(concentration_set='squid axon no arginine', 
         production_model='wilson', 
         rate_cap_per_pct_mito=0.032, 
         model_name='Squid axon concentrations\nWilson production rate\ncapped at 0.032 mM/s/% mito')
    ]

for terminal in terminals:
    for model in models:
        globals().update(model)
        settings  = create_settings(concentration_set, terminal, production_model, 
                                    rate_cap_per_pct_mito=rate_cap_per_pct_mito, 
                                    t_active=10, t_range=[-1,20])
        globals().update(settings)
        fn = data_file('fig3')
        run_simulation(settings, fn, verbose=False)
        print(fn)

# %% [code]
n_row    = 4
n_col    = len(models)
gridspec = dict(hspace=0.55, wspace=0.55, left=0.15, 
                right=0.98, bottom=0.07, top=0.95)
fig,axs  = plt.subplots(n_row, n_col, figsize=(8.5*cm,3.5*n_row*cm), 
                        gridspec_kw=gridspec)

for terminal in terminals:
    for j,model in enumerate(models):
        globals().update(model)
        globals().update(np.load(data_file('fig3'), allow_pickle=True))
        globals().update(settings.item())
        axs[1,j].plot(data[0], data[4], zorder=zorders[terminal], **term_opt(terminal))
        axs[2,j].plot(data[0], data[2], zorder=zorders[terminal], **term_opt(terminal))
        acc = np.gradient(data[2],data[0])
        axs[3,j].plot(data[0], acc, zorder=zorders[terminal], **term_opt(terminal))
    axs[0,0].plot(times, power_consumed, lw=1, zorder=zorders[terminal], **term_opt(terminal))

for i,ax in enumerate(axs.flatten()[[0,1,2,4,6,3,5,7]]):
    ax.spines.right.set_visible(False)
    ax.spines.top.set_visible(False)
    ax.set_xlabel('Time (s)')
    ax.locator_params(nbins=5)
    ax.text(s=chr(65+i), transform=ax.transAxes, **panel_opt)
axs[0,0].set_ylabel('ATP cons. rate (mM/s)')
axs[0,0].legend(bbox_to_anchor=(1.05, 1.05))
for j in range(1,n_col):
    axs[0,j].axis('off')
for j in range(2):
    axs[1,j].set_ylabel('[ATP] (mM)')
#     axs[1,j].set_ylim(0,1.1*At)
    axs[2,j].set_ylabel('ATP prod. rate (mM/s)')
    axs[3,j].set_ylabel('ATP prod. accel. (mM/s/s)')
for i in range(n_row):
    ylim = list(zip(*[ax.get_ylim() for ax in axs[i]]))
    ylim = min(ylim[0]), max(ylim[1])
    ylim = min(0, ylim[0]), ylim[1]
    for ax in axs[i]:
        ax.set_ylim(ylim)

savefig('fig3')
plt.show()

# %% [markdown]
### Figure 6

# %% [code]
models = [
    dict(concentration_set='squid axon no arginine',
         production_model='wilson', 
         rate_cap_per_pct_mito=0.032, 
         model_name='Squid axon, no arginine\nmax prod 0.032 mM/s/%'),
    dict(concentration_set='squid axon',
         production_model='wilson', 
         rate_cap_per_pct_mito=0.032, 
         model_name='Squid axon\nmax prod 0.032 mM/s/%'),
#     dict(concentration_set='dog brain',
#          production_model='wilson', 
#          rate_cap_per_pct_mito=0.025, 
#          model_name='Dog brain concentrations\nmax prod=0.025 mM/s/%\nw/ creatine'),
    dict(concentration_set='human grey matter',
         production_model='wilson', 
         rate_cap_per_pct_mito=0.025, 
         model_name='Human grey matter\nmax prod 0.025 mM/s/%'),
    dict(concentration_set='human skeletal',
         production_model='wilson', 
         rate_cap_per_pct_mito=0.032, 
         model_name='Human skeletal muscle\nmax prod 0.032 mM/s/%')
    ]

# Extra models for supplementary figure:
models2 = [
    models[0],
    dict(concentration_set='squid axon',
         production_model='wilson', 
         rate_cap_per_pct_mito=0.079, 
         model_name='Squid axon\nmax prod 0.079 mM/s/%'),
    dict(concentration_set='squid axon',
         production_model='wilson', 
         rate_cap_per_pct_mito=0.154, 
         model_name='Squid axon\nmax prod 0.154 mM/s/%'),    
    dict(concentration_set='human skeletal',
         production_model='wilson', 
         rate_cap_per_pct_mito=0.382, 
         model_name='Human skeletal muscle\nmax prod 0.382 mM/s/%'),    
    ]

terminals_ = ['M6Is','M6Ib']

for terminal in terminals_:
    for model in models+models2:
        globals().update(model)
        settings = create_settings(concentration_set, terminal, production_model, 
                                   rate_cap_per_pct_mito=rate_cap_per_pct_mito, 
                                   t_active=10, t_range=[-10,600])
        globals().update(settings)
        fn = data_file('fig6')
        run_simulation(settings, fn, verbose=False)

# %% [code]
for figname,models_ in [ ('fig6',models), ('figS3',models2) ]:

    n_row    = 4
    n_col    = len(models_)
    gridspec = dict(hspace=0.55, wspace=0.55, left=0.08, 
                    right=0.98, bottom=0.07, top=0.95)
    fig,axs  = plt.subplots(n_row, n_col, figsize=(17.6*cm,3.5*n_row*cm), 
                            gridspec_kw=gridspec)

    for i,terminal in enumerate(terminals_):
        for j,model in enumerate(models_):
            globals().update(model)
            globals().update(np.load(data_file('fig6'), allow_pickle=True))
            globals().update(settings.item())
            x1 = -5
            I = (data[0]>=x1+2) & (data[0]<=40)
            T,Rp,ATP = data[[0,2,4]][:,I]
            axs[1,j].plot(T, ATP, **term_opt(terminal))

            # Show production caps as arrow heads at appropriate height along the y axis.
            # If they're too large, have the arrow head point up and write the value.
            style = ArrowStyle('wedge', tail_width=1.5, shrink_factor=0.5)
            opt   = dict(mutation_scale=1, arrowstyle=style, color=colors[terminal])
            dx    = 3.5
            ax    = axs[2,j]
            if rate_cap<0.3:
                ax.add_patch(FancyArrowPatch((x1+dx,rate_cap), (x1,rate_cap), **opt))
            else:
                x1b   = [-2,2][i]
                y,dy  = 0.15,0.025
                ax.add_patch(FancyArrowPatch((x1b,y), (x1b,y+dy), **opt))
                ax.text(x1b+0.5, y+1.1*dy, f'{rate_cap:.2g} mM/s', fontsize=5, rotation=90, 
                        ha='center', va='bottom', color=colors[terminal])
            
            axs[2,j].plot(T, Rp, **term_opt(terminal))
            acc = np.gradient(Rp, T)
            axs[3,j].plot(T, acc, **term_opt(terminal))
            axs[1,j].set_ylim(*ATP[0]*np.array([-0.,1.2]))
        I = times<=40
        axs[0,0].plot(times[I], power_consumed[I], **term_opt(terminal))

    for j in range(1,n_col):
        axs[0,j].axis('off')
    axs_ = axs[0,:2].tolist() + [ax for col in axs.T for ax in col[1:]]
    for i,ax in enumerate(axs_):
        ax.spines.right.set_visible(False)
        ax.spines.top.set_visible(False)
        ax.set_xlabel('Time (s)')
        ax.locator_params(nbins=4)
        ax.text(s=chr(65+i), transform=ax.transAxes, **panel_opt)
        ax.set_xlim(x1,None)
    axs[0,0].set_ylabel('ATP cons. rate (mM/s)')
    axs[0,0].legend(frameon=False) #loc='center left', bbox_to_anchor=(1.6, 0.5))
    for j in range(axs.shape[1]):
        axs[1,j].set_ylabel('[ATP] (mM)')
        axs[2,j].set_ylabel('ATP prod. rate (mM/s)')
        axs[3,j].set_ylabel('ATP prod. accel. (mM/s/s)')
    for i in [2,3]:
        ylim = list(zip(*[ax.get_ylim() for ax in axs[i]]))
        ylim = min((0,)+ylim[0]), max(ylim[1])
        for ax in axs[i]:
            ax.set_ylim(ylim)
    axs[0,0].set_yticks([0,0.4,0.8])

    savefig(figname)
    plt.show()
#     break

# %% [markdown]
### Supplement to figure 6: hydrolysis energy and long-time behavior of production

#### Figure S4: ATP hydrolysis energy

# %% [code]
models_  = models

n_row    = 2
n_col    = len(models_)
gridspec = dict(hspace=0.55, wspace=0.55, left=0.08, 
                right=0.98, bottom=0.28/n_row, top=1-0.2/n_row)
fig,axs  = plt.subplots(n_row, n_col, figsize=(17.6*cm,3.5*n_row*cm), 
                        gridspec_kw=gridspec)
axs      = np.atleast_2d(axs)

for terminal in terminals_:
    for j,model in enumerate(models_):
        globals().update(model)
        globals().update(np.load(data_file('fig6'), allow_pickle=True))
        globals().update(settings.item())
        T,Rc,Rp,ADP,ATP,Ph = data[:,data[0]<=40]
        PhP = Pht - Ph
        Pi  = Pt - ADP - 2*ATP - PhP
        
        # ATP hydrolysis energy.
        ΔG0 = -30.6e3 # ΔG under standard conditions ([ATP]=[ADP]=[P]=1M), in J/mol
        Tmp = 300 # temperature in K
        ΔG  = ΔG0 + R*Tmp*np.log(ADP*Pi/ATP)
        ΔG *= 1e-3 # in kJ/mol
        axs[0,j].plot(T, -ΔG, **term_opt(terminal))
        axs[0,j].set_ylabel('ATP hydrolysis\nenergy (kJ/mol)')

# #         axs[1,j].plot(T, -ΔG*Rp, **term_opt(terminal))
# #         axs[1,j].set_ylabel('Energy production\nrate (kJ/s)')
#         axs[1,j].plot(T, -ΔG*ATP, **term_opt(terminal))
#         axs[1,j].set_ylabel('ATP "energy content" (kJ)')
        
axs_ = axs.flatten()
for i,ax in enumerate(axs_):
    ax.spines.right.set_visible(False)
    ax.spines.top.set_visible(False)
    ax.set_xlabel('Time (s)')
    ax.locator_params(nbins=4)
    ax.text(s=chr(65+i), transform=ax.transAxes, **panel_opt)
#     ax.set_xlim(x1,None)
axs[0,0].legend(frameon=False) #loc='center left', bbox_to_anchor=(1.6, 0.5))
for i in range(axs.shape[0]):
    ylim = list(zip(*[ax.get_ylim() for ax in axs[i]]))
    ylim = min(ylim[0]), max(ylim[1])
    for ax in axs[i]:
        ax.set_ylim(ylim)

savefig('figS4')
plt.show()

# %% [markdown]
#### Figure S2: Total ATP production beyond base rate

For each terminal, every production curve that doesn't crash should have the same area under the curve. The third row of figure 6 made look that unlikely. Here we plot the same curves over a longer period of time.

# %% [code]
models_  = models

n_row    = 1
n_col    = len(models_)
gridspec = dict(hspace=0.55, wspace=0.55, left=0.08, 
                right=0.98, bottom=0.25/n_row, top=1-0.2/n_row)
fig,axs  = plt.subplots(n_row, n_col, figsize=(17.6*cm,4*cm), 
                        gridspec_kw=gridspec)
axs      = np.atleast_2d(axs)

for terminal in terminals_:
    for j,model in enumerate(models_):
        globals().update(model)
        globals().update(np.load(data_file('fig6'), allow_pickle=True))
        globals().update(settings.item())
        T,Rc,Rp,ADP,ATP,Ph = data[:,data[0]<=600]
        PhP = Pht - Ph
        Pi  = Pt - ADP - 2*ATP - PhP
        # Production rate.
        axs[0,j].plot(T, Rp, **term_opt(terminal))
        axs[0,j].set_ylabel('ATP prod. rate (mM/s)')

axs_ = axs.flatten()
for i,ax in enumerate(axs_):
    ax.spines.right.set_visible(False)
    ax.spines.top.set_visible(False)
    ax.set_xlabel('Time (s)')
    ax.locator_params(nbins=4)
    ax.text(s=chr(65+i), transform=ax.transAxes, **panel_opt)
#     ax.set_xlim(x1,None)
axs[0,0].legend(frameon=False) #loc='center left', bbox_to_anchor=(1.6, 0.5))
for i in range(axs.shape[0]):
    ylim = list(zip(*[ax.get_ylim() for ax in axs[i]]))
    ylim = min((0,)+ylim[0]), max(ylim[1])
    for ax in axs[i]:
        ax.set_ylim(ylim)

savefig('figS2')
plt.show()

# %% [markdown]
### Figure 7

# %% [code]
# Use the same models and terminals as figure 6 & S6.

for terminal in terminals_:
    for model in models+models2:
        globals().update(model)
        settings = create_settings(concentration_set, terminal, production_model, 
                                   rate_cap_per_pct_mito=rate_cap_per_pct_mito, 
                                   t_active=120, t_range=[-20,360])
        globals().update(settings)
        fn = data_file('fig7')
        run_simulation(settings, fn, verbose=False)

# %% [code]
models_  = models # +models2

n_row    = 5
n_col    = 2 # len(models_)
gridspec = dict(hspace=0.6, wspace=0.6, left=0.15, 
                right=0.96, bottom=0.03, top=0.96)
fig,axs  = plt.subplots(n_row, n_col, figsize=(8.5*cm,3.5*n_row*cm), 
                        gridspec_kw=gridspec)

times   = np.arange(-20,300)
cycles  = np.array([times[:-1],times[1:]]).T
def cycle_stat(T, X, stat_func=np.nanmean):
    tmax = T[len(X)-1]
    # filter: tmax>t1 includes the last unfinished cycle in case of crash. tmax>t2 doesn't.
    TX = [ ( (t1+t2)/2, stat_func(X[(T>=t1)&(T<t2)]) ) for t1,t2 in cycles if tmax>t2 ]
    return np.array(TX).T

for j,terminal in enumerate(terminals_[::-1]):
    shades = create_color_shades(colors[terminal])
    for i,model in enumerate(models_):
        globals().update(model)
        globals().update(np.load(data_file('fig7'), allow_pickle=True))
        globals().update(settings.item())
        T,Rc,Rp,ADP,ATP,Ph = data[:,data[0]<=300]
        # Plot [ATP].
        axs[1,j].plot(T, ATP/ATP[0], color=shades[i], label=model_name)
        # Plot production rate.
        axs[2,j].plot(T, Rp, color=shades[i], label=model_name)
        # Show production caps.
        style = ArrowStyle('wedge', tail_width=1.5, shrink_factor=0.5)
        opt   = dict(mutation_scale=1, arrowstyle=style, color=shades[i], clip_on=False)
        dx    = 25
#         x     = [300, 312, 324, 324][i] + 5
        x     = [0, 12, 0, 24][i] - 50
        axs[2,j].add_patch(FancyArrowPatch((x+dx,rate_cap), (x,rate_cap), **opt))
        # Plot production acceleration.
        acc = np.gradient(Rp,T)
        axs[3,j].plot(T, acc, color=shades[i], label=model_name)
    # Plot cycle-mean of power consumption, plus min and max as a shaded area.
    T,Cavg = cycle_stat(times, power_consumed, np.mean)
    T,Cmin = cycle_stat(times, power_consumed, np.min)
    T,Cmax = cycle_stat(times, power_consumed, np.max)
    w = 0.1 # equivalent opacity (eps output doesn't allow transparency)
    c = [w*x+1-w for x in to_rgb(colors[terminal])]
    axs[0,j].fill_between(T, Cmin, Cmax, color=c, label=model_name)
    axs[0,j].plot(T, Cavg, color=colors[terminal], label=model_name)

for ax in axs[-1]:
    ax.axis('off')
for i,ax in enumerate(axs[:-1].flatten()):
    ax.spines.right.set_visible(False)
    ax.spines.top.set_visible(False)
    ax.set_xlabel('Time (s)')
#     ax.set_xlim(-25,300)
    ax.set_xlim(-50,300)
    ax.locator_params(nbins=4)
    ax.text(s=chr(65+i), transform=ax.transAxes, **panel_opt)
for i in range(2):
    axs[3,i].legend(loc='upper center', bbox_to_anchor=(0.3, -0.55), labelspacing=0.7)
for j in range(axs.shape[1]):
#     axs[0,j].set_ylabel('ATP cons. rate: cycle mean\nand envelope (mM/s)', labelpad=2)
    axs[0,j].set_ylabel('ATP cons. rate (mM/s)')
    axs[1,j].set_ylabel('[ATP] / [ATP]$_{rest}$')
    axs[1,j].set_ylim(0,1.1)
    axs[2,j].set_ylabel('ATP prod. rate (mM/s)')
    axs[3,j].set_ylabel('ATP prod. accel. (mM/s/s)')
for row in axs[:-1]:
    ylim = list(zip(*[ax.get_ylim() for ax in row]))
    ylim = min((0,)+ylim[0]), max(ylim[1])
    for ax in row:
        ax.set_ylim(ylim)

savefig('fig7')
plt.show()

# %% [code]
```
