## Supplementary File S1 for "Mitochondrial phosphagen kinases support the volatile power demands of motor nerve terminals": process-terminal-data.ipynb

```
# %% [code]
import numpy as np
from scipy.constants import Avogadro
import pandas as pd
from openpyxl.styles import Alignment

# %% [markdown]
### Load data from Justs2022 supplementary table.

# %% [code]
# filename  = 'Justs et al 2022 - Extended Data Set - c_table.xlsx'
filename  = 'Justs et al 2022 - Extended Data Set - symbols replaced.xlsx'
terminals = ['MN6-Ib', 'MN6-Is', 'MN13-Ib', 'MN13-Is', 'MN12-Ib', 'MN12-Is']

df  = pd.read_excel(filename, sheet_name='Table 1-1', header=5, index_col=[1,2]).T
df1 = df.loc[terminals, ['Volume','Surface Area']]

df  = pd.read_excel(filename, sheet_name='Table 2-1', header=4, index_col=[1,2]).T
df2 = df.loc[terminals, ['Endogenous Firing Rate (EFR)', 'Duty Cycle', 
                         'ATP Spent on Glutamate / AP']]
df2 = df2.dropna(axis=1)

df  = pd.read_excel(filename, sheet_name='Table 3-1', header=5, index_col=[1,2]).T
df3 = df.loc[terminals, ['ATP Spent on Ca2+ Extrusion / AP']]

df  = pd.read_excel(filename, sheet_name='Table 4-1', header=5, index_col=[1,2]).T
df4 = df.loc[terminals, ['ATP Spent on Na+ / AP']]
df4 = df4.dropna(axis=1).rename(columns={'molecules':'molecules 106'})

df  = pd.read_excel(filename, sheet_name='Table 5-1', header=5, index_col=[1,2]).T
df5 = df.loc[terminals, ['Non-Signaling']]
df5 = df5.rename(columns={'106 molecules':'106 molecules/sec', 
                          'Non-Signaling':'Non-Signaling Power Demand (ATP)'})

df  = pd.read_excel(filename, sheet_name='Table 6-1', header=5, index_col=[1,2]).T
df6 = df.loc[terminals, ['Mitochondrial Density']]
df6 = df6.dropna(axis=1)

data = pd.concat([df1, df2, df3, df4, df5, df6], axis=1)
data = data.rename(columns={'micron<3>':'μm^3', 'micron<2>':'μm^2', 'Seconds':'s', 
                            'molecules 106':'10^6 molecules', 
                            '106 molecules/sec':'10^6 molecules/s'})
# data.index.name = 'Terminal'
data.columns.names = ['Quantity', 'Unit']
display(data)

# Strip error values and convert to float.
data = data.astype(str).applymap(lambda x: x.split('±')[0]).astype(float)
# display(data)

data0 = data.copy()

# %% [markdown]
### Convert to SI

# %% [code]
data = data0.copy()
data['Volume'] *= 1e-18
data['Surface Area'] *= 1e-12
for k in ['Glutamate', 'Ca2+ Extrusion', 'Na+']:
    data[f'ATP Spent on {k} / AP'] *= 1e6 #/Avogadro
data['Non-Signaling Power Demand (ATP)'] *= 1e6 #/Avogadro
data = data.rename(columns={'μm^3':'m^3', 'μm^2':'m^2', 
                            '10^6 molecules':'molecules', 
                            '10^6 molecules/sec':'molecules/s'})
data

# %% [markdown]
### Add variable names for simulations

# %% [code]
data = data.T
data['Variable Name'] = ['volume', 'area', 'firing_rate', 'duty_cycle', 'committed_nt', 
                         'committed_ca', 'committed_na', 'base_rate', 'mito_density']
data = data.reset_index().set_index(['Quantity','Unit','Variable Name']).T
data

# %% [markdown]
### More variables for simulations

# %% [code]
extra_variables = [
    ('Cycle duration', 's', 'full_cycle', 1), 
    ('Ca2+ Decay Time', 's', 'tau_ca', 0.0389), 
    ('Na+ Decay Time', 's', 'tau_na', 63.5), 
    ('Glutamate Decay Time 1', 's', 'tau_nt1', 0.175), 
    ('Glutamate Decay Time 2', 's', 'tau_nt2', 7.21), 
    ('Glutamate Decay Time 3', 's', 'tau_nt3', 20),
    ('Relative ATP Cost of Glutamate process 1', '% of Total Glutamate Cost', 'alpha_nt1', 0.13), 
    ('Relative ATP Cost of Glutamate process 2', '% of Total Glutamate Cost', 'alpha_nt2', 0.34), 
    ('Relative ATP Cost of Glutamate process 3', '% of Total Glutamate Cost', 'alpha_nt3', 0.53),
    ]
extra_variables = pd.DataFrame(extra_variables, columns=['Quantity', 'Unit', 'Variable Name', 'Value'])
extra_variables = extra_variables.set_index('Quantity')
extra_variables

# %% [markdown]
### Save as an excel spreadsheet

# %% [code]
writer = pd.ExcelWriter('terminal-data.xlsx', engine='openpyxl')
data0.to_excel(writer, sheet_name='Original Units')
data.to_excel(writer, sheet_name='SI Units')
extra_variables.to_excel(writer, sheet_name='Extra Variables')
for k,sheet in writer.sheets.items():
    for col in sheet.columns:
        sheet.column_dimensions[col[0].column_letter].width = 15
        for cell in col:
            cell.alignment = Alignment(wrap_text=True, horizontal='center')
writer.save()

# %% [markdown]
### Save as a simulation-ready numpy file

Loading the variables in this pickle file is essentially equivalent to loading the old terminal_data module.

# %% [code]
terminal_data = {}
for terminal in terminals:
    term = terminal.replace('N','').replace('-','')
    df = data.T.reset_index().set_index('Variable Name')
    D  = df[terminal].to_dict()
    D.update(extra_variables.set_index('Variable Name')['Value'].to_dict())
    globals().update(D)
    
    D['firing_period']     = 1/D['firing_rate']
    D['committed_amounts'] = np.array([committed_ca, committed_na,
                                       committed_nt*alpha_nt1,
                                       committed_nt*alpha_nt2,
                                       committed_nt*alpha_nt3])
    D['committed_decays']  = np.array([tau_ca, tau_na, tau_nt1, tau_nt2, tau_nt3])
    D['avg_active_rate']   = (committed_ca+committed_na+committed_nt) \
                              *(int(duty_cycle*firing_rate)+1)/full_cycle
    D['avg_total_rate']    = D['avg_active_rate'] + base_rate
    
#     terminal_data[term]    = D
    terminal_data[term]    = { k:v for k,v in D.items() if k[:4] not in ['tau_','alph'] }
#     break

np.savez('terminal-data.npz', **terminal_data)
# display(terminal_data['M6Ib'])

# %% [markdown]
### Demo reading the npz file

# %% [code]
terminal_data = np.load('terminal-data.npz', allow_pickle=True)
terminal_data = { k:v.item() for k,v in terminal_data.items() }
terminal_data['M6Ib']

# %% [code]
```
