## Supplementary File S1 for "Mitochondrial phosphagen kinases support the volatile power demands of motor nerve terminals": README.txt

This folder contains the scripts needed to perform the numerical simulations and create initial versions of the simulation figures discussed in the manuscript "Mitochondrial phosphagen kinases support the volatile power demands of motor nerve terminals".

Files with the extension ".ipynb" are python jupyter notebooks.

"terminal-data.xlsx" contains the motor neuron terminal data from (Justs et al. 2022).

"process-terminal-data.ipynb" extracts the information contained in "terminal-data.xlsx" and saves it in "terminal-data.npz". The latter is used by "perform-simulations.ipynb" and "volatility-methods-figs.ipynb".

"perform-simulations.ipynb" performs all numerical simulations discussed in the manuscript and creates initial versions of figures 3, 6, 7, S2, S3, and S4. The output of the simulations, about 160MB, is saved in "figure-data/". The figures are saved in "figures/".

"volatility-methods-figs.ipynb" creates initial versions of figures S8 and S9. The figures are saved in "figures/".
