## Supplementary File S1 for "Mitochondrial phosphagen kinases support the volatile power demands of motor nerve terminals": setup_simulation.py

```
import numpy as np
import scipy as sp
from scipy.constants import Avogadro
from numba import jit, njit, cfunc, objmode
from NumbaMinpack import hybrd, lmdif, minpack_sig

#===============================================================================

# Settings from Wilson's 2017 paper.
wilson_settings = dict(
    Ke   = 6.4e11,   # NAD to cyt. c equil constant 
    k1   = 3.5e9,    # a 33+ -Cu 2+ + c 2+ ---> a 33+ - Cu 1+ + c 3+
    k1r  = 3.5e7,    # a 33+ - Cu 1+ + c 3+ ---> a 33+ - Cu 2+ + c 2+
    k2   = 3.5e8,    # a 33+ - Cu 1+ + O 2 ---> a 33+ - Cu 1+ - O 2
    k2r  = 1e1,      # a 33+ -Cu1+- O 2 ---> a 33+ -Cu1+ + O 2
    K3   = 2e6,      # a 33+ -Cu 1+ -O 2 + H + <---> O 2 -a 32+ -Cu 2+
    k4a  = 1.2e8,    # a 33+ -Cu 1+ -O 2 + c 2+ ----> a 33+ -O 2 -Cu 2+
    k4b  = 4e7,      # O 2 -a 32+ -Cu 2+ + c 2+ ----> a 33+ -O 22- -Cu 2+
    K5   = 1e25,     # a 33+ - O 22- -Cu 2+ + 2H + + 2 c 2+ <-> a 33+ - Cu 2+ + 2 c 3+ +H 2 O
    a3t  = 1e-6,     # cyt. oxidase (a, 2 Cu, a 3 ) conc.
    ct   = 2e-6,     # cyt. c conc. (2 uM)
    Emc  = 0.235,    # cyt. c half rededuction potential (Em in Volts)
    NAD  = 0.1,      # [NAD+]/[NADH]
    W    = 7.1,      # pH
    O    = 80e-6,    # O2 concentration, in M.
    )

indent = '     '

def setup(settings):
    globals().update(wilson_settings)
    globals().update(settings)

    # ---------- Equilibration ----------

    if (Kamp is None) and (Kph is None):
        @njit
        def enforce_equilibrium(AMP, ADP, Ph):
            return AMP,ADP,Ph,0,1
    if (Kamp is None) and (not Kph is None):
        @cfunc(minpack_sig)
        def d_eq(xvec, fvec, args):
            AMP  = args[0]
            ADP  = args[1]
            Ph   = args[2]
            At   = args[3]
            Pht  = args[4]
            Kph  = args[5]
            x    = xvec[0] # x = progress of PhP+ADP->Ph+ATP
            ADP_ = ADP - x
            Ph_  = Ph  + x
            fvec[0] = ( (At-AMP-ADP_) * Ph_ )/( Kph * ADP_ * (Pht-Ph_) ) - 1
        d_eq_ptr = d_eq.address
        @njit
        def enforce_equilibrium(AMP, ADP, Ph, At=At, Pht=Pht, Kamp=Kamp, Kph=Kph, xtol=1e-8):
            args = np.array([AMP,ADP,Ph,At,Pht,Kph])
            x0   = np.array([0.])
            sol  = hybrd(d_eq_ptr, x0, args, xtol)
            x    = sol[0][0]
            err  = abs(sol[1][0])
            return AMP,ADP-x,Ph+x,err,sol[3]
    if (not Kamp is None) and (Kph is None):
        @cfunc(minpack_sig)
        def d_eq(xvec, fvec, args):
            AMP  = args[0]
            ADP  = args[1]
            Ph   = args[2]
            At   = args[3]
            Pht  = args[4]
            Kamp = args[5]
            x    = xvec[0] # x = progress of AMP+ATP->2ADP
            AMP_ = AMP -   x
            ADP_ = ADP + 2*x
            ATP_ = At - AMP_ - ADP_
            fvec[0] = ADP_**2 / ( Kamp * AMP_ * ATP_ ) - 1
        d_eq_ptr = d_eq.address
        @njit
        def enforce_equilibrium(AMP, ADP, Ph, At=At, Pht=Pht, Kamp=Kamp, Kph=Kph, xtol=1e-8):
            args = np.array([AMP,ADP,Ph,At,Pht,Kamp])
            x0   = np.array([0.])
            sol  = hybrd(d_eq_ptr, x0, args, xtol)
            x    = sol[0][0]
            err  = abs(sol[1][0])
            return AMP-x,ADP+2*x,Ph,err,sol[3]
    if (not Kamp is None) and (not Kph is None):
        @cfunc(minpack_sig)
        def d_eq(xvec, fvec, args):
            AMP  = args[0]
            ADP  = args[1]
            Ph   = args[2]
            At   = args[3]
            Pht  = args[4]
            Kamp = args[5]
            Kph  = args[6]
            AMP_ = AMP -   xvec[0]           # xvec[0] = progress of AMP+ATP->2ADP
            ADP_ = ADP + 2*xvec[0] - xvec[1] # xvec[1] = progress of PhP+ADP->Ph+ATP
            ATP_ = At - AMP_ - ADP_
            Ph_  = Ph  + xvec[1]
            fvec[0] = ADP_**2 / ( Kamp * AMP_ * ATP_ ) - 1
            fvec[1] = ( ATP_ * Ph_ )/( Kph * ADP_ * (Pht-Ph_) ) - 1
        d_eq_ptr = d_eq.address
        @njit
        def enforce_equilibrium(AMP, ADP, Ph, At=At, Pht=Pht, Kamp=Kamp, Kph=Kph, xtol=1e-8):
            args = np.array([AMP,ADP,Ph,At,Pht,Kamp,Kph])
            x0   = np.array([0.,0.])
            sol  = hybrd(d_eq_ptr, x0, args, xtol)
            x    = sol[0]
            err  = np.sqrt(sol[1][0]**2+sol[1][1]**2)
            return AMP-x[0],ADP+2*x[0]-x[1],Ph+x[1],err,sol[3]

    # ---------- Consumption ----------

    # Times at which action potentials are fired.
    ftimes = np.array([ tf1+tf2 for tf1 in np.arange(0,min(t2,t_active),full_cycle) \
                                for tf2 in np.arange(0,duty_cycle,firing_period) \
                                if tf1+tf2<min(t2,t_active)])
    # Power consumption at time t.
    @njit
    def consumption(t):
        C   = base_rate # rest consumption
        for tf in ftimes:
            if tf<t:
                for a,tau in zip(committed_amounts,committed_decays):
                    C += a / tau * np.exp((tf-t)/tau)
        return C/(Avogadro*volume)
    
    # ---------- Production ----------

    if production_model=='rest':
        @njit
        def production_rate_(AMP, ADP, Ph):
            return base_rate/(Avogadro*volume)
    elif production_model=='wilson':
        @njit
        def production_rate__(AMP, ADP, Ph, At=1e-3*At, Pht=1e-3*Pht, Pt=1e-3*Pt):
            # First convert mM to M.
            AMP,ADP,Ph = 1e-3*AMP,1e-3*ADP,1e-3*Ph
            H   = 10**-W        # hydrogen ion conc., in M. 
            ATP = At-AMP-ADP        # [ATP].
            PhP = Pht - Ph          # [Ph]
            Pi  = Pt - ADP - 2*ATP - PhP    # Free (inorganic) phosphate
            Q   = ( 7.6 + 1.42*np.log10(ATP/(ADP*Pi)) )/46.183 # Gibbs energy (volts)
            z   = 10**(Q/0.059)     # effect of energy coupling.
            kf1 = k1/np.sqrt(z)     # couples k1 to energy state
            kr1 = k1r*np.sqrt(z)    # couples k1r to energy state
            D   = NAD**0.5 * z**2 * (H/Ke)**0.5 # uses energy state and NAD+/NADH to calc co
            co  = D*ct/(1+D)        # co is conc. oxidized cyt c
            cr  = ct - co           # cr is conc red cyt c
            A   = (k2r+k4a*cr+k4b*cr*K3*H)/(k2*O)       # variable A
            B   = (k2*O*A+(kr1*co)*A-k2r)/((kf1*cr))    # variable B
            C   = 1/(K5*H**2) * ((co/cr)**2 * z**2) * B # variable C
            III = a3t/(1 + K3*H + A + B + C)            # intermediate [III]
            return (k4a*cr+k4b*cr*K3*H) * III*4/ct      # cyt TN s -1
        # Rescale to match consumption at rest.
        production_scale = base_rate/(Avogadro*volume) / production_rate__(AMP, ADP, Ph)
        @njit
        def production_rate_(AMP, ADP, Ph):
            return production_scale * production_rate__(AMP, ADP, Ph)
    else:
        raise Exception('Unknown production model.')
    if rate_cap is None:
        production_rate = production_rate_
    else:
        @njit
        def production_rate(AMP, ADP, Ph, r0=rate_cap, n=rate_cap_exponent):
            r = production_rate_(AMP, ADP, Ph)
            return ( 1/r**n + 1/r0**n )**(-1/n)
    
    # ---------- Integration ----------

    # Times at which to predict/print concentrations.
    times = np.arange(t1,t2,dt) # Constant time step (for comparison with previous version).
    if {'dt1','dt2','dt3'}.issubset(settings.keys()):
        # Use a smaller time step dt2 near consumption peaks.
        times = np.array([t1])
        for tf in ftimes:
            if t1<tf<t2:
                times = np.append(times, np.arange(times[-1], tf, dt1)[1:])
                times = np.append(times, [tf-dt2])
                times = np.append(times, tf + np.arange(0, dt3, dt2))
        times = np.append(times, np.arange(times[-1], t2, dt1)[1:])
    
    # Power consumption at each simulation time.
    @njit
    def power_consumed_(times=times):
        return np.array([ consumption(t) for t in times ])
    power_consumed = power_consumed_()
    
    # Forward Euler method.
    @njit
    def step_forward(t, dt, AMP, ADP, Ph, rp_min=None, rp_max=None, rate_cap=rate_cap):
        c    = consumption(t)*dt
        rp   = production_rate(AMP, ADP, Ph)
        if not rp_min is None:
            rp = max(rp,rp_min)
        if not rp_max is None:
            rp = min(rp,rp_max)
        ADP  = ADP + c - rp*dt
        AMP,ADP,Ph,e,m = enforce_equilibrium(AMP, ADP, Ph)
        return c/dt,rp,AMP,ADP,Ph,e,m

    # Integrate with fixed time step dt but only save results at ptimes.
    @njit
    def integrate(AMP=AMP, ADP=ADP, Ph=Ph, acceleration_cap=acceleration_cap, verbose=False):
        output = []
        t      = times[0]
        rp_,t_ = None,t   # Last t and rp; used to cap the acceleration.
        for tp in times[1:]:
            first = True
            exit  = False
            while t<tp:
                dt_ = min(dt,tp-t)
                if (not acceleration_cap is None) and (not rp_ is None):
                    drp = (t-t_)*acceleration_cap
                    rc,rp,AMP_,ADP_,Ph_,err,msg = step_forward(t,dt_,AMP,ADP,Ph,rp_-drp,rp_+drp)
                    rp_,t_ = rp,t
                else:
                    rc,rp,AMP_,ADP_,Ph_,err,msg = step_forward(t,dt_,AMP,ADP,Ph)
                    rp_ = rp
                if first:
                    output.append([t,rc,rp,ADP,At-AMP-ADP,Ph])
                    first = False
                AMP,ADP,Ph = AMP_,ADP_,Ph_
                t += dt_
                if verbose and err>1e-6:
                    print(indent+'[Warning] Failed to equilibrate at t =', t, 'Error = ', err, 'Error code =', msg)
                if ( (not Kamp is None) and (AMP<=0) ) \
                        or AMP>=At or ADP<=0 or ADP>=At \
                        or ( not Kph is None and (Ph<=0 or Ph>=Pht) ) \
                        or np.isnan(AMP) or np.isnan(ADP) or np.isnan(Ph):
                    if verbose:
                        print(indent+'[Warning] Concentration out of bounds at t =',t)
                    exit = True
                    break
            if exit:
                break
        return np.array(output)

    return locals()

#===============================================================================

concentration_sets = {
    'squid axon no arginine': 
        dict(ATP=2.16, ADP=0.02, Pi=17.8, Kph=None, Tmp=288.15),
    'squid axon': 
        dict(ATP=2.16, PhP=7.5, Ph=3.33, Pi=17.8, Kph=39.64, Tmp=288.15),
    'human grey matter': 
        dict(ATP=3, PhP=3, Ph=3.4, Pi=0.93, Kph=166, Tmp=310.35),
    'human skeletal': 
        dict(ATP=6, PhP=18.9, Ph=12.3, Pi=8.48, Kph=166, Tmp=309),
}

for k in concentration_sets:
    globals().update(concentration_sets[k])
    if Kph is None:
        concentration_sets[k].update(Ph=0, PhP=0)
    if not 'ADP' in concentration_sets[k]:
        ADP = ATP*Ph/(Kph*PhP)
    Kamp = 1
    AMP  = ADP**2/(Kamp*ATP)
    concentration_sets[k].update(ADP=ADP, AMP=AMP, Kamp=Kamp)

terminal_data = {}
def load_terminal_data(path):
    global terminal_data
    terminal_data = np.load(path, allow_pickle=True)
    terminal_data = { k:v.item() for k,v in terminal_data.items() }
    return terminal_data


def create_settings(concentration_set, terminal, production_model, 
                    t_active, t_range, dt=1e-4, acceleration_cap=None, 
                    rate_cap_per_pct_mito=None, rate_cap_exponent=10 ):
    
    D = locals().copy()
    D.update(terminal_data[terminal])
    D.update(wilson_settings)
    D.update(concentration_sets[concentration_set])
    globals().update(D)
    
    At  = ATP + ADP + AMP
    Pht = Ph + PhP
    Pt  = Pi + ADP + 2*ATP + PhP
    D.update(At=At, Pht=Pht, Pt=Pt)
    
    # Production settings.
    rate_cap = None if rate_cap_per_pct_mito is None else rate_cap_per_pct_mito*mito_density
    D.update(rate_cap=rate_cap)
    
    # Integration settings.
    t1,t2 = t_range        # start and end time of the simulation 
    dt    = dt # 1e-3 # 2e-5           # t_start, t_end, time step
    dt1   = 5e-3           # large printing period (away from peaks)
    dt2   = dt             # smaller printing period (around peaks)
    dt3   = 2e-3           # how long to use dt2 after each peak
    D.update(t1=t1, t2=t2, dt=dt, dt1=dt1, dt2=dt2, dt3=dt3)
    
    return D
```
