## Supplementary File S1 for "Mitochondrial phosphagen kinases support the volatile power demands of motor nerve terminals": volatility-methods-figs.ipynb

```
# %% [code]
import numpy as np
import scipy as sp
from scipy.optimize import fsolve
import matplotlib.pyplot as plt
from matplotlib.colors import hsv_to_rgb, rgb_to_hsv, to_rgb
from matplotlib.patches import FancyArrowPatch, ArrowStyle
plt.rcParams['figure.facecolor'] = 'w'

from setup_simulation import *
terminal_data = load_terminal_data('terminal-data.npz')
terminals     = sorted(terminal_data.keys())[::-1]
terminals     = terminals[1::2]+terminals[::2]
print(terminals)

# %% [markdown]
# Consumption

# %% [code]
# for term in terminal_data.keys():

dt = 0.0001
term = 'M6Ib'
print(*terminal_data[term].keys(), sep=', ')
globals().update(terminal_data[term])

firing_period = 0.1
duty_cycle    = 0.61

τ = 0.2
N = τ

def single_process_consumption(t, i_process=0, t_AP=0):
    return (N/τ)*np.heaviside(t,1)*np.exp(-(t-t_AP)/τ) 

gskw = dict(wspace=0.3)
fig,axs = plt.subplots(1, 2, figsize=(8,3), gridspec_kw=gskw)

ax   = axs[0]
time = np.arange(-0.2,1,dt)
time2 = np.arange(-0.2,2,dt)
train_time = np.zeros(np.shape(time))
cons = single_process_consumption(time)
ax.plot(time, cons)
ax.set_xlim(-0.2,1)
ax.fill_between(time, cons, step="pre", alpha=0.3, color='lightblue')

t_max  = 2
ftimes = np.array([ tf1+tf2 for tf1 in np.arange(0,t_max,full_cycle) \
                            for tf2 in np.arange(0,duty_cycle,firing_period) \
                            if tf1+tf2<t_max])
#cons_train = single_process_consumption(time) + single_process_consumption(time - 0.1) + \
#                single_process_consumption(time - 0.2) + single_process_consumption(time - 0.3) + \
#                single_process_consumption(time - 0.4) + single_process_consumption(time - 0.5)
#train_time[ftimes+0.2] = np.ones(len(ftimes))


kwarg1 = dict(shape='full', length_includes_head=True, head_length=0.0, head_width=0.04 )

ax.arrow(-0.04, 0, 0, N/τ, color='green', **kwarg1)
ax.arrow(0, single_process_consumption(τ), τ, 0, color='purple', **kwarg1)

#ax.annotate()
ax.annotate(r'$\dfrac{N}{τ}$', xy =(-0.15, N), xytext =(-0.12, N/τ*0.8), color='green')
ax.annotate('N', xy =(-0.15, 5), xytext =(0.21, N/τ*0.1), alpha=0.7,color='blue')
ax.annotate('τ', xy =(-0.15, 1), xytext =(0.07, N/τ*0.4), color='purple')

for ax in axs:
    ax.spines['top'].set_visible(False)
    ax.spines['right'].set_visible(False)

ax = axs[1]
ax.spines['left'].set_visible(False)
for t in ftimes:
    ax.plot([t,t], [0,1], 'C0')
ax.set_xlim(-0.2,2)
ax.set_ylim(0,2)

kwarg = dict(length_includes_head=True, head_length=0.00, head_width=0.07,
             shape='full', lw=1)
x,xpad,y0,ypad = 0,0.1,1.15,0.2
for i,(s,dx,c) in enumerate([ (r'$(EFR)^{-1}$', firing_period, 'r'), 
                             ('Duty Cycle', duty_cycle, 'orange'), 
                             ('Full Cycle', full_cycle, 'brown') ]):
    y = y0 + i*ypad
    ax.arrow(x, y, dx, 0, color=c, **kwarg)
    ax.arrow(x+dx, y, -dx, 0, color=c, **kwarg)
    ax.text(x+dx+xpad, y, s, ha='left', va='center', color=c)
#     ax.text(full_cycle+xpad, y, s, ha='left', va='center', color=c)

for ax in axs:
    ax.set_xlabel('Time (au)')
    ax.locator_params(nbins=2)
axs[0].set_ylabel('ATP Consumption Rate')
kwarg = dict(fontsize=10, va='center', y=1.08)
axs[0].set_title('Time Course of Energy Demand\nfor One Signaling Process', **kwarg)
axs[1].set_title('Parameters of Action Potential Bursts', **kwarg)

axs[1].set_yticks([])
axs[1].set_ylabel('Action potentials')

plt.tight_layout()
plt.savefig('figures/figS8.png', dpi=300)
plt.show()

# %% [markdown]
# ATP production curve

# %% [code]
models = [
    dict(concentration_set='squid axon no arginine',
         production_model='wilson', 
         rate_cap_per_pct_mito=0.032, 
         model_name='Squid axon, no arginine\nmax prod 0.032 mM/s/%'),
    dict(concentration_set='squid axon',
         production_model='wilson', 
         rate_cap_per_pct_mito=0.032, 
         model_name='Squid axon\nmax prod 0.032 mM/s/%'),
#     dict(concentration_set='dog brain',
#          production_model='wilson', 
#          rate_cap_per_pct_mito=0.025, 
#          model_name='Dog brain concentrations\nmax prod=0.025 mM/s/%\nw/ creatine'),
    dict(concentration_set='human grey matter',
         production_model='wilson', 
         rate_cap_per_pct_mito=0.025, 
         model_name='Human grey matter\nmax prod 0.025 mM/s/%'),
    dict(concentration_set='human skeletal',
         production_model='wilson', 
         rate_cap_per_pct_mito=0.032, 
         model_name='Human skeletal muscle\nmax prod 0.032 mM/s/%')
    ]

# Extra models for supplementary figure:
models2 = [
    models[0],
    dict(concentration_set='squid axon',
         production_model='wilson', 
         rate_cap_per_pct_mito=0.079, 
         model_name='Squid axon\nmax prod 0.079 mM/s/%'),
    dict(concentration_set='squid axon',
         production_model='wilson', 
         rate_cap_per_pct_mito=0.154, 
         model_name='Squid axon\nmax prod 0.154 mM/s/%'),    
    dict(concentration_set='human skeletal',
         production_model='wilson', 
         rate_cap_per_pct_mito=0.382, 
         model_name='Human skeletal muscle\nmax prod 0.382 mM/s/%'),    
    ]

# %% [code]
# gs     = dict(bottom=0.2, top=0.95, left=0.15, right=0.7)
# fig,ax = plt.subplots(1, 1, figsize=(4.5,3), gridspec_kw=gs)
fig = plt.figure(figsize=(4.5,4))

terminal = terminals[0]
print(terminal)
for i,model in enumerate([models[1], models2[-1]]):
# for i,model in enumerate(models+models2):
    color = f'C{i}'

    globals().update(model)
    settings = create_settings(concentration_set, terminal, production_model, 
                               rate_cap_per_pct_mito=rate_cap_per_pct_mito, 
                               t_active=10, t_range=[-10,600])
    globals().update(settings)

    def get_concentrations(atp, At=At, Pht=Pht, Kamp=Kamp, Kph=Kph):
        f   = lambda adp: Kamp*(At-adp-atp)*atp - adp**2
        adp = fsolve(f, x0=At-atp)[0]
        amp = At-adp-atp
        m   = (atp/adp)/Kph
        ph  = Pht/(m+1)
        r   = production_rate if capped else production_rate_
        return amp,adp,atp,ph,r(amp, adp, ph)
    
    def get_energy_state(AMP, ADP, Ph, At=At, Pht=Pht, Pt=Pt):
        ATP = At-AMP-ADP        # [ATP].
        PhP = Pht - Ph          # [Ph]
        Pi  = Pt - ADP - 2*ATP - PhP    # Free (inorganic) phosphate
        return ATP/(ADP*Pi)

    # es=energy state=[ATP]/([ADP][P]), in 1/mM
    def production_rate__(es):
        es_ = 1e3*es # Convert from 1/mM to 1/M
        H   = 10**-W        # hydrogen ion conc., in M. 
        Q   = ( 7.6 + 1.42*np.log10(es_) )/46.183 # Gibbs energy (volts)
        z   = 10**(Q/0.059)     # effect of energy coupling.
        kf1 = k1/np.sqrt(z)     # couples k1 to energy state
        kr1 = k1r*np.sqrt(z)    # couples k1r to energy state
        D   = NAD**0.5 * z**2 * (H/Ke)**0.5 # uses energy state and NAD+/NADH to calc co
        co  = D*ct/(1+D)        # co is conc. oxidized cyt c
        cr  = ct - co           # cr is conc red cyt c
        A   = (k2r+k4a*cr+k4b*cr*K3*H)/(k2*O)       # variable A
        B   = (k2*O*A+(kr1*co)*A-k2r)/((kf1*cr))    # variable B
        C   = 1/(K5*H**2) * ((co/cr)**2 * z**2) * B # variable C
        III = a3t/(1 + K3*H + A + B + C)            # intermediate [III]
        return (k4a*cr+k4b*cr*K3*H) * III*4/ct      # cyt TN s -1

    # Rescale to match consumption at rest.
    es0 = get_energy_state(AMP, ADP, Ph)
    production_scale = base_rate/(Avogadro*volume) / production_rate__(es0)
    def production_rate_(es):
        return production_scale * production_rate__(es)

    production_rate_ = np.vectorize(production_rate_)

    if rate_cap is None:
        production_rate = production_rate_
    else:
        def production_rate(es, r0=rate_cap, n=rate_cap_exponent):
            r = production_rate_(es)
            return ( 1/r**n + 1/r0**n )**(-1/n)

    ES = 1/np.geomspace(1e-2,1e5,1000)
    R  = production_rate(ES)
    R_ = production_rate_(ES)
    name = model_name.split('\n')[0]
    opt1 = dict(lw=5, alpha=0.3, color=color)
    opt2 = dict(lw=2, alpha=1, color=color)
#     opt1 = dict(lw=2, alpha=1, color=color, zorder=5)
#     opt2 = dict(lw=5, alpha=0.5, color=color)
    plt.plot(1/ES, R, label=name+'\n'+r'$p_{max}=$'+f'{rate_cap:.2g}mM/s', **opt1)
    plt.plot(1/ES, R_, dashes=(3,2), **opt2)
    plt.axhline(rate_cap, dashes=(4,2,1,2), **opt2)
    plt.scatter([1/es0], [base_rate/(Avogadro*volume)], s=200, color=color)

plt.xscale('log')
plt.yscale('log')
plt.xlim(1e-2,1e4)
plt.ylim(3e-3,None)
plt.minorticks_off()
# plt.xlabel('Inverse Energy State')
plt.xlabel(r'Inverse Energy State $\dfrac{[ADP][P_i]}{[ATP]}$  (in mM)')
plt.ylabel('ATP Production Rate  (in mM/s)')
plt.legend(loc='lower right')
plt.gca().spines['top'].set_visible(False)
plt.gca().spines['right'].set_visible(False)

plt.tight_layout()
plt.savefig('figures/figS9.png', dpi=300)
plt.show()

# %% [code]
```
